# Glucose is a ligand-like trigger for secretion of postprandial GLP-1

**DOI:** 10.64898/2026.01.04.697549

**Authors:** Chun Li, Mengqi Li, Heng-Ye Yang, Xiao Tian, Chen-Song Zhang, Lin Zhao, Jianfeng Wu, Cixiong Zhang, Shiqi Liu, Xiaotong Yang, Xiufeng Sun, Junjie Chen, Baoding Zhang, Chuyu Yun, Jun Lin, Wenxuan Ma, Teng Ma, Qingchao Zhu, Zhong-Zheng Zheng, Xuehui Hong, Xianming Deng, Kai Wang, Changtao Jiang, Meiqin Hu, Xiaoying Li, Haoxing Xu, Sheng-Cai Lin

## Abstract

Glucagon-like peptide 1 (GLP-1) is secreted from the intestinal L cells in a glucose-dependent manner. However, the mechanism underlying how glucose stimulates the secretion remains unclear. Here, we found that glucose at postprandial concentrations near the *K*_m_ of GK for glucose (∼10 mM), as well as nonmetabolizable glucose analogs, triggers GLP-1 secretion by intestinal L cells through directly binding to glucokinase (GK), without increasing intracellular ATP levels. We also show that in postprandially high glucose, glucose-bound GK interacts and inhibits K_ATP_, as assessed by immunoprecipitation and FRET-FLIM analyses, as well as patch clamping. In addition, we have utilized GK mutants identified in diabetics, demonstrating that those mutants defective in binding to glucose impair GLP-1 secretion in however high glucose, whereas mutants constitutively bound to glucose with defects in catalysis, promote GLP-1 secretion even in low glucose. Furthermore, the K_ATP_ mutant unable to bind to GK blocks GLP-1 secretion in high glucose. We have also provided evidence showing that sodium-glucose cotransporter 1 (SGLT1) plays an important role in the basal secretion of GLP-1, but not in high glucose. We have thus revealed that glucose the molecule itself and GK act as a ligand and the receptor, respectively, to induce GLP-1 secretion in response to high glucose, providing new therapeutic avenues targeting metabolic diseases.

## INTRODUCTION

Glucagon-Like Peptide-1 (GLP-1) is produced as a product of alternative post-translational processing of proglucagon,^1–3^ primarily in the enteroendocrine L cells in the intestine,^3–9^ and preproglucagon neurons in the brain as well.^10^ GLP-1 has pleiotropic actions,^11^ including inhibition of appetite,^12,13^ protection of neuronal and cardiac cells,^14,15^ modulation of immune cells,^16^ and stimulation of natriuresis and diuresis,^17^ beyond its initial identification as an incretin for controlling glucose homeostasis.^18–22^ As an incretin, GLP-1 is secreted in response to high glucose during the early postprandial phase,^23–27^ and binds to its receptors (GLP1Rs) in the pancreas to stimulate insulin secretion, among other effects.^28–31^ In addition to glucose, GLP-1 secretion can also be enhanced by dietary lipids, such as triglycerides,^32,33^ 2-monoacylglycerol,^34^ and free fatty acids,^35^ as well as by proteins, including oligopeptides^36^ and amino acids.^37^ Pharmacological doses of GLP1 have been shown to improve control of blood glucose by promoting insulin secretion in the patients of type 2 diabetes mellitus (T2DM), with little risk of hypoglycaemia.^38–40^ GLP-1 receptor agonists (GLP1RAs) with longer half-lives than native GLP-1 have been developed,^41–44^ and are recommended as one of the first-line therapies for T2DM patients with a risk of atherosclerotic cardiovascular diseases or unable to tolerate metformin.^45,46^

In addition to glucose, the ATP-sensitive potassium channels (K_ATP_) play a crucial role in the release of GLP-1. K_ATP_ is closed in response to high glucose, leading to depolarization of the plasma membrane, which triggers secretion of GLP-1-containing vesicles.^47,48^ It has been shown that K_ATP_ inhibitors can directly promote the secretion of GLP-1 from L cell lines or primary L cells, bypassing the requirement of high glucose,^47^ whereas activation of K_ATP_ with its activators prevents glucose-induced GLP-1 secretion.^8,9^ Also because of the involvement of ATP in the closure of K_ATP_, it has been proposed that higher levels of ATP is generated in high glucose,^49^ especially due to the observation that the nonmetabolizable glucose analog 2-deoxy-D-glucose (2-DG) that cannot be converted to ATP fails to trigger GLP-1 secretion.^50,51^ Another critically important clue came from the indispensable role of GK (glucokinase; also known as hexokinase 4, HK4), which has the *K*_m_ of around 10 mM,^52,53^ and thus effectively carries out glucose catalysis when glucose levels reach higher than 10 mM, near or above.^54^ The GLUT2 glucose transporter, which is either present on or relocated to the apical (luminal) side of L cells^55–58^ has been shown to be critical for transport of high glucose,^56,57,59–61^, by virtue of its high *K*_m_ close to that of GK.^62^ In summary, the current understanding on the stimulation of GLP-1 release by high glucose is that the GLUT2-GK-ATP-K_ATP_ axis plays a pivotal role,^8,48,51,54,55^ similarly to that for the secretion of insulin from pancreatic β cells.^63–68^

However, there are some caveats to the above-mentioned mechanisms of GLP-1 secretion. Studies on maturity-onset diabetes of the young type 2 (MODY2) patients have shown that the GK mutations, which are defective in glucose catalysis and impair insulin secretion, do not display a deficiency in GLP-1 secretion.^69,70^ The sodium-glucose cotransporter 1 (SGLT1), which co-transports positively charged sodium ions along with glucose into L cells, has been elegantly shown to directly cause membrane depolarization to induce GLP-1 secretion,^71–73^ in parallel to the effect of membrane depolarization caused by K_ATP_ channels. In particular, it was shown that mice with intestine-specific depletion of *SGLT1* exhibit significantly impaired GLP-1 secretion as determined at 5 min after oral glucose gavage.^74^ In primary L cells and the L cell line GLUTag, treatment with the inhibitor of SGLT1 blocks the secretion of GLP-1 in the presence of 1 mM glucose.^72^ Depletion of extracellular sodium has also been shown to block GLP-1 secretion in these cells.^72^ Moreover, the observation that 2-DG cannot be co-transported by SGLT1, and is unable to stimulate GLP-1 secretion, was also attributed to the role of SGLT1.^50,51,75^ Also in support of the argument, methyl-α-D-glucopyranoside (MDGP, α-MG), another nonmetabolizable glucose analog that can be transported by both SGLT1 and GLUT2, is able to promote GLP-1 secretion.^50,75,76^

However, SGLT1 is saturated in glucose below 8 mM at most, ranging from 0.1 (ref. ^71,76,77^) to 8 mM (ref. ^56^), depending on the studies. In contrast, the secretion of GLP-1 remains unsaturated even at glucose levels exceeding 20 mM,^24,56,78–80^ a level commonly encountered in the intestinal lumen postprandially.^56,81^ Importantly, studies have indicated that high concentrations of glucose or the nonmetabolizable analog MDGP exceed the *K*_m_ of SGLT1 and hence saturate SGLT1,^56,75^ can lead to a greater amount of secretion of GLP-1 compared to glucose levels below the *K*_m_ of SGLT1,^9,24,71,79,80^ indicating that other mechanisms will account for a major portion of postprandial GLP-1 secretion.

Here, we have revealed that it is the physical binding of glucose itself to GK that directly triggers GLP-1 secretion, in dependency of structure rather energy of glucose. We have shown that the nonmetabolizable glucose analog MDGP, but not 2-DG, promotes GLP-1 secretion. The difference between the two nonmetabolizable analogs of glucose is attributed to their different ability to bind GK, i.e., MDGP can bind GK, whereas 2-DG cannot. We have also shown that, while SGLT1 is responsible for GLP-1 secretion at lower, basal glucose levels, consistent with its low *K*_m_ for glucose, GK plays a dominant role in mediating high glucose to stimulate the robust postprandial GLP-1 release. Mechanistically, we have demonstrated that when occupied by glucose or its analog, GK directly interacts with the Kir6.2 subunit of K_ATP_, leading to closure of the channel. Using a whole-cell patch clamping technique, we show that glucose or the analog stimulates GK to interact and close the K_ATP_ channel. We also performed inside-out membrane patch clamping in a reconstituted cell-free system, showing that addition of GK and glucose causes closure of the K_ATP_ channel on the membrane patch. Together, unlike the role of glucose in insulin secretion from the pancreatic β cells, glucose acts as a ligand-like stimulus, which at concentrations over the *K*_m_ is sensed by GK. The glucose-occupied GK in turn binds and inhibits K_ATP_ to trigger the secretion of GLP-1 from the intestinal L cells.

## RESULTS

### High glucose induces secretion of GLP-1 without increasing cellular ATP levels

We fasted mice for 16 h and re-fed them with a normal chow diet (67.4% saccharides) for 10 min at approximately 10 g/kg of body weight (modified based on ref. ^82^; Figure S1A). Stimulation of secretion of GLP-1, assessed for combined levels of both its full-length and two truncated forms in plasma, was observed from 5 to 45 min after the diet was given (Figure 1A; see also the accompanying insulin levels in Figure S1B), in line with previous reports,^27,83–87^. The course of the stimulation correlated positively with the concentrations of glucose in the duodenal and jejunal lumens (Figure 1A; no change in glucose levels in the lumen of the ileum). To our surprise, we found that the levels of intracellular ATP, as well as ADP and AMP, determined by mass spectrometry, did not change in the tissues of the duodenum and jejunum during the periods of GLP-1 secretion, compared to fasting (Figure 1A). We also did not find any increase of ATP or changes in the ratios of the adenylates in the duodenum and jejunum tissues from mice gavaged with 1.5 g/kg glucose (Figure 1B), or from mice directly injected with a solution of 75 g/L glucose (10 µL/g of body weight) to the duodenal and jejunal ducts via enterostomy (Figures 1C and 1D), which mimicked postprandial levels of glucose (Figures 1B-D). In addition, in dissected duodenal and jejunal tissues exposed ex vivo to glucose at concentrations similar to postprandial levels (above 12.5 mM; defined as “high glucose”), adenylate levels remained unchanged during GLP-1 secretion (Figures S1F-S1I). Moreover, to exclude the possibility that adenylate levels in non-L epithelial cells mask changes in adenylate levels in L cells, we performed cell sorting on intestinal epithelial cells, among which the L cells expressed GFP (green fluorescent protein) driven by the proglucagon promoter (ref. ^88^; also validated in Figure S1J). We then determined the levels of adenylates in these primary GFP-positive L cells in high glucose, and again observed unchanged intracellular adenylate levels (Figures 1E and 1F). Such a lack of increase of ATP was also seen in GLP-1-secreting L cell lines, including STC-1 (ref. ^89,90^) (Figures S1K and S1L), GLUTag^91^ (Figures S1M and S1N), and NCI-H716 (ref. ^92^) (Figures S1O and S1P). Since local intracellular increase of ATP, particularly near the plasma membrane, can also lead to the closure of K_ATP_ channels and induce insulin secretion in pancreatic β cells,^93,94^ we also determined the local intracellular concentrations of ATP in STC-1 cells. For this, we expressed the indicator that detects ATP:ADP ratios, PercevalHR,^95^ which is fused with Lck to its N-terminal to target this indicator to the plasma membrane (Figure 1G; validated in Figure S1Q). As shown in Figure 1H, we did not detect any change in local intracellular ATP:ADP ratios in these cells in high glucose. Therefore, glucose induces GLP-1 release from the intestinal L cells in an ATP-independent manner.

**Figure 1.**
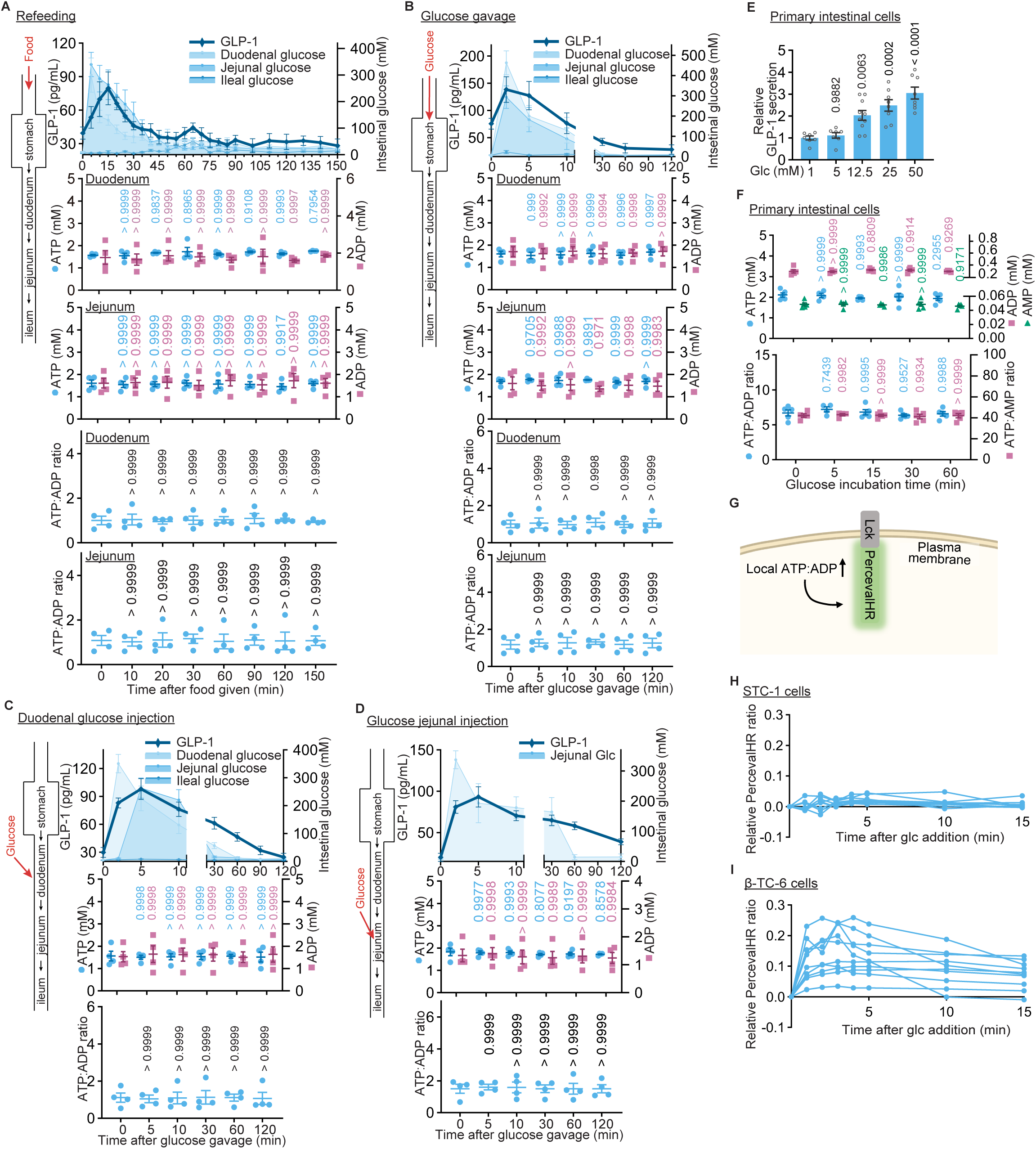
High glucose triggers GLP-1 secretion without elevating ATP in the intestinal L cells. (A-D) High glucose-induced secretion of GLP-1 is not accompanied with an increase of ATP levels in the intestinal tissues. Mice were fasted for 16 h and re-fed with normal chow diet for 10 min (A; Each mouse consumed food approximately 10 g/kg of body weight, as validated in Figure S1A), or were gavaged with 1.5 g/kg glucose (B), or injected via enterostomy with 75 g/L glucose directly into the duodenum (C), or jejunum (D) at a volume of 10 µL/g to mimic peak luminal glucose concentrations at around 5 min postprandial as measured in the upper panel of A). Mice were sacrificed at indicated time intervals after refeeding or glucose injection. The plasma GLP-1 (total levels, including full-length and two truncated forms) and the glucose concentrations in the intestinal lumen (upper panel) were measured, along with ATP and ADP levels (middle panel) and the ATP:ADP ratios (lower panel) in the intestinal tissues (by HPLC-MS). Serum insulin and blood glucose were also measured in Figures S1B-E. Results are shown as mean ± SEM; n = 8 (GLP-1 and luminal glucose of A), 7 (GLP-1 and luminal glucose of B), 6 (GLP-1 and luminal glucose of C and D), or 4 (others) mice for each time point, and *p* values by one-way ANOVA, followed by Dunnett (ATP and ADP levels of jejunum) or Tukey (others), all compared to the glucose-untreated group. (E and F) Glucose triggers secretion of GLP-1 from primary L cells without increasing ATP levels. Mice engineered to express GFP in L cells were fasted for 16 h, and primary intestinal epithelial cells (containing L cells) were isolated from small intestine tissues. These cells were either cultured in DMEM for 2 days, followed by two washes with PBS and subsequent incubation in GLP-1 KRBH buffer containing varying concentrations of glucose for 1 h to measure GLP-1 concentration in the medium (E), or were directly sorted using flow cytometry (F). The sorted cells were incubated in DMEM for 2 h, followed by two washes with PBS and subsequent incubation in GLP-1 KRBH buffer containing 25 mM glucose for the indicated durations to measure adenylate levels (F; by HPLC-MS). Results are shown as mean ± SEM; n = 10 (12.5 mM of E), 8 (25 mM and 50 mM of E), 7 (E, others), or 5 (F) biological replicates for each treatment, and *p* value by one-way ANOVA, followed by Dunnett (E) or Tukey (F), compared to the 1 mM Glc group (E) or the zero time point (F). (G-I) High glucose does not change local intracellular concentrations of ATP in L cells. STC-1 cells (H) and β-TC-6 cells, a pancreatic β cell line used as a control (I), were infected with a lentivirus that carried a plasma membrane-localized, Lck-fused PercevalHR indicator (depicted in G and validated in Figure S1Q). Cells were incubated in 25 mM glucose, and the ratios of fluorescent intensities of emitted light, excited at 490 nm versus 400 nm (F_490_/F_400_), were determined in real-time using confocal microscopy. The changes of F_490_/F_400_ relative to the resting (glucose-free) F_490_/F_400_ of the indicator, calculated by subtracting the resting values from the measured F_490_/F_400_ values, were then plotted, with n = 9 cells for H, and 10 cells for I. Experiments in this figure were performed three times, except the data of GLP-1 and glucose levels in the lumen of the duodenum, jejunum and ileum in A, B and C, which were repeated four times.

As a control, we also measured ATP levels in pancreatic β cells, which is increased in high glucose.^96,97^ We indeed observed an increase of approximately twofold in ATP levels in the β-TC-6 and INS-1 pancreatic cells (Figures S1T and S1U) when glucose concentrations were increased to 25 mM, which sufficiently stimulated secretion of insulin (Figures S1R and S1S). We also observed an increase of local intracellular ATP:ADP ratios beneath the plasma membrane in these cells (Figure 1I), consistent with the previous observations.^93,94^

### GK acts as a sensor of high glucose to trigger GLP-1 secretion

The requirement of high glucose yet without increased ATP production in L cells suggested a possibility that the presence of glucose per se may play a triggering role in GLP-1 secretion. Indeed, we found that gavaging of 1.5 g/kg nonmetabolizable MDGP into mice, which gave rise to peak concentrations of MDGP to 150 mM in the intestinal lumen after 5 min and decreased to below 10 mM after 30 min, induced GLP-1 release (Figure 2A; see also insulin secretion in Figure S2A). Similarly, addition of the nonmetabolizable MDGP instead of glucose into the culture medium of primary L cells or of STC-1 cells also induced the release of GLP-1 (Figures 2B and 2C), indicating that the structure of glucose is critical for inducing GLP-1 secretion. Of note, 2-DG, another glucose analog, did not stimulate GLP-1 release as does MDGP, the possible molecular explanation of which is to follow below (e.g., Figures 2M and S2E). As SGLT1 co-transports sodium ions along with glucose, we then analyzed whether SGLT1 plays a role in this process by treating STC-1 cells with the SGLT1 inhibitor phloridzin. We found that inhibition of SGLT1 blocked GLP-1 secretion during exposure to relatively low concentrations of glucose or MDGP (below 12.5 mM; defined as the “low” range of glucose/MDGP; Figures 2D and 2E), consistent with previous studies showing that phloridzin blocked GLP-1 secretion at 1 mM glucose.^72^ However, phloridzin had little effect on the robust GLP-1 secretion in “high” (above 12.5 mM) glucose and MDGP (Figures 2D and 2E). These results suggest that SGLT1 plays an important role in GLP-1 secretion in low glucose, but not in high glucose. As GLUT2 is a major glucose transporter for the uptake of high glucose by intestinal cells,^56,59–61^ we also investigated the role of GLUT2 that transports glucose in high concentrations, and found that inhibition of GLUT2 largely blocked GLP-1 secretion in high glucose, but not the GLP-1 secretion happening at low glucose (Figures 2F and 2G).

**Figure 2.**
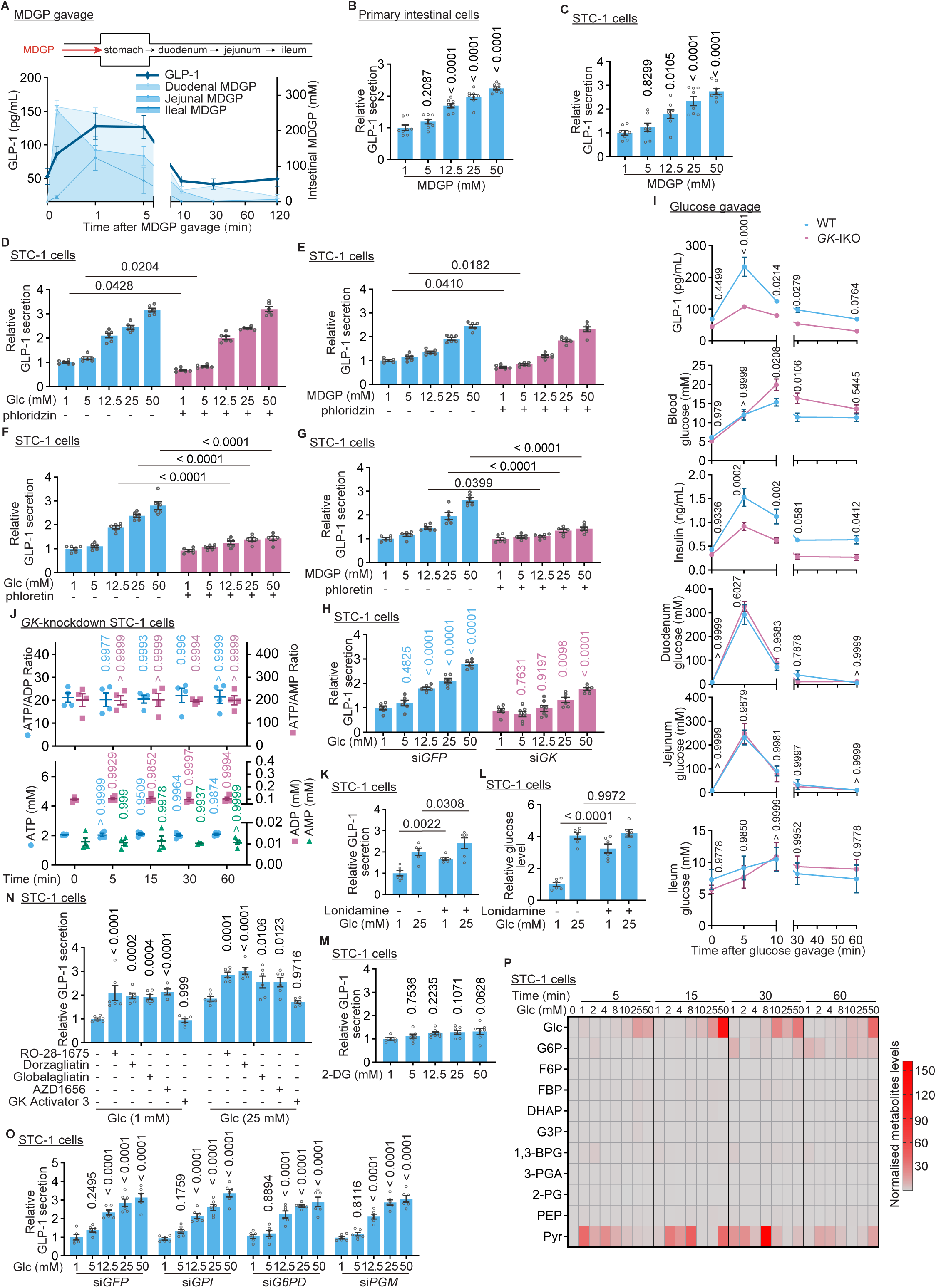
Glucokinase acts as a glucose-sensitive trigger for GLP-1 secretion. (A-C) Nonmetabolizable glucose analog MDGP, capable of binding to GK, promotes GLP secretion. GLP-1 in mice fasted for 16 h and gavaged with MDGP at 1.5 g/kg (A), or from the MDGP-treated primary intestinal L cells (B) and STC-1 cells (C) cultured in glucose-free GLP-1 KRBH buffer, was determined alongside blood levels of glucose (A), and duodenal, jejunal, and ileal MDGP (A), and serum insulin in Figure S2A (for A). Results are shown as mean ± SEM; n = 8 mice (A) or biological replicates (B and C) for each time point/MDGP concentration, and *p* value in B and C by one-way ANOVA, followed by Dunnett (B) or Tukey (C), all compared to the 1 mM MDGP group. See also validation data regarding the binding affinity of MDGP to GK in Figure S2D. (D-G) SGLT1 regulates GLP-1 secretion in low glucose levels. STC-1 cells were incubated for 1 h in a glucose-free GLP-1 KRBH buffer, supplemented with either glucose (D and F) or MDGP (panels E and G) at indicated concentrations, together with 5 µM phloridzin, an SGLT1 inhibitor (D and E) or 100 µM phloretin, a GLUT2 inhibitor used as a control (F and G). Results are shown as mean ± SEM; n = 6 biological replicates for each concentration of glucose or MDGP, and *p* value by two-way ANOVA, followed by Tukey. (H and J) GK is required for postprandially high glucose-induced secretion of GLP-1 in L cells. *GK*-knockdown STC-1 cells (validated in Figure S2B) were incubated in GLP-1 KRBH buffer containing varying concentrations of glucose for 1 h (H) or 25 mM glucose for indicated time durations (J), followed by measurement of GLP-1 secretion (H) and the analysis of intracellular adenylate levels (J; by CE-MS). Results are shown as mean ± SEM; n = 7 (H, 5 mM and 12.5 mM of *GK*-knockdown cells), 6 (H, others), or 4 (J) biological replicates for each glucose concentration/time point, and *p* value by two-way (H) or one-way (J) ANOVA, followed by Sidak (H, for comparisons within each genotype against the 1 mM glucose group) or Dunnett (J, for comparisons against the glucose untreated group). (I) GK is required for high glucose-induced secretion of GLP-1 in mice. Mice with intestine-specific *GK* knockout (*GK*-IKO, validated in Figure S2C) were starved for 16 h, and then gavaged with 1.5 g/kg glucose. Plasma GLP-1 (first panel) and serum insulin (third panel) levels, alongside blood (second panel), duodenal (fourth panel), jejunal (fifth panel), and ileal (sixth panel) glucose levels, were measured. Results are shown as mean ± SEM; n = 6 mice for each time point, and *p* value by two-way ANOVA, followed by Sidak (comparisons between different genotypes at each time point). (K and L) Elevation of intracellular glucose levels resulting from inhibition of HK2 promotes GLP-1 secretion. STC-1 cells were treated with the HK2 inhibitor lonidamine at 100 µM for 1 h, followed by determination of GLP-1 secretion (K) and intracellular glucose (L) after an additional 1-h incubation of either 1 mM or 25 mM glucose (supplemented in GLP-1 KRBH buffer). Results are shown as mean ± SEM; n = 6 biological replicates for each treatment, and *p* value by two-way ANOVA, followed by Sidak. (M) Nonmetabolizable glucose analog 2-DG, unable to bind GK, does not promote GLP-1 secretion. STC-1 cells were treated with indicated concentrations of 2-DG (supplemented in GLP-1 KRBH buffer) for 1 h, followed by determination of GLP-1 secretion. Results are shown as mean ± SEM; n = 6 (1 mM of M), or 7 (others) biological replicates for each concentration, and *p* value by one-way ANOVA, followed by Dunnett, all compared with the 1 mM 2-DG group. See also validation data regarding the binding affinity of 2-DG to GK in Figure S2E. (N) GK activators that increase the affinity of GK for glucose reduce the threshold concentrations of glucose for stimulating GLP-1 secretion. STC-1 cells were treated with GK activators that increase the affinity of GK towards glucose, including dorzagliatin, RO-28-1675, globalagliatin, and AZD1656, all at 10 µM for 1 h, followed by determination of GLP-1 secretion after an additional 1-h incubation with different concentrations of glucose. As a control, cells were treated with 10 µM GK Activator 3, which increases catalytic activity of GK but without affecting its glucose binding affinity, for 1 h. Results are shown as mean ± SEM; n = 5 (AZD1656, 1 mM glucose) or 6 (others) biological replicates for each treatment, and *p* value by two-way ANOVA, followed by Sidak (comparisons within each glucose concentration against the drug-untreated group). (O) Enzymes downstream of GK are not required for secretion of GLP-1 induced by high glucose. STC-1 cells with knockdown of *GPI*, *G6PD* (both *G6PD2* and *G6PDx*), or *PGM* (both *PGM1* and *PGM2*) were treated with indicated concentrations of glucose for 1 h, followed by determination of the GLP-1 secretion. Results are shown as mean ± SEM; n = 6 biological replicates for each treatment, and *p* value by two-way ANOVA, followed by Dunnett, all compared to the 1 mM Glc group. See also validation data for the knockdown efficiency of each gene in Figure S2G. (P) GLP-1 secretion is positively correlated with intracellular glucose concentrations, but not glycolytic intermediates. STC-1 cells were treated with varying concentrations of glucose for indicated time durations, followed by determination of intracellular levels of glucose and glycolytic intermediates, using CE-MS (for FBP) and HPLC-MS (for others). Results are shown as heatmap, with accompanying color scale (right panel) indicating the average metabolite levels (normalized to the glucose-free group), ranging from grey (lowest levels) to red (highest levels). The values are scaled row-wise to reflect relative level patterns across different glucose treatment concentrations. See also the complete list of relative levels of individual metabolites, shown as dot plots, in Figure S3. Experiments in this figure were performed three times, except P six times.

It is intriguing that GK, also known as hexose kinase 4, has a *K*_m_ of around 10 mM for glucose,^52^ much higher than the *K*_m_ values other hexokinases (HK1-3).^98^ We pondered whether it is GK that responds to high glucose and acts as the molecular sensor of glucose. Indeed, when *GK* was knocked down in STC-1 cells (validated in Figure S2B) or specifically knocked out in the intestine (*GK*-IKO, validated in Figure S2C), we observed impaired secretion of GLP-1 following incubation in high glucose or glucose gavage (Figures 2H and 2I). Surprisingly, we found that the total levels of cellular ATP and ADP, or their ratios, remained unchanged in the *GK*-knockdown cells (Figure 2J). These observations also suggested that GK controls GLP-1 secretion in L cells. Counterintuitively, we found that inhibition of HK2 by lonidamine,^99^ promoted GLP-1 secretion, even at low glucose levels (1 mM, Figure 2K), which was likely due to an accumulation of glucose inside the cell as a consequence of reduced glucose catabolism after inhibition of HK2 (Figure 2L). As mentioned above, we found that 2-DG did not induce GLP-1 secretion as does MDGP (Figure 2M). To investigate why the two glucose analogs MDGP and 2-DG behave differently in the stimulation of GLP-1 secretion, we compared the binding affinities of MDGP and 2-DG to GK. We found that MDGP had an affinity for GK with a *K*_D_ of around 25 mM (Figure S2D), whereas 2-DG did not bind to GK (Figure S2E). The stark difference in binding to GK provides an explanation for why only MDGP, and not 2-DG, can mimic glucose to stimulate GLP-1 release (Figures 2B, 2C, and 2M). Furthermore, the relatively lower affinity of GK for MDGP (higher *K*_D_ or *K*_M_) compared to glucose is in line with the requirement of higher concentrations of MDGP (above 12.5 mM) to stimulate GLP-1 secretion to comparable levels (Figures 2D and 2E). In contrast, MDGP had a similar affinity for SGLT1 to glucose,^75^ and stimulated GLP-1 secretion as effectively as glucose, at the low concentration range of glucose (below 12.5 mM) – as abovementioned, SGLT1-mediated glucose response only during the low range of glucose (Figures 2D and 2E). Furthermore, we found that STC-1 cells treated with GK activators, which increase the affinity of GK towards glucose (decreases *K*_m_), including dorzagliatin,^100^ RO-28-1675,^101^ globalagliatin (validated in Figure S2F), and AZD1656,^100^ the glucose concentrations needed to secrete a similar amount of GLP-1 are lower compared to untreated cells (Figure 2N). Such an effect was not observed with the compound GK Activator 3 (Figure 2N), which increases only catalytic activity (V_max_) without affecting the *K*_m_ of GK.^102^ As a control, knockdown of the glycolytic enzymes downstream of GK, including glucose-6-phosphate isomerase (GPI), glucose-6-phosphate dehydrogenase (G6PD), or phosphoglucomutase (PGM), did not affect GLP-1 secretion in high glucose (Figure 2O; validated in Figure S2G; note that glucose-6-phosphatase (G6Pase) was not targeted for knockdown or expression, as it was below detectable levels in L cells, as shown in Figure S2H). As an additional control, we found that GLP-1 secretion positively correlated with intracellular glucose concentrations in cultured STC-1 cells, but not with those of the glycolytic metabolites (Figure 2P, Figure S3).

In humans, loss-of-function mutations of GK have been identified in MODY2 patients.^103,104^ Among them, the G80A mutation is unable to bind and hence unable to catalyze glucose as a result of much reduced V_max_ and a much increased *K*_m_, as validated in Figures S4A and S4B. We found that, ectopic expression of G80A in wildtype STC-1 cells or re-introduction of G80A into *GK*-knockdown STC-1 cells impaired GLP-1 secretion induced by high glucose (Figures 3A and 3B), but had no effect on low glucose-induced GLP-1 secretion which was inhibited by SGLT1 inhibitor phloridzin (Figure 3C). These observations support the notion that SGLT1 but not GK plays a dominant role in GLP-1 secretion in low glucose. In contrast, the L309P mutation of GK is defective in catalysis and hence is constitutively in a glucose-bound state even in low glucose, due to a much lower V_max_ with unchanged *K*_m_ towards glucose, as validated in Figure S4C. We found that this glucose-bound mutant L309P constitutively stimulated GLP-1 secretion (Figures 3A and 3B). In addition, the inhibition of SGLT1 by phloridzin only marginally reduced GLP-1 secretion from cells expressing GK-L309P that mimics high glucose (Figure 3C), reinforcing the importance of GK in GLP-1 secretion in higher range of glucose levels compared to SGLT1. Neither GK-G80A nor GK-L309P changed intracellular ATP levels in these cells (Figures S4E-S4G; see validation data in Figure S4D), similar to the result seen in *GK* knockdown (Figure 2J). The opposite effects of GK-G80A and GK-L309P on the secretion of GLP-1 were recapitulated in mice when knocked-in into the intestines of mice with the background of intestine-specific knockout of GK (*GK*-IKO) (Figure 3E, validated in Figure S4L). As a control, we also examined the effects of GK-G80A and GK-L309P on insulin secretion, and found that both mutations impaired insulin secretion in high glucose when re-introduced into the *GK*-knockdown β-TC-6 cells (Figure 3D, validated in Figure S4H). These data are consistent with the MODY2 patients carrying the GK-G80A and GK-L309P mutations exhibiting deficiency in insulin secretion,^103,104^ and with the requirement of increased ATP production in high glucose (Figures S4I-S4K). In addition, we identified another MODY2 patient carrying a heterozygous R397L mutation of GK (Figure S5A), which is constitutively occupied with glucose yet unable to catalyze it due to a reduced V_max_ (enzymatic data listed in Figure S5B). Furthermore, we found an additional MODY2 patient who has a heterozygous, truncated form of GK, the mRNA for which contains a repeated nucleotide sequence from position 242 to 258, and hence encodes GK with a frameshift mutation at amino acid (aa) 87 (p.M87Afs*6; Figure S5C). This frameshift alters the amino acids from positions 87 to 91 and leads to premature termination of translation at amino acid 92 (referred to as GK^Δ92-CT^), yielding a truncated GK lacking the glucose-binding domain (Figures S5C and S5D). We found that GLP-1 secretion was much lower in the patient carrying the GK^Δ92-CT^ truncation compared to the patient with the R397L mutation, despite both patients having similar blood glucose levels after glucose intake (Figure 3F). Both patients exhibited impaired glucose-induced insulin secretion, with insulin-to-glucose ratios around 20 pmol/mmol (Figure 3F), which is typical for MODY2 patients, whereas healthy individuals typically have ratios close to 40 pmol/mmol.^105–107^ These blunting effects of GK-L309P and GK^Δ92-CT^ on GLP-1 secretion could also be recapitulated in STC-1 cells after introducing these mutations into the *GK*-knockdown STC-1 cells (Figure 3G; see validation data in Figure S5E). In addition, both mutations, when reintroduced into β-TC-6 cells, failed to rescue the knockdown of *GK* in the production of ATP and insulin secretion in pancreatic β-cells (Figures 3H, S5F, and S5G; see validation data in Figure S5E). Together, these mutational studies indicate that the binding of glucose to GK underlies GLP-1 secretion.

**Figure 3.**
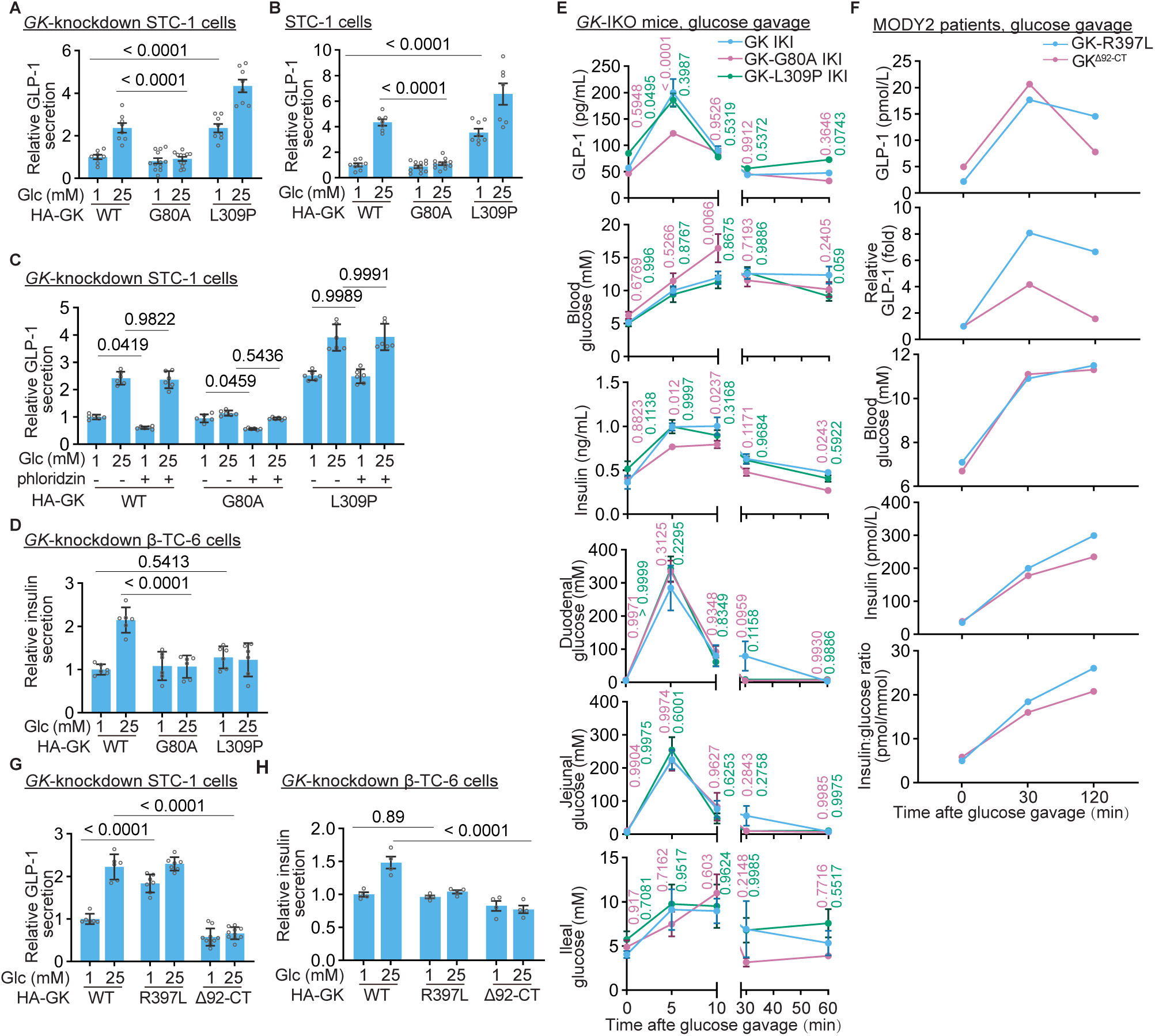
The affinity for glucose underlies the ability of GK to trigger GLP-1 secretion. (A and B) Glucose-binding affinity of GK, rather than catalytic activity, determines its ability to induce GLP-1 secretion in L cells. Wildtype (WT) GK, its glucose-binding-defective G80A mutant, and the glucose-constitutive-binding L309P mutant, were separately expressed in wildtype STC-1 cells (A) or re-introduced into *GK*-knockdown STC-1 cells (B) via lentiviral infection. Cells were then treated with indicated concentrations of glucose for 1 h, followed by determination of GLP-1 secretion. Data are shown as mean ± SEM; n = 7 (A; 25 mM of WT, and L309P, B; L309P), 12 (G80A), or 8 (others) biological replicates for each treatment, and *p* value by two-way ANOVA, followed by Tukey (A) or Sidak (B). See also the enzymatic properties of GK and the two mutants in Figures S4A-C. (C) GK plays a dominant role in sensing glucose of higher levels to stimulate GLP-1 secretion, compared to SGLT1. Wildtype GK or G80A and L309P mutants were individually re-introduced into the *GK*-knockdown STC-1 cells. Cells were incubated in different concentrations of glucose, together with 5 µM phloridzin, for 1 h, followed by determination of GLP-1 secretion. Data are shown as mean ± SEM; n = 6 biological replicates for each treatment, and *p* value by two-way ANOVA, followed by Tukey. (D) GK-G80A and GK-L309P mutants, both unable to catalyze conversion of glucose to ATP, impair insulin secretion in pancreatic β cells. Wildtype GK and its G80A and L309P mutant were separately re-introduced into the *GK*-knockdown β-TC-6 cells. Cells were then treated with indicated concentrations of glucose for 1 h, followed by determination of insulin secretion. Data are shown as mean ± SEM; n = 6 biological replicates for each treatment, and *p* value by two-way ANOVA, followed by Tukey. See also adenylate levels in these cells in Figures S4I-K. (E) Glucose-binding abilities of GK proteins underlie GLP-1 secretion in mice. Mice with intestinal GK replaced with either GK-G80A, GK-L309P, or wildtype GK as a control (GK mutants were knocked-in (KI) into the intestine of mice with intestine-specific knockout of *GK*; validated in Figure S4L) were gavaged with glucose at a dosage of 1.5 g/kg. Plasma GLP-1 and serum insulin levels, along with blood and intestinal glucose levels, were measured. Results are shown as mean ± SEM; n = 6 mice for each time point/genotype, and *p* value by two-way ANOVA (comparisons between different genotypes), followed by Tukey (jejunal glucose) or Dunnett (others). (F-H) Glucose-binding abilities of GK determine GLP-1 secretion in humans. Human patients carrying either the glucose-constitutively-bound GK-R397L mutant (validated in Figure S5B) or the glucose-binding-defective GK^Δ92-CT^ mutant (validated in Figure S5D) took glucose orally at a dose of 1 g/kg, dissolved in a total volume of 250 mL of water. Levels of GLP-1 (F; upper panel), insulin (F; lower panel), and blood glucose (F; middle panel) were measured at indicated time points after glucose intake, with data shown as mean ± SEM. As controls, the GK-R397L and GK^Δ92-CT^ mutants were re-introduced into the *GK*-knockdown STC-1 (G) and β-TC-6 cells (H), followed by assessing GLP-1 (G) or insulin (H) secretion after 1 h of high glucose treatment (results are shown as mean ± SEM; n = 6 (G; GK-WT), 7 (G; GK-R397L), 10 (G; GK^Δ92-CT^) or 4 (H) biological replicates for each genotype/condition, and *p* value by two-way ANOVA, followed by Sidak (G) or Tukey (H). Experiments in this figure were performed three times.

### Glucose-occupied GK binds and inhibits K_ATP_

We next investigated how GK in its glucose-bound state triggers GLP-1 secretion. We found that the GLP-1 secretion induced by the constitutively-glucose-bound GK-L309P could be blocked by the K_ATP_ channel openers diazoxide and tifenazoxide (NN414) (Figures 4A and S6A), indicating that the GK-induced GLP-1 secretion depends on the closure of K_ATP_ channels. Consistently, the K_ATP_ inhibitors glibenclamide and repaglinide could directly induce GLP-1 secretion even in the presence of the glucose-binding-defective GK-G80A, effectively bypassing the dominant negative effect of this mutant on GLP-1 secretion (Figures 4B and S6B). These results suggest that K_ATP_ functions downstream of GK binding to glucose in GLP-1 secretion (depicted in Figure 4C).

**Figure 4.**
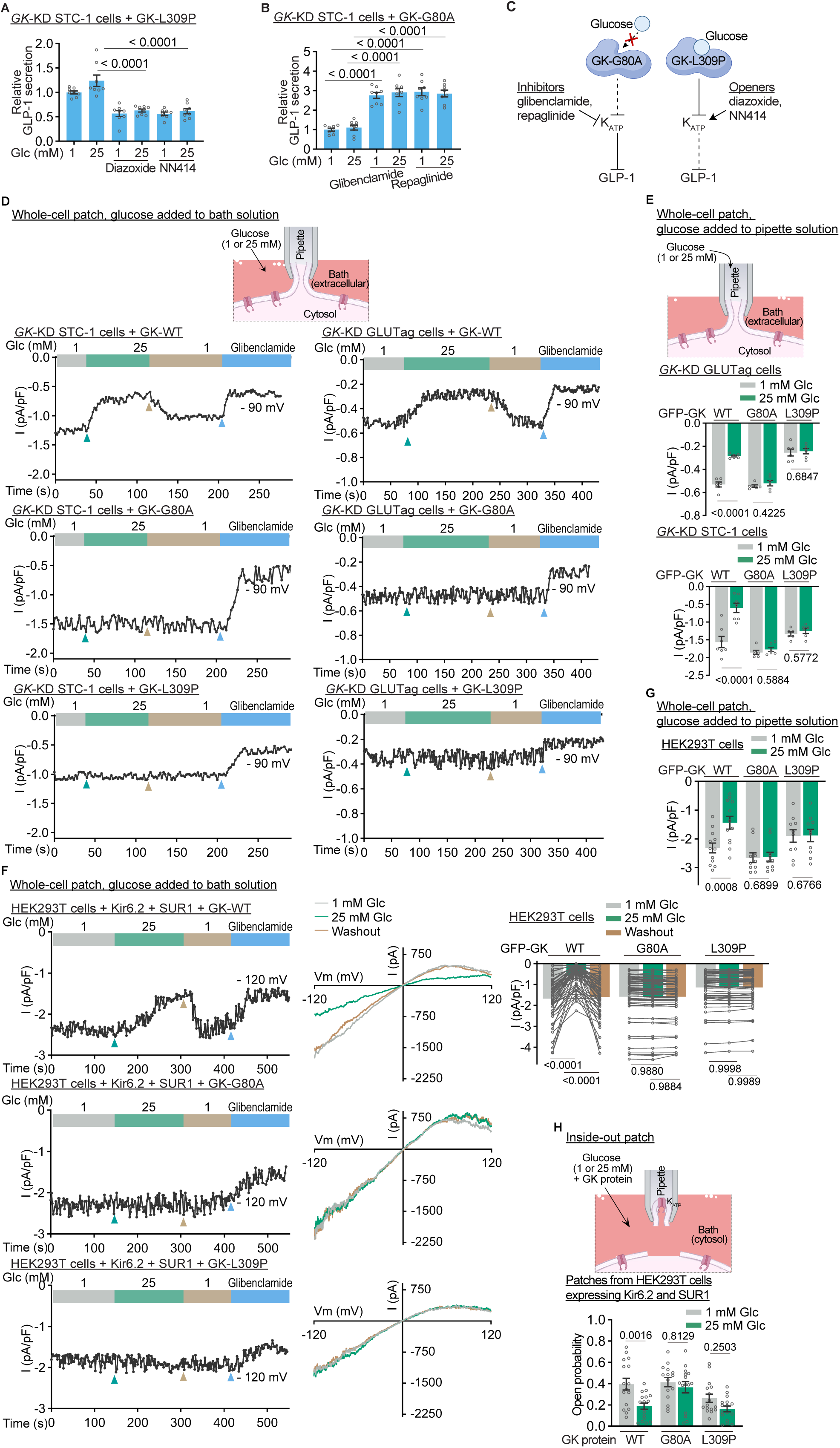
Glucose binding is a prerequisite for GK to inhibit K_ATP_. (A and B) K_ATP_ activity is modulated by glucose binding of GK in controlling the secretion of GLP-1. *GK*-knockdown (*GK*-KD) STC-1 cells were reintroduced with glucose-binding-defective GK-G80A (B) or glucose-constitutive-binding GK-L309P (A). Cells were then treated for 1 h with K_ATP_ inhibitors glibenclamide and repaglinide (B) or K_ATP_ openers diazoxide and NN414 (A), at a concentration of 10 µM. After the treatment, GLP-1 secretion was measured following an additional 1-h incubation with different concentrations of glucose. Results are shown as mean ± SEM; n = 8 biological replicates for each treatment, and *p* value by one-way (B) or two-way (A) ANOVA, followed by Tukey. (C) Schematic illustrating K_ATP_ channels are closed on binding to glucose-bound GK in the stimulation of GLP-1 secretion by high glucose. (D-G) GK in glucose-bound state inhibits K_ATP_. *GK*-KD STC-1 (D and E), *GK*-KD GLUTag cells (D and E), or the volume-regulated anion channels (VRAC)-deficient/*LRRC8A*-KO HEK293T cells pre-transfected with Kir6.2-mCherry and Myc-SUR1 (F and G), were infected with lentivirus carrying GFP-tagged GK-G80A or GK-L309P, and then starved for glucose for 2 h. The plasma membrane of the individual cells was then suctioned using a glass micropipette (containing buffer with 2 mM ATP and 200 μM ADP), forming a gigaohm seal (illustrated in the upper panels of D and E). The membrane was then ruptured to achieve whole-cell access, followed by recording the whole-cell K^+^ current evoked by a linear voltage ramp from −140 mV to +60 mV (D and E) or from −120 mV to +120 mV (F and G) over a duration of 400 msec, applied repeatedly every 2 sec. Glucose were either supplemented in the bath solution (extracellular), which was varied from 1 mM (depicted as in grey bar) to 25 mM (green bar), and then returned to 1 mM (washout with 1 mM solution; yellow bar), followed by treatment with 10 µM glibenclamide (blue bar) using a perfusion system during the recording (D and F), or maintained in the pipette solution (cytosolic face) at a constant concentration during the recording, with an 5-min equilibration period after achieving whole-cell access (E and G). The representative trace (lower panel of D, and left panel of F) and the statistical analysis (lower panel of E, G, and right panel of F) of the ramp-induced currents recorded at −90 mV (D and E) or −120 mV (F and G) are shown (mean ± SEM, n = 7 (STC-1 + GK-WT of E), 6 (E, others), 60 (GK-G80A of F), 59 (F, others), 12 (GK-WT of G), or 8 (G, others) cells for each genotype/treatment, with the *p* value determined by two-way ANOVA, followed by Tukey). In the middle panel of F, the representative steady-state current-voltage (I-V) curves, measured under conditions where current density (cell capacitance) varied by less than 1 pA/pF at each glucose concentration (indicated by triangles in the upper middle panel as an example), are shown. (H) GK in glucose-bound state inhibits K_ATP_ in vitro. HEK293T cells were transfected with Kir6.2 and SUR1 subunits of K_ATP_, followed by excising a membrane patch and exposing it in the bath solution (intracellular side) containing 2 mM ATP, 200 μM ADP, varying concentrations of glucose, along with bacterially expressed and purified wildtype GK, GK-G80A, and GK-L309P, Steady-state current was continuously recorded at a constant voltage of +60 mV (illustrated in the left panel). The channel open probability was quantified as the percentage of the total opening time within a 1-min analysis window. Data are shown as mean ± SEM; n = 17 (WT, 25 mM glucose), 15 (G80A) or 16 (others) membrane patches for each condition, with the *p* value determined by two-way ANOVA, followed by Sidak. See also representative 20-sec current traces for wildtype GK in Figure S6G, with downward deflections indicating the channel openings. Experiments in this figure were performed three times.

We also performed whole-cell patch clamping experiments to examine the effect of GK on K_ATP_ channel activities. We set the concentrations of ATP and ADP respectively at 2 mM and 200 μM (in the pipette buffer, as depicted in Figures 4D and 4E), to resemble their respective intracellular concentrations (measured in Figures 1F, S1L, and S1N), with concentrations of glucose varying. We found that high concentrations of glucose (25 mM), whether introduced in the pipette (representing intracellular compartment; Figure 4E) or bath solutions (representing extracellular compartment; Figure 4D), led to a decrease in the endogenous whole-cell K^+^ currents in L cell lines, including GLUTag cells and STC-1 cells (Figures 4D and 4E). When we washed out the glucose with a low-glucose (1 mM) bath solution, the currents were restored (Figure 4D). Such a glucose-, rather than ATP-regulated K_ATP_ activity could also be observed in HEK293T cells (lacking expression of GK^108^ and K ^109,110^) transfected with GK and the Kir6.2 and SUR1 subunits (the type of K_ATP_ expressed in L cells, as confirmed by mass spectrometry; Figure S6C) of the K_ATP_ (Figures 4F and 4G; see validation data in Figure S6D). In addition, MDGP also inhibited the K^+^ currents (Figure S6E). We also found that expression of GK-L309P, constitutively glucose-bound, could still inhibit the currents in low glucose in these cells, while GK-G80A, defective in glucose binding, was unable to inhibit the currents even in high glucose (Figures 4F and 4G; see validation data for GLUTag cells in Figure S6F).

We further reconstituted the regulation of GK on K_ATP_ by using membrane patches from HEK293T cells transfected with Kir6.2 and SUR1, and recorded currents from inside-out membrane patches. The purified bacterially expressed GK, along with 2 mM ATP, 200 μM ADP, and varying concentrations of glucose, was then added to the bath solution facing the intracellular side of K_ATP_ (depicted in Figure 4H). Consistent with previous findings,^111^ we observed a basal open-state probability (*p*_open_) of K_ATP_ in low glucose (Figures 4H and S6G). When high glucose was introduced, the open-state probability of K_ATP_ reduced (Figure 4H). In addition, GK-G80A stimulated the opening of K_ATP_ in high glucose, while GK-L309P reduced the opening of K_ATP_ in high glucose even in low glucose (Figure 4H). These results indicate that GK can directly modulate the activity of K_ATP_.

Finally, to explore how GK regulates K_ATP_, we tested if GK forms a complex with the subunits (Kir6.2 and SUR1) of K_ATP_. Through immunoprecipitation, we found that when GK was ectopically expressed alongside either Kir6.2 or SUR1 in HEK293T cells, GK interacted with each of the subunits (Figures 5A-5D). We also found that the interaction between GK and the Kir6.2 subunit was significantly enhanced with the addition of high glucose in the culture medium or in cell lysates (Figures 5A and 5B), whereas the interaction between GK and SUR1 was constitutive (Figures 5C and 5D). In STC-1 cells, we also observed similar interaction patterns between endogenous GK-Kir6.2 and GK-SUR1 (Figures 5E and 5F). These observations are consistent with several studies showing that the association between Kir6.2 and SUR1 becomes more robust when the channel is inhibited,^112–115^ and further support the notion that the interaction of GK with Kir6.2 is enhanced for the inhibition of the K_ATP_ channel. We also found that the GK-L309P mutant showed a robust, constitutive interaction with Kir6.2, whereas the GK-G80A mutant showed marginal interaction with Kir6.2 (Figure 5G). Neither of the mutations affected the binding of GK to SUR1 (Figure 5H). The glucose-induced interaction between GK and Kir6.2 could also be observed inside the living cells by FRET-FLIM analyses (Figure 5I). As a control, we found that other hexokinases (HK1 to HK3) hardly interacted with K_ATP_, at either low or high glucose (Figures S7A-S7C). Domain mapping experiments showed that the segment of aa 205-228 in Kir6.2 was required for its interaction with GK (Figure 5J and S7D-S7F). For the Kir6.2^Δ205–228^ truncation showed severe degradation (Figure S7F), we created a mutation in which amino acids 205-228 were replaced with alanine (Kir6.2-205-228A). This mutant did not affect the activity of K_ATP_ in low glucose (at the open state; Figure S7G); however, it blocked the GK-mediated inhibition of K_ATP_ in high glucose (Figure S7G), and consequentially impaired the high glucose-induced secretion of GLP-1 when expressed in STC-1 cells or knocked-in into the intestines of mice with the background of intestine-specific knockout of Kir6.2 (*Kir6.2*-IKO) (Figures 5K and 5L; see validation data in Figure S7H). Together, we found that GK, when bound to glucose, interacts with and inhibits K_ATP_ to trigger the release of GLP-1.

**Figure 5.**
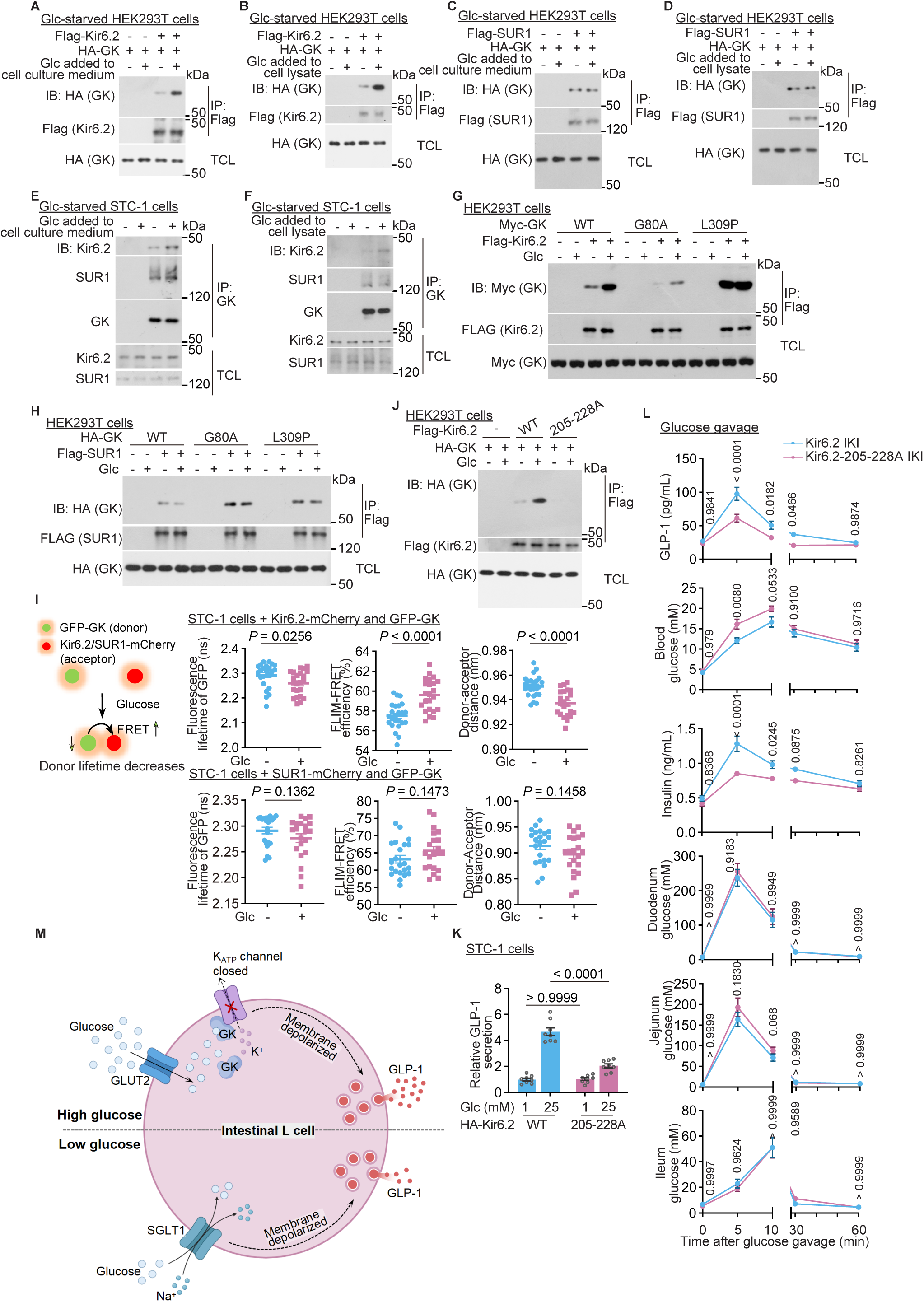
Glucose stimulates GK to interact with K_ATP_. (A-F) GK interacts with both the Kir6.2 and SUR1 subunits of K_ATP_, with the GK-Kir6.2 interaction enhanced by glucose and GK-SUR1 constitutive. HEK293T cells transfected with different combinations of HA-tagged GK, FLAG-tagged Kir6.2 (A and B), and FLAG-tagged SUR1 (C and D) for 24 h, and STC-1 cells (E and F) were used. Cells were treated with 0 mM or 25 mM glucose, which was either included in the DMEM culture medium for a 2-h incubation (A, C, and E) or added into the lysates after cell lysis (B, D, and F; mannitol was used to adjust the total concentration of glucose or mannitol alone to 25 mM, and the same hereafter for all immunoprecipitation assays involving glucose supplementation in the lysates). Immunoprecipitation of the FLAG epitope tag (A-D) or the endogenous GK (E and F) using the above cell lysates was performed. The co-immunoprecipitated GK (A-D) or K_ATP_ (E and F) was determined by immunoblotting. (G and H) GK in glucose-bound state displays enhanced binding to Kir6.2. HEK293T cells were transfected with different combinations of HA-tagged GK or its G80A and L309P mutants, FLAG-tagged Kir6.2 (G), and FLAG-tagged SUR1 (H). At 24 h post-transfection, cells were incubated in DMEM medium containing 0 mM or 25 mM glucose, followed by immunoprecipitation of the FLAG epitope tag and immunoblotting using the indicated antibodies. (I) GK interacts with K_ATP_ in living L cells. STC-1 cells infected with a lentivirus carrying Kir6.2-mCherry (upper three panels) or mCherry-SUR1 (lower three panels), along with GFP-GK, were incubated in DMEM medium containing 0 mM or 25 mM glucose for 2 h, followed by determination of the fluorescence lifetime of GFP of the FRET donor (see the assay strategy in the left panel), calculation of the FRET-FLIM efficiency, and estimation of donor-acceptor distance. Data are shown as mean ± SEM; n = 24 (0 mM glucose of Kir6.2), 23 (25 mM glucose of Kir6.2), 22 (0 mM glucose of SUR1) or 21 (25 mM glucose of SUR1) cells for each treatment, with the *p* value determined by two-sided Mann-Whitney test (GFP lifetime of SUR1) or unpaired two-sided Student’s *t*-test (others). (J) GK interacts with K_ATP_ via the region of aa 205-228 in Kir6.2. HEK293T cells were transfected with FLAG-tagged Kir6.2 or FLAG-tagged Kir6.2-205-228A, in which aa 205-228 required for GK binding (as determined in Figures S7D-S7F) were mutated to alanine. At 24 h post-transfection, cells were incubated in DMEM medium containing 0 mM or 25 mM glucose, followed by immunoprecipitation of the FLAG epitope tag and immunoblotting using the indicated antibodies. (K) Interaction of GK with K_ATP_ is required for glucose to stimulate GLP-1 secretion by high glucose. STC-1 cells infected with a lentivirus carrying wildtype Kir6.2 or Kir6.2-205-228A, which both exhibit similar channel activity (validated in Figure S7G), were treated with 25 mM glucose for 1 h, followed by determination of GLP-1 secretion. Data are shown as mean ± SEM; n = 8 biological replicates for each genotype/treatment, with the *p* value determined by two-way ANOVA, followed by Sidak. (L) The abilities of GK proteins to interact with K_ATP_ underlie the stimulation of GLP-1 secretion by high glucose. Mice with intestinal Kir6.2 replaced with either the GK-binding-defective Kir6.2-205-228A mutant, or wildtype Kir6.2 as a control (Kir6.2 was knocked-in into the intestine of mice with intestine-specific knockout of *Kir6.2*; validated in Figure S7H) were gavaged with glucose at a dosage of 1.5 g/kg. Plasma GLP-1 and serum insulin levels, along with blood and intestinal glucose levels, were measured. Results are shown as mean ± SEM; n = 6 mice for each time point/genotype, and *p* value by two-way ANOVA (comparisons between different genotypes), followed by Sidak. (M) Schematic illustrating how high glucose acts in a ligand-like manner to stimulate secretion of GLP-1. Glucose increased in early postprandial period is absorbed from the intestinal lumen into the L cells by GLUT2 transporters. Inside the L cells, GK becomes occupied with glucose, and interacts with the K_ATP_ channel, causing the closure of the K_ATP_ channel. As a result, the plasma membrane of the L cells becomes depolarized, stimulating the exocytosis of vesicles to release GLP-1 therein into the bloodstream (shown in the upper part of the diagram). In comparison, glucose at low levels is transported by SGLT1, which co-transports sodium ions into L cells, directly causing membrane depolarization and consequently secretion of GLP-1 (lower part). Experiments in this figure were performed three times, except A-D, G, and H, five times.

## DISCUSSION

We have provided multiple lines of evidence showing that glucose acts as a direct stimulus to induce the release of GLP-1 in L cells, in the sense that the glucose molecule physically triggers the secretion (Figure 5M). First of all, we found that high glucose stimulates GLP-1 secretion in L cells without increasing ATP levels. In particular, the nonmetabolizable glucose analog MDGP, unable to generate ATP but still binds to GK, effectively inhibits the K_ATP_ channel and stimulates GLP-1 secretion. In support of a requirement of physical binding of glucose, the other analog 2-DG, which does not bind to GK, fails to stimulate GLP-1 release. Second, we have employed various GK mutants identified in human MODY2 patients,^103,104^ and found that constitutively-glucose-occupied GK mutants can inhibit K_ATP_ and promote GLP-1 secretion even in low glucose, whereas glucose-binding-defective GK mutants fail to stimulate GLP-1 secretion in high glucose. Furthermore, GK activators that enhance the affinity of GK for glucose can lower the threshold of glucose required for stimulating GLP-1 secretion. In particular, knockdown of *GK* or introduction of the catalysis-defective mutants of GK does not lead to a reduction of ATP production. This is likely because other hexokinases (HK1 to HK3) which have with low *K*_m_ values, are present in L cells^116^ and are sufficient for producing ATP. In contrast, pancreatic β cells primarily express GK that can only generate ATP at a high rate when glucose levels are high, due to its high *K*_m_ for glucose.^117^ Additionally, the pancreatic β cells heavily depend on glucose for energy as they also do not express monocarboxylate transporter 1 (MCT1; ref. ^118^ and Figure S7I) and have very low levels of lactate dehydrogenase,^119,120^ which help provide pyruvate and lactate, respectively, as substitutes for glucose to fuel mitochondrial ATP production. These differences in protein profiling may provide an explanation as to why pancreatic β cells generate higher ATP in high glucose. It is interesting to add that the strict requirement for the pancreatic β cells of glucose may safeguard against over secretion of insulin that may cause hypoglycemia, in case GLP-1 is overproduced by other stimuli such as amino acids.

We have also demonstrated that on binding to glucose, GK directly inhibits K_ATP_ channels, a critical step before the release of GLP-1. We found that GK bound with glucose, occupied by the nonmetabolizable glucose analog, or constitutively bound with glucose as in the catalysis-defective mutant L309P, enhances its interaction with the Kir6.2 subunit of K_ATP_ channel, via the region of aa 205-228. Consistently, the segment aa 205-228 of Kir6.2 is required for GLP-1 release stimulated by glucose. According to the previously resolved structure, this region is distinct from the ATP binding pocket of the channel,^121^ which supports our finding that GK controls K_ATP_ independently of an increase of ATP. Moreover, the inhibition of K_ATP_ by GK can be recapitulated in vitro through inside-out patch clamping in the presence of physiologically relevant concentrations of ATP. These experiments show that the glucose-bound GK reduces the activity (open probability) of K_ATP_ by approximately 40% (Figure 4H), which is also consistent with the percentages observed in whole-cell patch clamping experiments (Figures 4F). Given that the resting membrane potential of L cells in low glucose is primarily governed by K^+^ efflux through K_ATP_,^47,48,122^ such a reduction in K_ATP_ activity can lead to a decrease in membrane potential to around −25 mV, as calculated using the Goldman-Hodgkin-Katz equation.^123^ This membrane potential exceeds the threshold (around −36 mV) necessary to trigger further membrane depolarization through voltage-dependent calcium channels, which ultimately facilitates the exocytosis of vesicles containing GLP-1.^124^ It is also noteworthy that the independence of the increase of ATP in L cells does not imply lack of necessity of basal ATP for the closure of the channel, as it has been previously shown that reduction of ATP levels by mitochondrial inhibitors that grossly inhibit mitochondrial oxidative phosphorylation, prevents the K_ATP_ closure even in high glucose.^9,47^ In fact, our finding that the ATP-binding-defective mutant of K_ATP_ (Kir6.2-K185E/R50G/R201G; ref. ^121^) displays constitutive channel activity and blocks GLP-1 secretion in L cells, which cannot be overridden by the constitutively-glucose-bound GK-L309P mutant (Figure S7J), confirmed a requirement of a priori binding of ATP to K_ATP_.

Our study cannot rule out other mechanisms that sense glucose and somehow contribute to GLP-1 secretion in vivo. For example, we found that the intestine-specific knockout of *GK*, the intestine-specific knock-in of glucose-binding-defective mutant GK-G80A, or the intestine-specific knock-in of Kir6.2-205-228A that cannot bind GK, all did not completely eliminate glucose-induced GLP-1 secretion (Figures 2I, 3E, and 5L). For example, efferent signals from the vagus nerve have been shown to depolarize L cells, leading to GLP-1 secretion even without direct contact with glucose.^125,126^ This effect is particularly evident in L cells located in the distal gut, such as the ileum and colon,^127–129^ where GLP-1 can be released when the vagus nerve is activated by its efferent fibers in the taste bud^130–132^ or in the proximal gut (such as duodenum and jejunum), which are exposed to glucose.^126^ In addition, as previously shown,^71,72,74^ and as confirmed in this study, SGLT1, by virtue of co-transporting the sodium ion, plays an important role in mediating the secretion of GLP-1 through causing membrane depolarization when glucose is present in low concentrations. It is also interesting to note that secretion of GLP-1 can be stimulated by additional stimuli, such as fat and proteins. Interestingly, such secreted GLP-1 does not cause hypoglycemia, for the secretion of insulin depends on a priori presence of high glucose. Such a scenario also applies to metformin and lithocholic acid that both effectively increase GLP-1 secretion do not cause hypoglycemia.^133,134^ The distinctive mechanisms for the secretion of the two hypoglycemic hormones provide a safeguard against hypoglycemia in cases of GLP-1 secretion stimulated by non-carbohydrate stimuli. Finally, our newly uncovered glucose-sensing role of GK in GLP-1 secretion might offer a new avenue for developing drugs that specifically stimulate the secretion of native GLP-1 with potentially less adverse effect,^135^ which is especially important for patients who prioritize blood glucose control over weight loss.

### Limitations of the study

To analyze GLP-1 release in L cells, in vitro culture of cells is required. However, these cells in culture may have lost cell polarity compared to their in vivo state, which may affect their physiological and biophysical behaviors. Nevertheless, many studies have shown that the glucose-induced GLP-1 secretion in cultured primary cells share characteristics of cells with polarity maintained (such as those examined using the intestine ex vivo;^8,78^ this was also performed by us, as shown in Figures S1F and S1G). This suggests that the loss of polarity may not have a significant impact on the properties of L cells. It is reassuring that results on GLP-1 secretion obtained with cultured cells are re-capitulated in vivo in various genetic models. In addition, in this study, we analyzed the glucose-stimulated secretion of GLP-1 specifically in the lumen of the proximal gut, because the refeeding regimen we used allows glucose to accumulate only in the proximal gut (Figure 1A; also demonstrated by others; ref. ^86,87^). As the primary L cells we used were derived from whole intestinal tissues and thus included L cells from both proximal and distal regions (Figure 1F), it is reasonable to suggest that the distal intestines may also not have any increase in ATP in high glucose.

## Supporting information

Full scan of western blot

Soource data

## RESOURCE AVAILABILITY

### Lead contact

Further information and requests for resources and reagents should be directed to and will be fulfilled by the lead contact, Sheng-Cai Lin.

### Materials availability

All reagents, cell lines and mouse strains generated in this study are available from the lead contact upon request.

### Data and code availability

- The datasets generated and analyzed during the current study are contained within the manuscript and supplemental figures.
- The MS proteomics data have been deposited to the ProteomeXchange Consortium (http://proteomecentral.proteomexchange.org) through the iProX partner repository^136,137^ with the dataset identifier IPX0011750000.
- Full immunoblots are provided as a “Full scans” file.
- Raw data and statistical analysis data for all graphs are provided with this paper as a “Source data” file.
- This paper does not report original code.
- Any additional information required to reanalyze the data reported in this paper is available from the lead contact upon request.

## ACKNOWLEDGEMENTS

We thank Dr. Lei Chen and Tianyi Hou (Peking University) for sharing insights into the structure and regulation of K_ATP_, along with the expression plasmids for K_ATP_; Ximin Chi (Xiamen University) for designing the domain mapping strategy according to the K_ATP_ structure; Changchuan Xie, Yaying Wu, and Zheni Xu of the core facility center of Xiamen University for the technical support with protein mass spectrometry; Qingfeng Liu (Xiamen University) for the assistance with the measurement of local intracellular ATP:ADP ratios by spinning disk confocal microscopy; Changchun Xiao (Sanofi China) for the *Rosa26*-CTV vector; Mingliang Zhang (Shanghai Jiao Tong University Affiliated Sixth People’s Hospital) for the instructions on the intestinal glucose measurement; Yunxia Zhu (Nanjing medical University) for the instructions on the culture of pancreatic β cells and measuring insulin secretion from them; and Su-Qin Wu and Ying He (Xiamen University) for the mouse in vitro fertilization; Zhiyun Ye, Wei Hong and Tian-Yu Tang (Xiamen University) for secretarial assistance; and all the other members of the S.-C.L. laboratory for the technical assistance. The artworks shown in Figures 4C (https://BioRender.com/xer61y9), 5M (https://BioRender.com/xer61y9), and S2F (https://BioRender.com/c7p1hxa) were created in BioRender. Pi, h. (2025). In addition, the artworks shown in Figures 4D, 4E (https://BioRender.com/xer61y9), and 4H (https://BioRender.com/xer61y9) were also created in BioRender. Pi, h. (2025), and include modified elements created by Servier Medical Art (https://smart.servier.com/, which are licensed under a Creative Commons Attribution 3.0 Unported Licence). This work was supported by grants from the National Natural Science Foundation of China (#82088102, #32471223 and #32200633), the National Key R&D Program of China (2022YFA0806501), the Scientific Research Innovation Capability Support Project for Young Faculty (ZYGXONJSKYCXNLZCXM-B5), the Joint Funds for the Innovation of Science and Technology, Fujian province (2021Y9232, 2021Y9227, 2023Y9448), the Fujian provincial health technology project (2022ZD01005, 2022ZQNZD009), the Special Research Funds for Local Science and Technology Development Guided by Central Government (2023L3020), the XMU-Fujian Cancer Hospital cooperation grant for the Center for Metabolism Research, and the Provincial College Students Innovation and Entrepreneurship Program (#S202410384385).

## AUTHOR CONTRIBUTIONS

C.L., X.T., C.-S.Z., and S.-C.L. conceived the study and designed the experiments. C.L. and X.T. explored the roles of GK, as well as the requirement of ATP, in regulating GLP-1 secretion in mice and cells. M.L. determined the regulation of K_ATP_ channel by GK, with assistance from S.L. and X.Y.; these experiments were performed in the laboratory of H.X., under the supervision of M.H. and H.X. H.-Y.Y. evaluated the enzymatic activity of GK and its mutants, as well as the binding affinity of GK towards glucose analogs, with the help of J.C. The binding domain of K_ATP_ to GK was determined by C.L. and H.-Y.Y., with assistance from W.M. GK mutations among MODY2 patients were screened, and the related patient experiments were carried out by L.Z. and X.L. Knock-in mice with GK mutants were generated by J.W. C.Z. measured the levels of adenylates and glycolytic intermediates using HPLC-MS and CE-MS. X.S. isolated primary L cells using flow cytometry. Z.-Z.Z. profiled protein expression in L cells through quantitative protein mass spectrometry. C.Y., J.L. and K.W. generated mice with L cells expressing GFP. X.H. helped the culture of primary intestinal cells. T.M. and Z.Q. guided the intestinal enterostomy procedures in mice. C.J. provided technical assistance for GLP-1 measurements. C.-S.Z. and S.-C.L. wrote the manuscript.

## SUPPLEMENTAL INFORMATION

Supplemental information can be found online at https://doi.org/10.1016/j.cell.

## STAR METHODS

### KEY RESOURCES TABLE

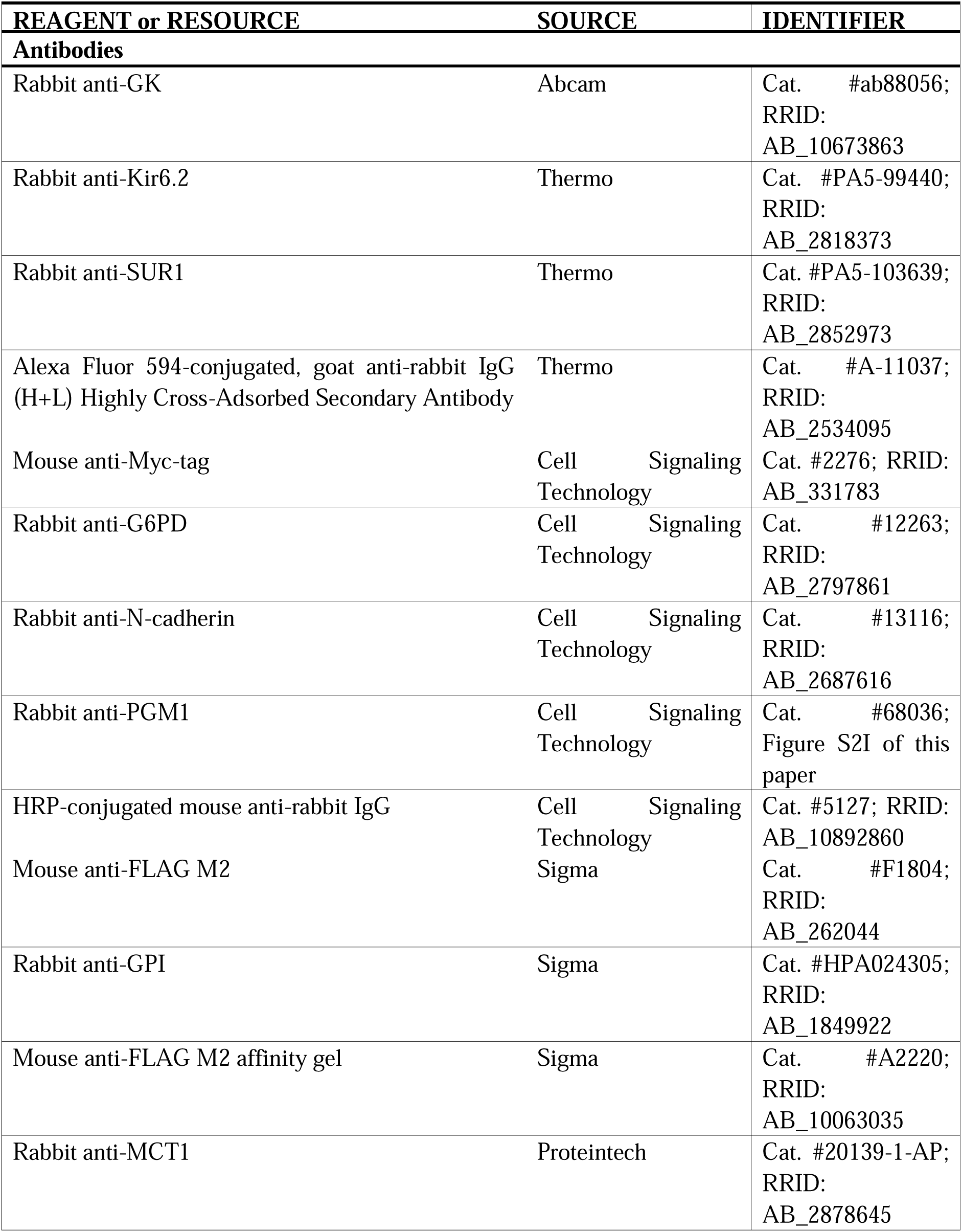
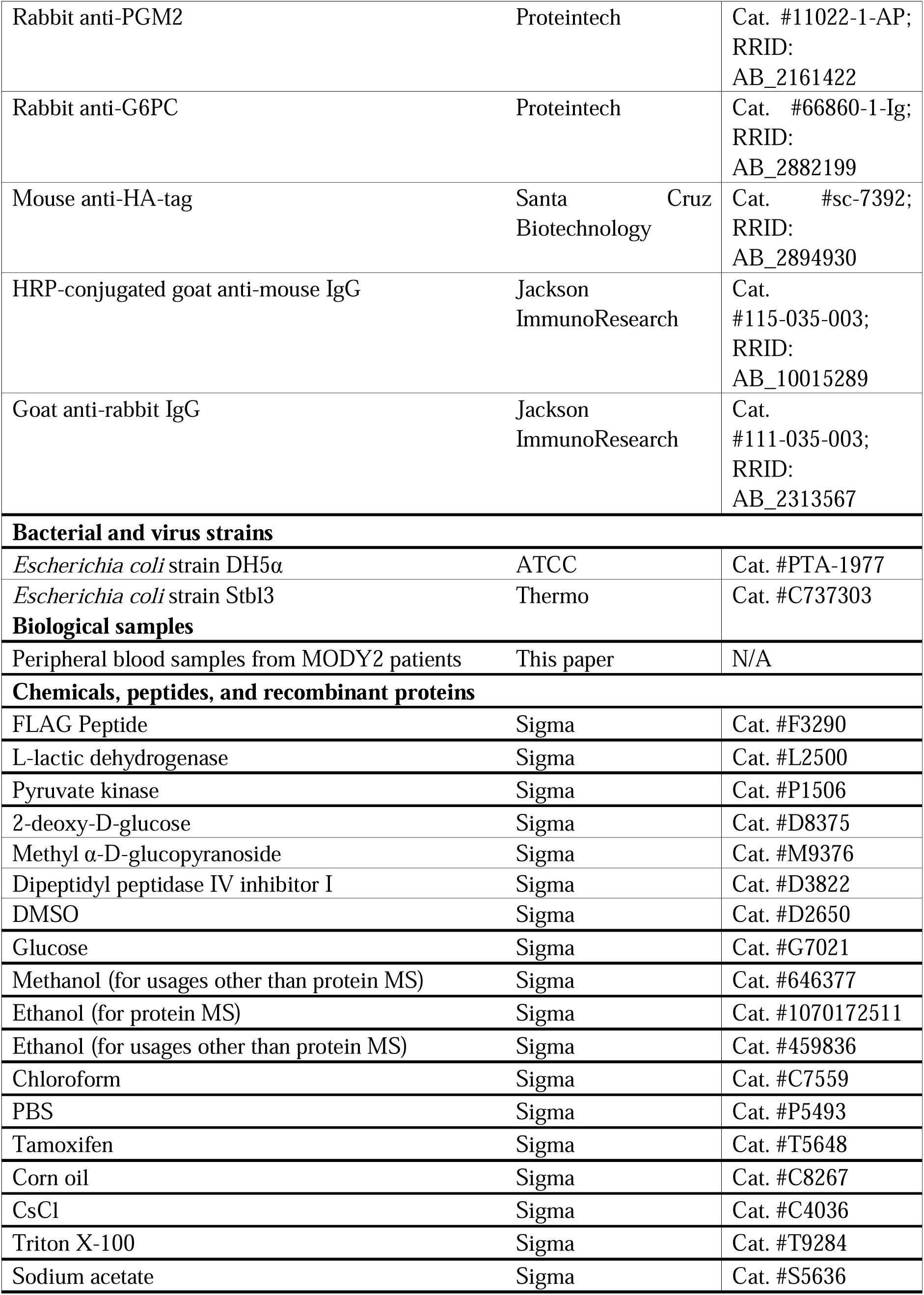
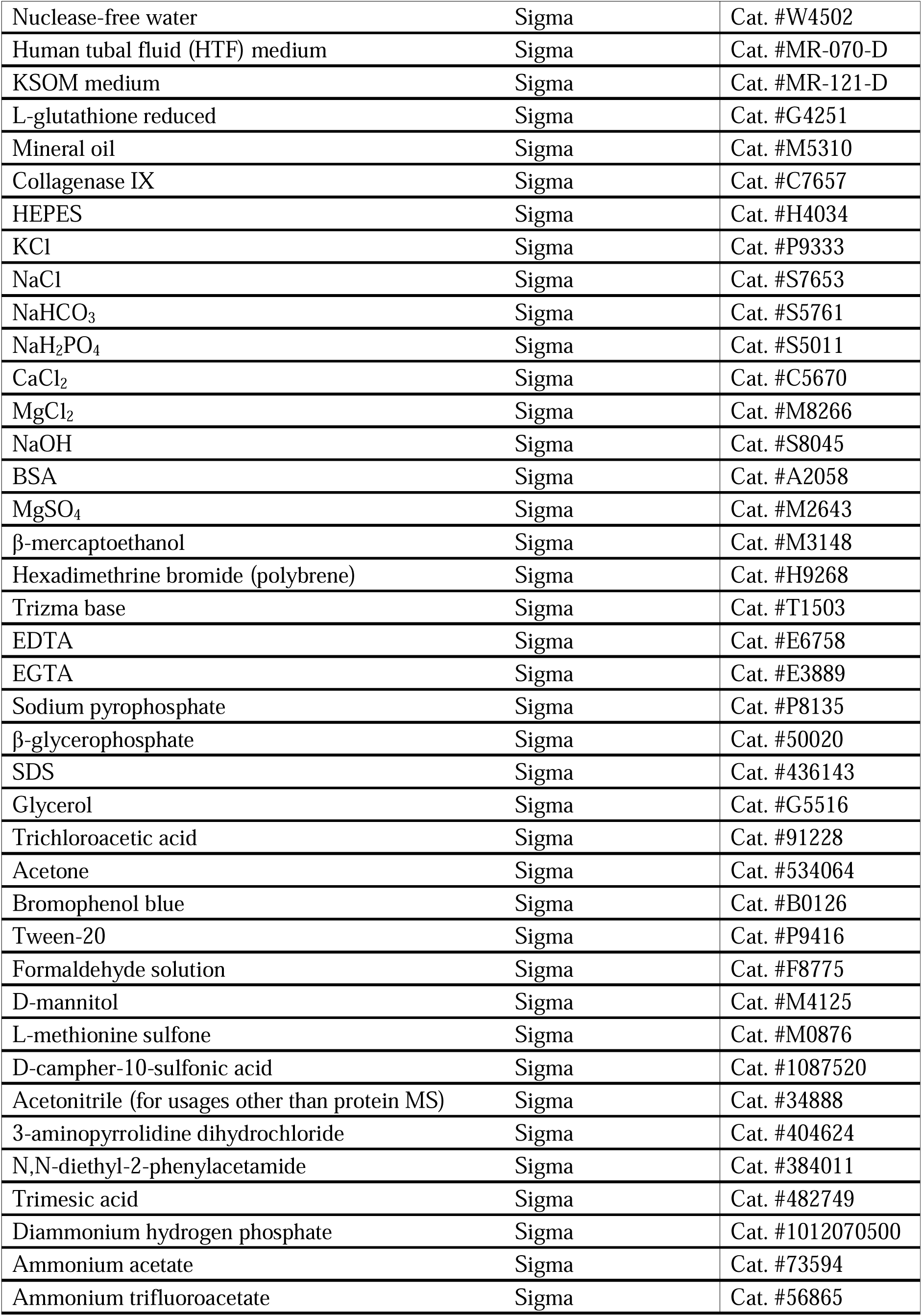
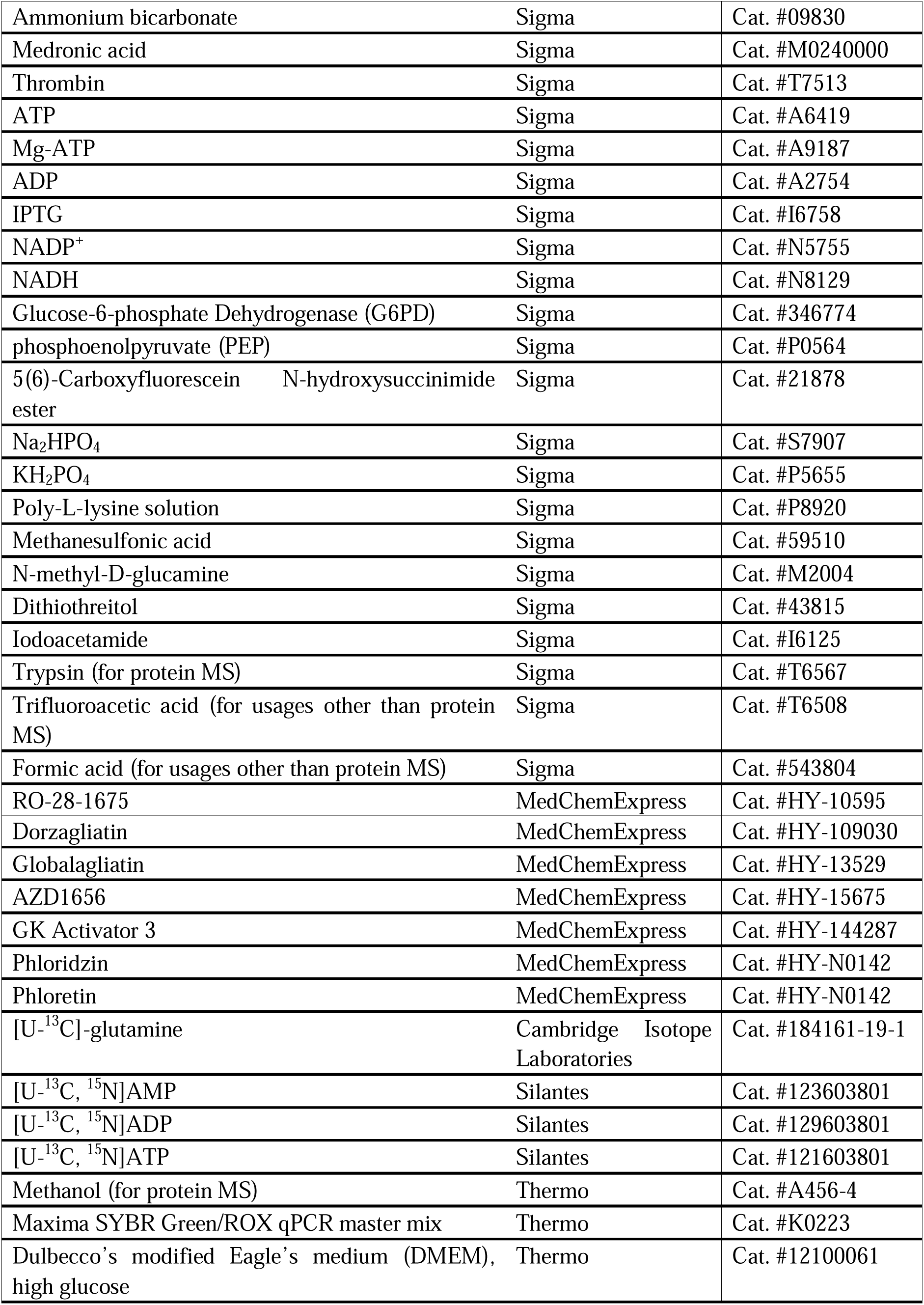
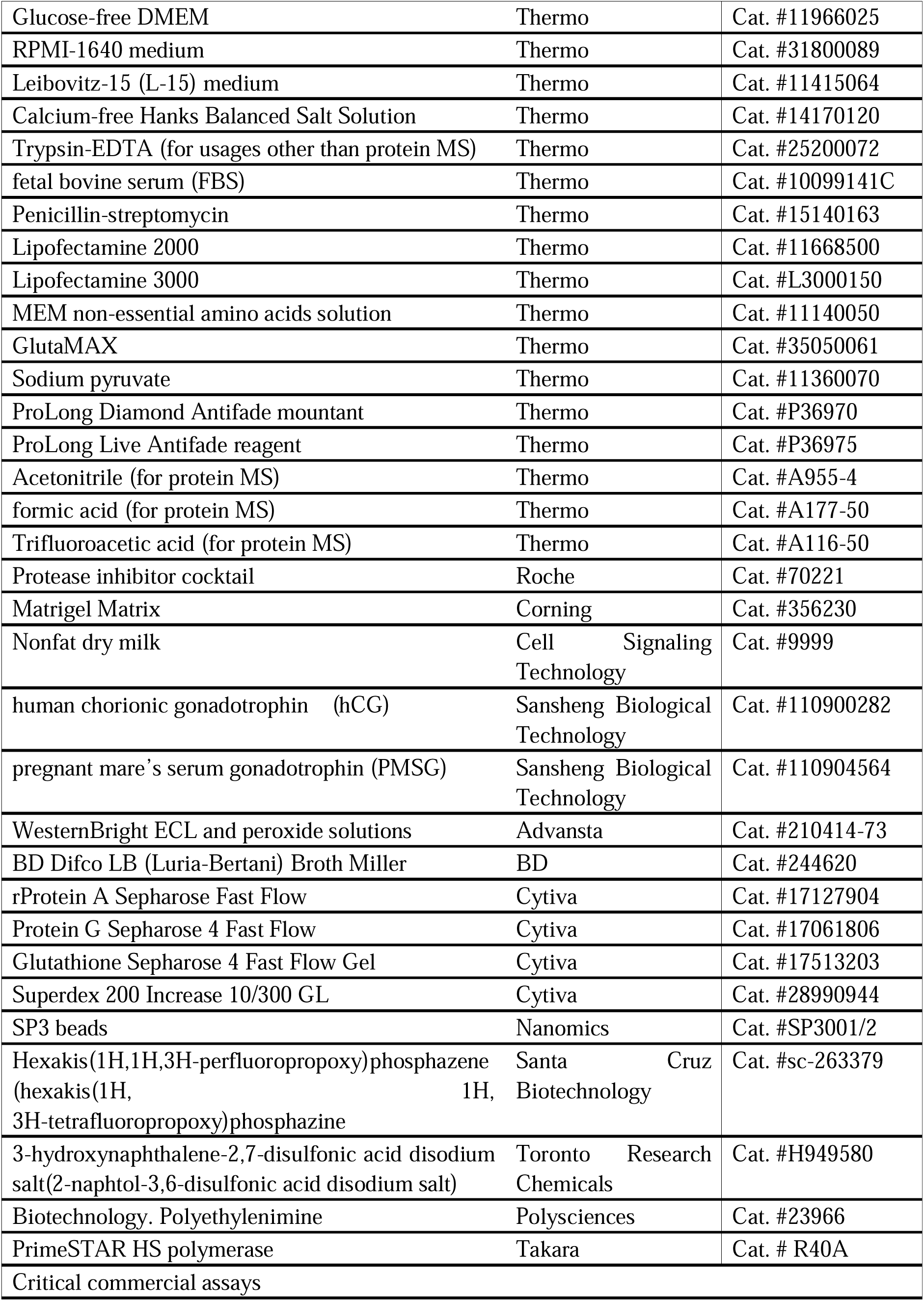
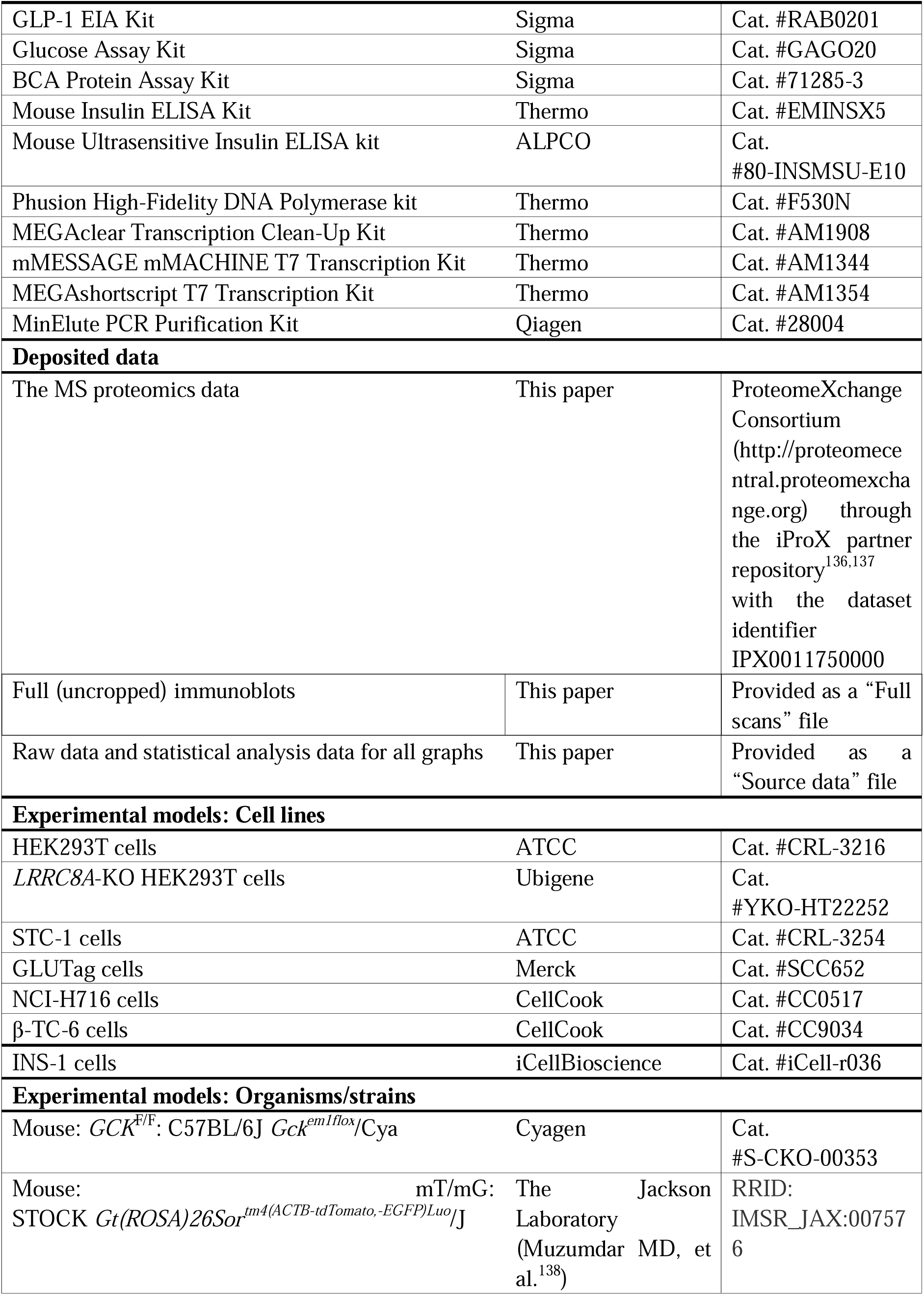
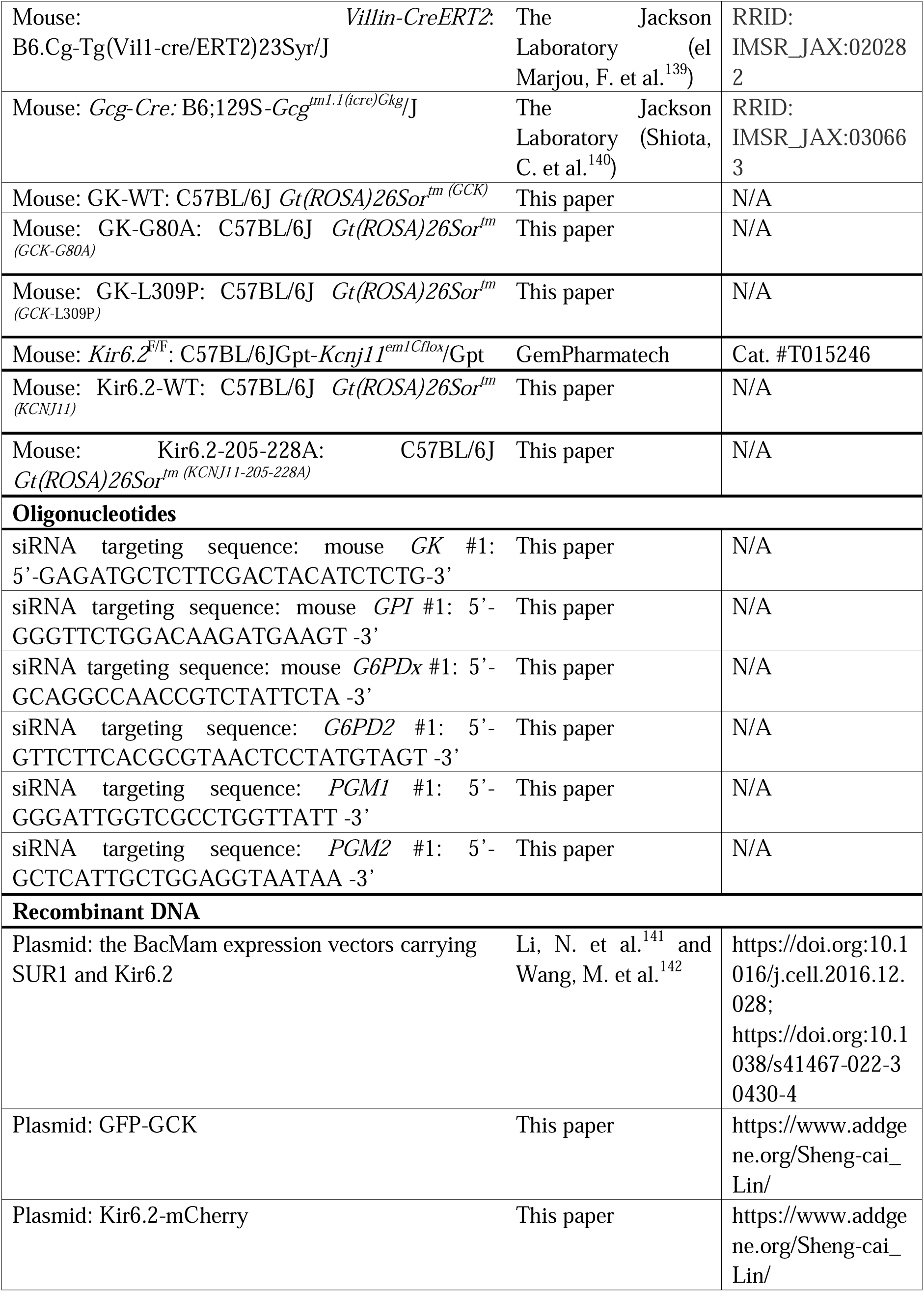
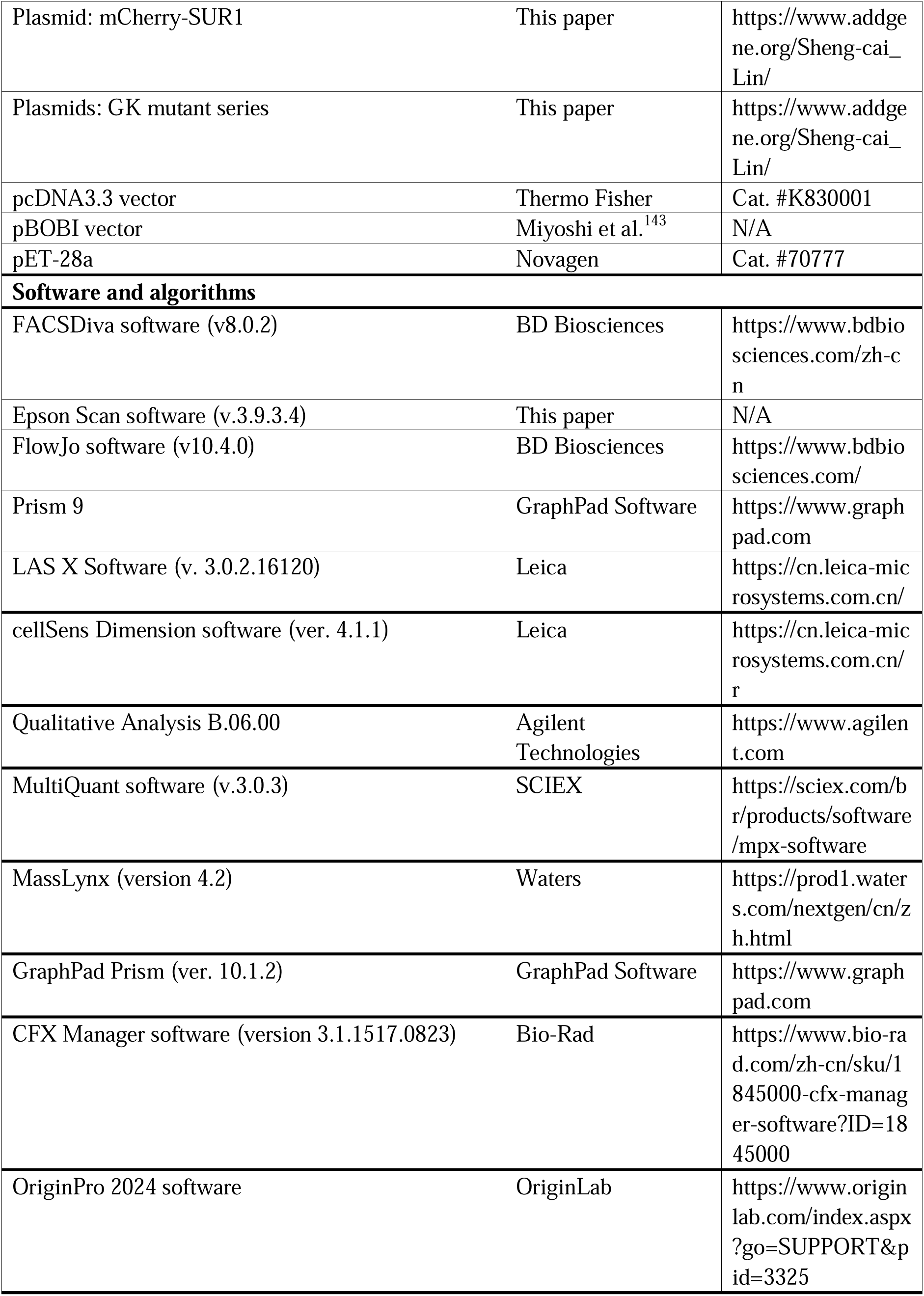
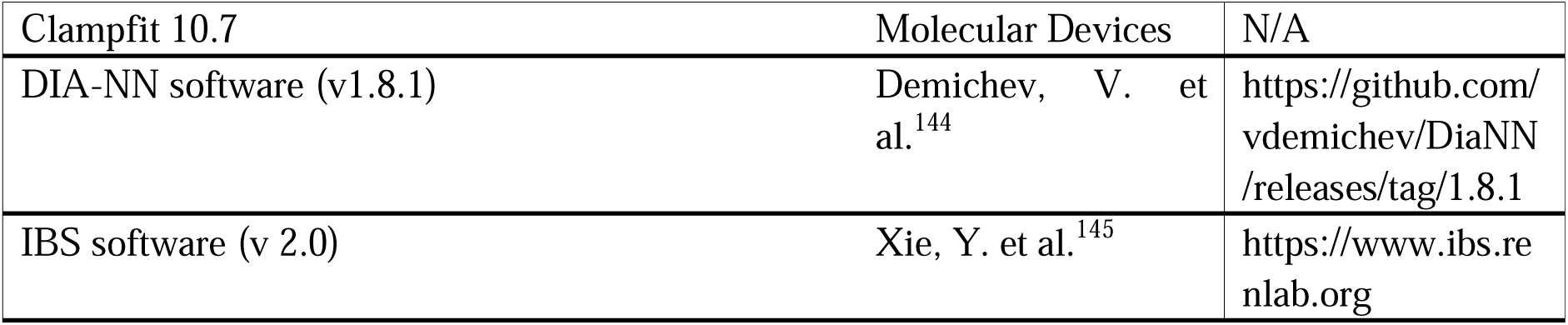

### EXPERIMENTAL MODEL AND STUDY PARTICIPANT DETAILS

#### Mouse strains

Protocols for all rodent experiments were approved by the Institutional Animal Care and the Animal Committee of Xiamen University (XMULAC20180028 and XMULAC20220050). Unless stated otherwise, mice were housed with free access to water and standard diet (65% carbohydrate, 11% fat, 24% protein) under specific pathogen-free conditions. The light was on from 8:00 to 20:00, with the temperature kept at 21-24 °C and humidity at 40-70%. Only male mice were used in the study, and male littermate controls were used throughout the study.

Mice of wildtype C57BL/6J (#000664), *Gcg*-*Cre* (#030663; provided by Dr. Klaus Kaestner^140^), *Villin*-*CreERT2* (#020282; provided by Dr. Sylvie Robine^139^), and mT (membrane-targeted tdTomato)/mG (membrane-targeted EGFP) (#007576; provided by Dr. Liqun Luo^138^) were obtained from The Jackson Laboratory. *GCK*^F/F^ (#S-CKO-00353) mice were purchased from Cyagen, and *Kir6.2*^F/F^ (*Kcnj11*^F/F^; #T015246) from GemPharmatech.

Mice with intestine-specific *GCK* knockout were generated by crossing *GCK*^F/F^ mice with *Villin*-*CreERT2* mice, followed by intraperitoneal injection with tamoxifen (dissolved in corn oil) at 200 mg/kg, 4 times a week for a week. The knockout offspring were validated through genotyping. For genotyping the *GCK* locus, the following programs were used: pre-denaturing at 95 °C for 300 sec; denaturing at 95 °C for 30 sec, annealing at 65 °C for 30 sec, followed by extension at 72 °C for 30 sec, repeated for 10 cycles, with the annealing temperature decreasing by 0.5 °C in each cycle; denaturing at 95 °C for 30 sec, annealing at 60 °C for 30 sec, then extending at 72 °C for 30 sec in each cycle for an additional 28 cycles; and final extending at 72 °C for 10 min. For genotyping *Villin*-*CreERT2*, the following programs were used: pre-denaturing at 95 °C for 300 sec; denaturing at 95 °C for 30 sec, annealing at 65 °C for 30 sec, followed by extension at 72 °C for 90 sec, repeated for 10 cycles, with the annealing temperature decreasing by 0.5 °C in each cycle; denaturing at 95 °C for 30 sec, annealing at 60 °C for 30 sec, then extending at 72 °C for 90 sec in each cycle for an additional 28 cycles; and final extending at 72 °C for 10 min. The following primers: 5’-AACATGAAGCTAGGGCCAGCGCT-3’ and 5’-ATCTCCTCATCCCTTTGTCCCTTT-3’ for *LoxP*; 5’-TCAGTGCCTTGTTGGATG-3’, 5’-GGAACCGAGATGATGTAGC-3’, 5’-GTGTGGGACAGAGAACAAACC-3’, and 5’-ACATCTTCAGGTTCTGCGGG-3’ for *Villin-CreERT2* were used.

Wildtype GK, along with G80A and L309P mutants, was introduced to the intestines of *GK*^F/F^ mice through the *Rosa26*-*LSL*(*LoxP*-*Stop*-*LoxP*) system as described previously,^146,147^ followed by crossing with the *Villin*-*CreERT2* mice. The removal of the intestinal *GK* gene, and the *LSL* cassette ahead of the introduced GK and its mutants (to trigger the expression of introduced GK), was then achieved by intraperitoneal injection of tamoxifen. To introduce GK or its mutants into *GK*^F/F^ mice, cDNA fragments encoding GK, GK-G80A, or GK-L309P were inserted into the *Rosa26*-CTV vector,^148^ followed by purification of the plasmids using CsCl density gradient ultracentrifugation method. Some 100 μg of plasmid was then diluted with 500 μL of di-distilled water, followed by concentrating via centrifugation at 14,000 g at room temperature in a 30-kDa-cutoff filter (UFC503096, Millipore) to 50 μL of solution. The solution was diluted with 450 μL of di-distilled water, followed by another two rounds of dilution/concentration cycles. The plasmid was then mixed with 50 μL of di-distilled water to a final volume of 100 μL, followed by mixing with 10 μL of NaAc solution (3 M stock concentration, pH 5.2). The mixture was then mixed with 275 μL of ethanol, followed by incubating at room temperature for 30 min to precipitate the plasmid. The precipitated plasmid was collected by centrifugation at 16,000 g for 10 min at room temperature, followed by washing with 800 μL of 75% (v/v) ethanol (in di-distilled water) twice. After being dissolved in 100 μL of nuclease-free water, the plasmid, along with SpCas9 mRNA and the sgRNAs against the mouse *Rosa26* locus, was then microinjected into the in vitro fertilized (IVF) embryos of the *GK*^F/F^ mice. To generate the SpCas9 mRNA, 1 ng of pcDNA3.3-hCas9 plasmid (constructed by inserting the Cas9 fragment released from Addgene #41815 (ref. ^149^), into the pcDNA3.3 vector; diluted to 1 ng/μL) was amplified using the Phusion High-Fidelity DNA Polymerase kit on a thermocycler (T100, Bio-Rad) with the following programs: pre-denaturing at 98 °C for 30 sec; denaturing at 98 °C for 10 sec, annealing at 68 °C for 25 sec, then extending at 72 °C for 2 min in each cycle; and final extending at 72 °C for 2 min; cycle number: 33. The following primer pairs were used: 5’-CACCGACTGAGCTCCTTAAG-3’, and 5’-TAGTCAAGCTTCCATGGCTCGA-3’. The PCR product was then purified using the MinElute PCR Purification Kit following the manufacturer’s instructions. The purified SpCas9 PCR product was then subjected to in vitro transcription using the mMESSAGE mMACHINE T7 Transcription Kit following the manufacturer’s instructions (with minor modifications). Briefly, 5.5 μL (300 ng/μL) of SpCas9 PCR product as the template was mixed with 10 μL of 2× NTP/ARCA solution, 2 μL of 10× T7 Reaction Buffer, 0.5 μL of RNase inhibitor, 2 μL of T7 Enzyme Mix, and 4.5 μL of nuclease-free water, followed by incubating at 37 °C for 2 h. The mixture was then mixed with 1 μL of Turbo DNase, followed by incubating at 37 °C for 20 min to digest the template. The mixture was then mixed with 20 μL of 5× *E*-PAP Buffer, 10 μL of 25 mM MnCl_2_, 10 μL of 10 mM ATP, 4 μL of *E*-PAP enzyme, and 36 μL of nuclease-free water, followed by incubating at 37 °C for 20 min for poly(A) tailing. The tailed product was then purified using the MEGAclear Transcription Clean-Up Kit following the manufacturer’s instructions (with minor modifications). Briefly, 20 μL of tailed RNA was mixed with 20 μL of Elution Solution, followed by mixing with 350 μL of Binding Solution Concentrate. Some 250 μL of ethanol was then added to the mixture, followed by passing the mixture through the Filter Cartridge and washing with 250 μL of Wash Solution twice. The RNA was then eluted with 50 μL of pre-warmed (at 90 °C) Elution Solution. The sgRNAs were prepared as in the SpCas9 mRNA preparation, except that: a) the gRNA Cloning Vector (Addgene, #41824, ref. ^149^) was used as the template, and the following programs: pre-denaturing at 98 °C for 30 sec; denaturing at 98 °C for 10 sec, annealing at 60 °C for 25 sec, then extending at 72 °C for 20 sec in each cycle; and final extending at 72 °C for 2 min; cycle number: 33; and the following primers: 5’-GAAATTAATACGACTCACTATAGGCGCCCATCTTCTAGAAAGACGTTTTA GAGCTAGAAATAGC-3’, and 5’-AAAAGCACCGACTCGGTGCC-3’; were used; b) in vitro transcription was performed using the MEGAshortscript T7 Transcription Kit, in which the mixture containing: 7.5 μL (100 ng/μL) of purified PCR product, 2 μL of T7 10× T7 Reaction Buffer, 2 μL of T7 ATP solution, 2 μL of T7 CTP solution, 2 μL of T7 GTP solution, 2 μL of T7 UTP solution, 0.5 μL of RNase inhibitor, 2 μL of T7 Enzyme Mix, and 7.5 μL of nuclease-free water; was prepared. In addition, the poly(A) tailing assay was not performed.

The prepared *Rosa26*-CTV-GK, SpCas9 mRNA and *Rosa26* sgRNA plasmids were then microinjected into each of the zygotes of *GK*^F/F^ mice. To prepare the zygotes, the *GK*^F/F^ mice were first subjected to IVF (ref. ^150^, with minor modifications^151^). Briefly, the 4-week-old *GK*^F/F^ female mice were intraperitoneally injected with pregnant mare’s serum gonadotrophin (PMSG) at a dose of 10 U/mouse. At 46 h after the PMSG injection, 10 U/mouse human chorionic gonadotrophin (hCG) was intraperitoneally injected. At 12 h after the hCG injection, oocytes from the oviducts of female mice, along with sperms from cauda epididymides and vasa deferentia of 16-week-old, proven stud *GK*^F/F^ male mice, were isolated. To isolate oocytes, oviducts were briefly left on a filter paper, followed by incubating in a human tubal fluid medium (HTF)/GSH drop on an IVF dish (prepared by placing 200 μL of HTF solution supplemented with 125 mM GSH on a 35-mm dish to form a drop, followed by covering the drop with mineral oil and pre-balancing in a humidified incubator containing 5% CO_2_ at 37 °C for 0.5 h before use). The ampulla was then torn down by forceps, and the cumulus oocyte masses inside were collected and transferred to another HTF/GSH drop. To isolate sperms, cauda epididymides and vasa deferentia were briefly left on a filter paper, followed by penetrating with a 26 G needle on the cauda epididymides 5 times. Sperms were then released to an HTF drop on sperm capacitation dish (prepared by placing 200 μL of HTF solution on a 35-mm dish to form a drop, followed by covering the drop with mineral oil and pre-balancing in a humidified incubator containing 5% CO_2_ at 37 °C for 12 h before use) by slightly pressing/squeezing the cauda epididymides, followed by incubation in a humidified incubator containing 5% CO_2_ at 37 °C for 0.5 h. The capacitated, motile sperms (located on the edge of each HTF drop) were then collected, followed by adding to the oocyte masses soaked in the HTF/GSH drop, 8 μL per drop. The IVF dishes containing oocyte masses and sperms were then cultured in a humidified incubator containing 5% CO_2_ at 37 °C for 4 h, followed by collecting and washing oocytes in a KSOM drop (freshly prepared by placing 20 μL of KSOM medium on a 35-mm dish to form a drop, followed by covering the drop with mineral oil and pre-balancing in a humidified incubator containing 5% CO_2_ at 37 °C for 0.5 h) twice. The oocytes were then cultured in an HTF/GSH drop on an IVF dish for another 12 h in a humidified incubator containing 5% CO_2_ at 37 °C. The presumptive zygotes (in which 2 pronuclei and an extruded, second polar body could be observed) were then picked up. Some 10 pl of DNA mixture containing *Rosa26*-CTV-GK plasmid (20 ng/μL final concentration), SpCas9 mRNA (120 ng/μL final concentration), and *Rosa26* sgRNA (100 ng/μL), was microinjected into each of the zygotes and were cultured in KSOM medium at 37 °C in a humidified incubator containing 5% CO_2_ for 16 h. The zygotes/embryos at the two-cell stage were picked up and transplanted into pseudopregnant ICR female mice (8-10 weeks old, >26 g; prepared by breeding the in-estrus female with a 14-week-old, vasectomized male at a day before the transplantation), 20 zygotes/embryos per mouse, and the offspring carrying the *LSL*-*GK* allele were further outcrossed 6 times to C57BL/6 mice before crossing with the *Villin-CreERT2* mice. Mice were validated by genotyping. For genotyping the *Rosa26* locus, the following programs were used: pre-denaturing at 98 °C for 300 sec; denaturing at 95 °C for 30 sec, annealing at 64 °C for 30 sec, then extending at 72 °C for 45 sec in each cycle for 5 cycles; denaturing at 95 °C for 30 sec, annealing at 61 °C for 30 sec, then extending at 72 °C for 45 sec in each cycle for 5 cycles; denaturing at 95 °C for 30 sec, annealing at 58 °C for 30 sec, then extending at 72 °C for 45 sec in each cycle for 5 cycles; denaturing at 95 °C for 30 sec, annealing at 55 °C for 30 sec, then extending at 72 °C for 45 sec in each cycle for 5 cycles; and final extending at 72 °C for 10 min. For genotyping other genes and elements, the following programs were used: pre-denaturing at 95 °C for 300 sec; denaturing at 95 °C for 30 sec, annealing at 58 °C for 30 sec, then extending at 72 °C for 30 sec in each cycle; and final extending at 72 °C for 10 min; cycle number: 35. The following primers: 5’-AGAGAATTCGGATCCATGCTGGACGACAGAGCCAG-3’ and 5’-CTTCCATGGCTCGAGCTTATCGATGTCTTCGTGCCT-3’ for generating PCR products for sequencing GK-WT and GK-G80A; 5’-AGAGAATTCGGATCCTGTGCGTCAATACCGAGTGG-3’ and 5’-CTTCCATGGCTCGAGGCGTGCTCAGGATGTTGTAG-3’ for generating PCR products for sequencing GK-L309P.

Mice with intestine-specific *Kir6.2* knockout were generated using the same method as that for the intestine-specific *GCK* knockout mice. Geneotyping was performed in a similar manner to the *GCK* knockout mice, except that the following primers: 5’-TATGCACAGGCCAAGCAAAC-3’ and 5’-ATCGTGGACCCTGCTACAAAG-3’, were used for genotyping the *Kir6.2* locus. The PCR consisted of pre-denaturing at 95 °C for 300 sec; denaturing at 95 °C for 30 sec, annealing at 65 °C for 30 sec, followed by extension at 72 °C for 30 sec, repeated for 10 cycles, with the annealing temperature decreasing by 0.5 °C in each cycle; denaturing at 95 °C for 30 sec, annealing at 60 °C for 30 sec, then extending at 72 °C for 30 sec in each cycle for an additional 28 cycles; and final extending at 72 °C for 10 min; were used to genotype *Kir6.2* locus. Wildtype Kir6.2 and 205-228a mutants were introduced into the intestines of *Kir6.2*^F/F^ mice through the *Rosa26*-*LSL* system, similar to the method used for the *GCK* mutants. This was followed by crossing with the *Villin*-*CreERT2* mice. The primers used for generating PCR products for sequencing the Kir6.2 and its mutant were: 5’- AGAGAATTCGGATCCTGGGGCTCATGATCAACGC-3’ and 5’- CTTCCATGGCTCGAGATGATCTCGAGGTCCTGGTGG −3’.

Transgenic mice with L cells expressing GFP were generated and validated as described previously.^88^ Briefly, mT/mG mice, which possess *LoxP* sites flanking an mT cassette (driven under the *Actin* promoter, resulting in red fluorescence in all tissues and cell types), were crossed with *Gcg*-*Cre* mice (Cre expression under the glucagon (*Gcg*) or preproglucagon promoter). This leads to removal of the mT cassette, allowing the expression of the downstream mG cassette specifically in L cells (as well as pancreatic α cells, although only intestinal tissues were analyzed in this study). For genotyping the mT/mG mice, the following programs were used: pre-denaturing at 94 °C for 180 sec; denaturing at 94 °C for 30 sec, annealing at 61 °C for 60 sec, then extending at 72 °C for 60 sec in each cycle for 35 cycles; and final extending at 72 °C for 2 min. For genotyping the *Gcg*-*Cre* mice, the following programs were used: pre-denaturation at 95 °C for 300 sec; denaturation at 95 °C for 30 sec, annealing at 65 °C for 30 sec, followed by the extension step at 68 °C for 30 sec, repeated for 10 cycles, with the annealing temperature decreasing by 0.5 °C in each cycle; denaturation at 94 °C for 30 sec, annealing at 60 °C for 30 sec, then extension at 72 °C for 30 sec in each cycle for an additional 28 cycles; and final extension at 72 °C for 10 min. The following primers: 5’-CTCTGCTGCCTCCTGGCTTCT-3’, 5’-CGAGGCGGATCACAAGCAATA-3’, and TCAATGGGCGGGGGTCGTT for mT/mG; 5’-CATGGCATTGGAGCCATAAG-3’, 5’-CTGGGGTTCTCCTCTGTGTC-3’, and 5’-TGTTGGATGGTCTTCACAGCC-3’ for *Gcg*-*Cre* were used.

The following ages of mice were used for the experiments: a) for analyzing glucose levels in the blood and intestinal lumen, as well as circulating GLP-1 and insulin, 6-month-old wildtype mice and GK/Kir6.2 mutant-KI mice were used (with tamoxifen injected at 5 months old); b) for isolating primary intestinal cells, 5-month-old mice were used; c) for other experiments, 4-month-old mice were used.

#### Determination of GLP-1 secretion in patients with GK mutations

Patients were recruited from the Department of Endocrinology at Zhongshan Hospital, Fudan University. Mutations in the GK gene were identified using Illumina-based whole-exome sequencing (WES) using genomic DNA isolated from EDTA-anticoagulated peripheral blood samples, achieving a mean coverage depth of over 50× and a target region sensitivity of 98%. All candidate variants were subsequently validated through Sanger sequencing. Two female patients diagnosed with GCK-MODY, aged 42 and 22 years, were included in the study; they carried GK-R397L and GK^Δ^^92-CT^ mutations, respectively. Written informed consents were obtained from each participant, and the study protocol was approved by the Institutional Review Board of Zhongshan Hospital, Fudan University, in accordance with the principles outlined in the Declaration of Helsinki and the International Conference on Harmonization Good Clinical Practice Guidelines.

Before the assessment, participants discontinued all anti-diabetes medications for at least two weeks. They fasted overnight, beginning at 20:00 the night before the assessments. At 08:00 the following day, they took glucose orally at a dosage of 1 g/kg, dissolved in a total volume of 250 mL of water. Insulin, GLP-1, and C-peptide were determined as described previously.^152^ Briefly, blood samples were collected via intravenous catheters from the antecubital vein at 0, 30, and 120 min post-glucose administration. Blood was drawn into ice-cold K_2_EDTA spray-coated tubes (cat. 366420, BD P800 Blood Collection System) containing 6 μL of diprotin A (5 mg/mL in DMSO) and into SST II Advance Tubes (cat. 367954, BD Vacutainer), with approximately 5 mL collected for each time point. The blood samples were centrifuged at 3,000 g for 10 min at 4 °C, followed by plasma glucose measurement using the glucose oxidase method and GLP-1 measurement using an electrochemiluminescence immunoassay, both performed at KingMed Diagnostics Co., Ltd. (Shanghai, China). The blood collected in SST II Advance Tubes was similarly centrifuged at 3,000 g for 10 min at 4 °C. After centrifugation, serum insulin and C-peptide concentrations were determined using a two-site electrochemiluminescence immunoassay, conducted on the cobas e 801 analytical unit (Roche Modular Analytics) at the Department of Endocrinology at Zhongshan Hospital.

### METHODS DETAILS

#### Data reporting

The chosen sample sizes conform to those used in this field: n = 6-8 samples for determining the levels of GLP-1, insulin and glucose in mice;^133,134,153,154^ n = 4-12 samples for the secretion of GLP-1 in L cells and intestine tissues;^155,156^ n = 3-5 samples to evaluate the levels of metabolites in cells^157,158^ and tissues;^157–159^ n = 9-10 cells to determine the local intracellular ATP:ADP ratios;^94,95,160^ n = 3 replicates to determine GK activity and the binding affinity of GK towards glucose analogs;^161,162^ and n = 2-3 samples to determine the expression levels of a specific protein.^163^ No statistical methods were used to predetermine the sample size. All experimental findings were repeated as stated in the figure legends, and all additional replication attempts were successful. For animal experiments, mice were housed under the same conditions or in the same place. For cell experiments, cells of each genotype were cultured in the same CO_2_ incubator and were seeded in parallel. Each experiment was designed and performed along with proper controls, and samples for comparison were collected and analyzed under the same conditions. Randomization was applied wherever possible. For example, during MS analyses (for metabolites and proteins), samples were processed and subjected to the MS in random orders. In cell experiments, cells of each genotype were parallel seeded and randomly assigned to different treatments. Otherwise, randomization was not performed. For example, when performing immunoblotting, samples needed to be loaded in a specific order to generate the final figures. Blinding was applied wherever possible. For example, samples, cages, or dishes during sample collection and processing were labelled as code names that were later revealed by the individual who picked and treated animals or cells, but did not participate in sample collection and processing, until assessing the outcome. Similarly, during microscopy data collection and statistical analyses, the fields of view were chosen on a random basis, and are often performed by different operators, preventing potentially biased selection for desired conditions. Otherwise, blinding was not performed, such as the measurement of GK activity in vitro, as different reagents were added for particular reactions.

#### Isolation of primary intestinal cells

Primary intestinal epithelial cells were isolated from mouse intestine as described previously,^48^ with minor modifications. In brief, mice were sacrificed through cervical dislocation, and the small intestinal tissues were quickly excised. Tissues were flushed with ice-cold Leibovitz-15 (L-15) medium to remove luminal contents, then divided in half, cut open longitudinally, and sliced into 1-2 mm segments in a petri dish containing the ice-cold L-15 medium. For each mouse, the tissues were digested at 37 °C in a total of 2 mL of 0.4 mg/mL collagenase XI solution (freshly prepared in calcium-free Hanks Balanced Salt Solution (HBSS) for 30 min on an orbital shaker set to 80 rpm. After digestion, the cells were centrifuged at 300 g at room temperature. The pellets were then resuspended in 10 mL of collagenase XI for an additional 15 min of digestion before being filtered through a 70-μm cell strainer (cat. 350350; BD Falcon) and subjected to another round of centrifugation at 300 g at room temperature. The resulting cell pellets were resuspended in Dulbecco’s modified Eagle’s medium (DMEM) containing 25 mM glucose, and rinsed twice with DMEM. Cells were then suspended in DMEM supplemented with 10% (v/v) fetal bovine serum (FBS), 2 mM L-glutamine, 100 IU of penicillin, and 100 μg/mL of streptomycin. The cells were then seeded into a Matrigel-coated 24-well dish. To prepare the Matrigel-coated dish, Matrigel was thawed at 4 °C for 12 h, followed by dilution with ice-cold PBS to a concentration of 3 mg/mL. The diluted Matrigel was then rapidly dispensed to the wells of a pre-chilled 24-well dish at 100 µL per well, followed by a gentle swirl to ensure even coverage. The plates were incubated at 37 °C for 1 h to allow complete gel polymerization, followed by a rinse with PBS under gentle agitation. The cells were then cultured in these dishes for 2 days at 37 °C in a humidified incubator with 5% CO_2_ prior to further experiments.

Primary L cells were isolated as described previously.^88^ Briefly, primary intestinal epithelial cells were digested from mice engineered to express GFP in L cells as described above, except that collagenase XI at 1 mg/mL was used. In addition, the digested cells were resuspended in L-15 medium containing 10% FBS. The cell suspension was then filtered using a round-bottom polystyrene test tube with a cell strainer snap cap (cat. 352235; BD Falcon), and L cells were sorted using a FACSAria Fusion flow cytometer (BD Biosciences) equipped with 5 solid-state lasers (355 nm, 15 mW; 405 nm, 85 mW; 488 nm, 50 mW; 561 nm, 50 mW; and 640 nm, 100 mW) along with a forward scatter (FSC) detector, a side scatter (SSC) detector, and an 18-channel fluorophore detector. In this experiment, the 488-nm laser and the 530/30 filter were used to excite and detect the fluorescence of FITC (GFP), while the 561-nm laser and the 610/20 filter were employed for PE-Texas Red (tdTomato). Data were collected using the FACSDiva software (v8.0.2, BD Biosciences), followed by exporting in the FCS 3.1 format. The numbers of intestinal cells of each type were quantified using FlowJo software (v10.4.0, BD Biosciences). Gating strategies were defined based on the intensity of GFP fluorescence in comparison to the non-L (GFP-negative, tdTomato-positive) intestinal epithelial cells. Specifically, intact cells from each sample were selected using FSC-A and SSC-A (left panel of Figure S1J, on a linear scale), followed by FSC-H and FSC-A to exclude doublets (middle, on a linear scale). The fluorescence intensities of FITC (GFP) and PE-Texas Red (tdTomato) were measured and plotted (right, on a logarithmic scale). The plot displaying PE-Texas Red-negative and FITC-positive populations identifies the L cells. Typically, the purity of the sorted L cells exceeds 98%, which was confirmed by an Axio Observer live microscope (Zeiss) using a small aliquot. Some 10^5^ isolated L cells were collected for each measurement and incubated in DMEM medium supplemented with 10% FBS, 2 mM L-glutamine, 100 IU of penicillin, and 100 μg/mL of streptomycin. The cells were maintained at 37 °C for 2 h in a humidified incubator with 5% CO_2_ before proceeding with further experiments. It is important to note that due to the low abundance of L cell populations in the total intestinal epithelial cells, the sorting process for each sample often exceeded 0.5 h. Consequently, at least two sorters were used during the experiment.

#### Determination of GLP-1 and insulin secretion

Levels of GLP-1 were measured as described previously,^48,155,164^ with minor modifications. Briefly, mice were fasted for 16 h (from 17:00 to 9:00 of the next day) and then subjected to refeeding, glucose gavage, or glucose injection. Mice were sacrificed at different time points, and 300 μL of blood was collected from each mouse into an ice-cold K_2_EDTA spray-coated tube containing 6 μL of diprotin A (5 mg/mL). Plasma was then prepared by centrifuging the blood at 3,000 g for 10 min at 4 °C. From the plasma (the supernatant), 50 μL was used to determine the levels of GLP-1 using the GLP-1 EIA Kit, following the manufacturer’s instructions. For measuring GLP-1 secretion in L cells, cells were allowed to grow to approximately 70% confluence in a 12-well dish. Before measurement, cells were washed twice with PBS and incubated in KRBH buffer (GLP-1 Krebs-Ringer bicarbonate-HEPES buffer; containing 4.5 mM KCl, 138 mM NaCl, 4.2 mM NaHCO_3_, 1.2 mM NaH_2_PO_4_, 2.6 mM CaCl_2_, 1.2 mM MgCl_2_, and 10 mM HEPES, pH 7.4, adjusted with NaOH, along with 0.1% (w/v) BSA) supplemented with 0.01 mg/mL diprotin A and the desired concentrations of glucose for 1 h. The buffer was then collected and centrifuged at 3,000 g for 10 min at 4 °C, and 500 μL of the supernatant was taken for measuring GLP-1 levels using the GLP-1 EIA Kit.

Levels of insulin were measured as described previously.^158,165^ Briefly, blood was collected from mice as for the GLP-1 measurement, except that 1.5-mL Eppendorf tubes were used. The blood was left at room temperature for 30 min, followed by centrifugation at 3,000 g for 10 min at 4 °C. Some 25 μL of serum (the supernatant) was used to determine insulin levels using the Mouse Ultrasensitive Insulin ELISA kit according to the manufacturer’s instructions. For measuring insulin secretion in pancreatic β cells, cells were allowed to grow to approximately 70% confluence in a 12-well dish, followed by washing with PBS twice and incubation in KRBH buffer (insulin KRBH buffer; 3.6 mM KCl, 135 mM NaCl, 2 mM NaHCO_3_, 0.5 mM NaH_2_PO_4_, 1.5 mM CaCl_2_, 0.5 mM MgSO_4_, 10 mM HEPES, pH 7.4, adjusted with NaOH, and 0.2% BSA), containing varying concentrations of glucose for 1 h. Buffer was then collected and centrifuged at 3,000 g for 10 min at 4 °C, and 500 μL of supernatant was used to measure insulin levels using the Mouse Insulin ELISA Kit. A 5-parametre logistic fitted standard curve for calculating the concentrations of insulin was generated from the website of Arigo Biolaboratories Co. (https://www.arigobio.cn/ELISA-calculator).

#### Measurement of glucose in blood and intestinal lumen

Levels of mouse blood glucose were measured as described previously.^166^ Briefly, mice were individually housed for a week before the experiment. After fasting for 16 h, mice were refed with food, or given glucose through gavage or injection. Blood glucose levels were measured at different time points through tail vein bleeding, drawing approximately 20 µL of blood from each mouse using the OneTouch UltraVue automatic glucometer (LifeScan).

Levels of glucose in the lumen of different intestinal segments were determined as described previously,^167^ with minor modifications. Briefly, mice were sacrificed by cervical dislocation, and the intestinal tissue was quickly excised and placed in ice-cold PBS. The tissue was then divided into segments: duodenum, jejunum, and ileum. Each segment was gently flushed with 50 µL of sterile PBS using a syringe and collected in an ice-cold Eppendorf tube. The volume of each sample was adjusted to 100 µL with ice-cold PBS. Samples were centrifuged at 10,000 g for 15 min at 4 °C to remove debris. For analysis, 2 µL of the supernatant, diluted 500-fold (for the duodenum and jejunum) or 20-fold (for the other segments), was used to determine glucose concentration using the Glucose Assay Kit.

#### Cell lines

In this study, no cell line used is on the list of known misidentified cell lines maintained by the International Cell Line Authentication Committee (https://iclac.org/databases/cross-contaminations/). The following cell lines were utilized: HEK293T cells (cat. CRL-3216) and STC-1 cells (cat. CRL-3254), both purchased from ATCC; *LRRC8A*-KO HEK293T cells (cat. YKO-HT22252), purchased from Ubigene; GLUTag cells (cat. SCC652), purchased from Merck; NCI-H716 cells (cat. CC0517) and β-TC-6 cells (cat. CC9034), purchased from CellCook; and INS-1 cells (cat. iCell-r036), purchased from iCellBioscience. Cells were maintained in DMEM, except NCI-H716 cells that were cultured in RPMI-1640 medium, and INS-1 cells in RPMI-1640 medium supplemented with 0.2% (w/v) HEPES and 0.05 mM β-mercaptoethanol. All culture media were supplemented with 10% FBS, 100 IU of penicillin, 100 mg/mL of streptomycin, and NaHCO_3_ (3.5 g/L for DMEM and 2 g/L for RPMI-1640, adjusted to a pH of 7.2). Cells were kept at 37 °C in a humidified incubator with 5% CO_2_. All cell lines were verified to be free of mycoplasma contamination and were authenticated by STR sequencing (performed by Immocell Biotechnology Corporation in Xiamen, China). For ectopic expression, PEI was used to transfect HEK293T cells at a final concentration of 10 μM, while Lipofectamine 2000 was used at a concentration of 7 µL for 3 µg of DNA to transfect β-TC-6 cells. Total DNA to be transfected for each plate was adjusted to the same amount by using relevant empty vector. Transfected cells were harvested at 24 h after transfection.

Lentiviruses, including those for knockdown or stable expression (at close-to-endogenous levels), were packaged in HEK293T cells by transfection using Lipofectamine 2000, as described previously.^163^ At 30 h post-transfection, the medium (DMEM supplemented with 10% FBS, MEM non-essential amino acids, and sodium pyruvate; approximately 2 mL in volume) was collected and centrifuged at 5,000 g for 3 min at room temperature, yielding the supernatant containing lentivirus. To infect HEK293T cells, the supernatant was mixed with 10 μg/mL (final concentration) polybrene, and added to adhered HEK293T cells that were at 40% confluence, followed by centrifugation at 3,000 g for 30 min at room temperature (spinfection). Cells were incubated for another 12 h before further treatments. For infection of STC-1, GLUTag, and β-TC-6 cells, 60% confluence was adopted. After 12 h of infection, the cells were passaged and cultured for another 12 h before an additional round of transfection prior to subsequent treatments. Cautions were taken as follows: as STC-1 cells after lentivirus infection can become fragile in the next passage, they should not be passaged with fewer than half of the original parental cells. In addition, after the cells have attached (2-3 h post-seeding), the culture medium should be refreshed to remove any unattached dead cells. Subculturing or passaging only used cells of less than 80% confluence.

The sequences of siRNA used to knockdown mouse *GK* is: 5’-GAGATGCTCTTCGACTACATCTCTG-3’; to knockdown mouse *GPI* is: 5’-GGGTTCTGGACAAGATGAAGT-3’; to knockdown mouse *G6PDx* is: 5’-GCAGGCCAACCGTCTATTCTA-3’; to knockdown mouse *G6PD2* is: 5’-GTTCTTCACGCGTAACTCCTATGTAGT-3’; to knockdown mouse *PGM1* is: 5’-GGGATTGGTCGCCTGGTTATT-3’; and to knockdown mouse *PGM2* is 5’-GCTCATTGCTGGAGGTAATAA-3’.

#### Plasmids

The BacMam expression vectors carrying cDNA for SUR1 and Kir6.2, used for transfecting K_ATP_ into HEK293T cells for the inside-out patch clamping experiment, were provided by Dr. Lei Chen (Peking University; ref. ^141,142^). Full-length cDNAs used in this study were obtained either by PCR using cDNA from MEFs, or by purchasing from Origene or Sino Biological. Mutations of GK were performed by PCR-based site-directed mutagenesis using PrimeSTAR HS polymerase. Expression plasmids for various epitope-tagged proteins were constructed in the pcDNA3.3 vector (cat. #K830001; Thermo) for ectopic expression in mammalian cells via transfection, in the pBOBI vector for lentivirus packaging in mammalian cells, or in the pGEX-4T-1 vector (cat. 27-1542-01; Cytiva) for bacterial expression. The lentivirus-based vector pLV-H1-EF1a-puro (cat. #SORT-B19; Biosettia) was used for the expression of shRNA in STC-1 and β-TC-6 cells. PCR products were verified by sequencing (Invitrogen, China). *Escherichia coli* strain DH5α (cat. #PTA-1977) was purchased from ATCC, and Stbl3 (cat. #C737303) from Thermo. All plasmids were amplified in *E*. *coli* strain DH5α, except those for mutagenesis in Stbl3. All plasmids used in this study were purified by the CsCl density gradient ultracentrifugation method. All expression plasmids constructed in this study have been deposited to Addgene (https://www.addgene.org/Sheng-cai_Lin/).

#### Immunoprecipitation and immunoblotting

The interaction between endogenous GK and K_ATP_ was determined as described previously,^168^ with minor modifications. Briefly, STC-1 cells of a 10-cm dish (grown to 80% confluence) were collected for immunoprecipitation (IP) of GK. Cells were lysed with 750 μL/dish of ice-cold Triton lysis buffer (20 mM Tris-HCl, pH 7.5, 150 mM NaCl, 1 mM EDTA, 1 mM EGTA, 1% (v/v) Triton X-100, 2.5 mM sodium pyrophosphate, 1 mM β-glycerophosphate, with protease inhibitor cocktail), followed by sonication using a VCX 130PB sonicator (Sonics; equipped with a 5/64’’ (2 mm) stepped microtip, cat. 630-0423) for 3-5 sec at 30% maximum power on ice, and centrifugation at 4 °C for 15 min. Cell lysates were incubated with anti-GK antibody overnight. Overnight protein aggregates were pre-cleared by centrifugation at 20,000 g for 10 min; protein A/G beads (1:250, pre-balanced with lysis buffer) were then added into the lysate/antibody mixture for another 3 h at 4 °C. The beads were centrifuged and washed 3 times each with 100 times the volume of ice-cold lysis buffer (by centrifuging at 2,000 g) at 4 °C and then mixed with an equal volume of 2× SDS sample buffer before immunoblotting.

The interaction between ectopically expressed GK and K_ATP_ was determined as described previously,^169,170^ with minor modifications. Briefly, a 6 cm-dish of HEK293T cells was transfected with various combination of expression plasmids that include different epitope tags. At 24 h after transfection, cells were collected and lysed in 500 µL of ice-cold Triton lysis buffer, followed by sonication and centrifugation at 4 °C for 15 min. For IP of the FLAG epitope tag, anti-FLAG M2 Affinity Gel (1:200, pre-balanced in lysis buffer) was added into the supernatant and mixed for 1□h at 4□°C. The beads were washed with 200 times the volume of ice-cold Triton lysis buffer wash buffer 3 times at 4 °C, and then eluted with 30 μL of FLAG Peptide (400 μg/mL final concentration) for another 30 min at 4□°C. Some 30 μL of eluent was then collected, mixed with 7.5 μL of 5× SDS buffer, and then subjected to immunoblotting analysis. Note that samples containing K_ATP_, particularly those containing the transmembrane segments, were not boiled to avoid the formation of insoluble aggregates that would fail to run into the SDS-PAGE.

To analyze the levels of GK, MCT1 and G6Pase in L cells, cells grown to 70-80% confluence in a 6-well dish were lysed with 250 μL of ice-cold Triton lysis buffer. The lysates were then centrifuged at 20,000 g for 10 min at 4 °C, and an equal volume of 2× SDS sample buffer was added into the supernatant. Except MCT1, samples were boiled for 10 min and then directly subjected to immunoblotting.

To analyze the levels of GK in intestinal tissues, a protocol designed to enhance the protein recovery rate from tissue samples was used,^171^ with minor modifications. Briefly, freshly excised intestinal tissues were quickly infused with 10 mL of ice-cold PBS (supplemented with a 2× protease inhibitor cocktail) using a syringe to fill their lumens. Approximately 100 mg of tissue was then transferred to a 1.5-mL Eppendorf tube, followed by homogenization on ice using an electrical disperser (T 10 basic ULTRA-TURRAX 558 from IKA, equipped with an S10N-5G Dispersing Element) in 1 mL of 2× Sample Buffer (100 mM Tris-HCl, pH 6.8, 4% (m/v) SDS, 20% (v/v) glycerol, and 4% (v/v; added freshly before use) 2-mercaptoethanol). The homogenate was briefly vortexed and then sonicated on ice. It is important to avoid generating bubbles during sonication, given the presence of SDS in the buffer, as they can disrupt the stability of proteins in the lysate. Some 50 µL of the homogenate was then transferred to a new 1.5-mL Eppendorf tube, diluted with 450 µL of distilled water, and mixed with 50 µL of ice-cold, 10× TCA solution (72% trichloroacetic acid in water). This mixture was vortexed for 30 sec and then centrifuged at 20,000 g for 10 min at 4 °C. The resulting pellet was suspended in 1 mL of ice-cold acetone and centrifuged at 20,000 g for 10 min at 4 °C. This step was repeated two more times. After the final centrifugation, the sediment was roughly dried on ice for 5 min and then suspended in 200 µL of 2× Sample Buffer for 2 h at room temperature, with sonication every 30 min for 2-3 sec each time until the pellet was completely dissolved. Samples were then mixed with 2 µL of bromophenol blue solution (2.5% in water), boiled for 10 min, and then subjected to immunoblotting. In this study, all samples were subjected to IB on the same day of preparation, avoiding freeze-thaw cycles.

For IB, the SDS-polyacrylamide gels were prepared in-house, as described previously.^133^ The thickness of the gels used in this study was 1.0 mm. Samples of less than 10 μL were loaded into wells, and the electrophoresis was run at 100 V (by PowerPac HC High-Current Power Supply, Bio-Rad) in a Mini-PROTEAN Tetra Electrophoresis Cell (Bio-Rad). In this study, all samples were resolved on 8% resolving gels, except those for GK^Δ92-CT^ were run on 15% gels (prepared as those for 8%, except that a final concentration of 15% Acryl/Bis was added to the resolving gel solution). In addition, Kir6.2 and MCT1 were resolved on 10% gels. The resolved proteins were then transferred to the PVDF membrane (0.45 μm, cat. IPVH00010, Merck) as described previously.^133^ The PVDF membrane was then blocked by 5% (w/v) non-fat milk dissolved in TBST for 2 h on an orbital shaker at 60 rpm at room temperature, followed by rinsing with TBST (40 mM Tris, 275 μM NaCl, 0.2% (v/v) Tween-20, pH 7.6) twice, 5 min each. The PVDF membrane was then incubated with the desired primary antibody overnight at 4 °C on an orbital shaker at 60 rpm, followed by rinsing with TBST three times, 5 min each at room temperature, and then the secondary antibodies for 3 h at room temperature with gentle shaking. The secondary antibody was then removed, and the PVDF membrane was further washed with TBST 3 times, 5 min each, at room temperature. PVDF membrane was incubated in an ECL mixture (by mixing equal volumes of ECL solution and Peroxide solution for 5 min), then life with Medical X-Ray Film (FUJIFILM). The films were then developed with X-OMAT MX Developer (Carestream), and X-OMAT MX Fixer and Replenisher solutions (Carestream) on a Medical X-Ray Processor (Carestream) using Developer (Model 002, Carestream). Films were then scanned using a Perfection V850 Pro scanner (Epson) with an Epson Scan software (v.3.9.3.4), cropped using Photoshop 2023 software (Adobe), and formatted into panels using Illustrator 2022 (Adobe). Levels of total proteins and phosphorylated proteins were analyzed on separate gels, and representative immunoblots are shown. Uncropped immunoblots are provided.

#### Confocal microscopy

The plasma membrane localization of Lck-PercevalHR indicator was determined as described previously,^159,172^ with minor modifications. Briefly, cells grown to 80% confluence on coverslips in 6-well dishes were fixed for 20 min with 4% (v/v) formaldehyde in PBS at room temperature. The coverslips were rinsed twice with PBS and permeabilized with 0.1% (v/v) Triton X-100 in PBS for 5 min at 4 °C. After rinsing twice with PBS, the sections were blocked with PBS containing 5% BSA for 30 min at room temperature. Then the coverslips were incubated with anti-N-cadherin antibody (1:100, diluted in PBS) overnight at 4 °C. The cells were then rinsed 3 times with 1 mL of PBS, and then incubated with Alexa Fluor 594-conjugated, goat anti-rabbit IgG antibody (1:200 in PBS) for 8 h at room temperature in the dark. Cells were washed another 4 times with 1 mL of PBS and then mounted on slides using ProLong Diamond antifade mountant. Confocal microscopy images were taken using a STELLARIS 8 FALCON (Leica) systems equipped with HyD SMD detectors and an HC PL APO CS2 63x/1.40 OIL objective (Leica). All parameters were kept unchanged between imaging. Images were taken and analyzed by LAS X Software (v. 3.0.2.16120, Leica), and formatted using Photoshop 2023 software (Adobe).

The local intracellular fluctuation of the ADP:ATP ratios near the plasma membrane was assessed using the Lck-PercevalHR indicator, as previously described.^95,160^ In brief, cells expressing Lck-PercevalHR were cultured in 35-mm glass-bottom dishes (cat. D35-20-10-N, In Vitro Scientific) to 60-80% confluence. Cells were starved of glucose for 2 h, followed by equilibration at 37 °C and 5% CO_2_ in a humidified incubation chamber for an additional 30 min (cat. STXG-WELSX-SET; TOKAI HIT). Cells were imaged using a spinning disk confocal microscope (Olympus IXplore SpinSR, equipped with a Yokogawa CSU-W1 spinning disk with a 50-μm diameter) and a 60×/1.4 N.A. objective, utilizing the cellSens Dimension software (ver. 4.1.1). In the meantime, ProLong Live antifade reagent was added to the medium before imaging. Lck-PercevalHR was excited with 490-nm and 400-nm lasers, respectively. The emitted light was separated from the excitation light by a 490-nm short-pass dichroic mirror and collected through a 520/10 nm band-pass filter. The ratio of the intensities of the emission light at 490 nm to that at 400 nm reflects the ATP:ADP ratio.

#### FRET-FLIM assay

FRET-FLIM experiments were carried out as described previously,^147,173^ with minor modifications. Briefly, STC-1 cells stably expressing GFP (“donor only”, as a control) or different combinations of GFP-GK, Kir6.2-mCherry and mCherry-SUR1 were cultured in 35-mm glass-bottom dishes to 60-80% confluence. Cells were starved for glucose or treated with 25 mM glucose, followed by determining the fluorescence lifetime of GFP in different cells cultured in a humidified chamber with 5% CO_2_ at 37 °C using a STELLARIS 8 FALCON (Leica) system equipped with HyD X and HyD SMD detectors and an HC PL APO CS2 63x/1.40 OIL objective (Leica). Cells were excited with a 460-nm laser via the systems’ tunable White Light Laser (WLL), and photon arrival times were recorded with a HyD X detector covering the GFP emission spectrum (460-510 nm). All parameters were kept unchanged between imaging. Images were taken, and the donor lifetime, FRET efficiency, and donor-accepter distance were analyzed using LAS X Software (Leica). In all experiments, the position of the focal plane was actively stabilized using the Leica Auto Focus Control (AFC) to prevent any focal drift or focus artefacts.

#### Measurement of adenylates

ATP, ADP, and AMP from intestinal and pancreatic β cell lines, as specified in the legends, were analyzed by CE-MS as described previously,^157,174^ with minor modifications. Briefly, each measurement required cells collected from a 10-cm dish (60-70% confluence). Cells were rinsed with 20 mL of 5% (m/v) mannitol solution (dissolved in water) and instantly frozen in liquid nitrogen. Cells were then lysed with 1 mL of methanol containing IS1 (50 µM L-methionine sulfone, 50 µM D-campher-10-sulfonic acid, dissolved in water; 1:500 (v/v) added to the methanol and used for standardization of the metabolite intensity and to adjust the migration time), and were scraped from the dish. The lysate was then mixed with 1 mL of chloroform and 400 μL of water by 20 sec of vortexing. After centrifugation at 15,000 g for 15 min at 4 °C, 450 μL of aqueous phase was collected and then filtered through a 5-kDa cutoff filter (cat. OD003C34, PALL) by centrifuging at 12,000 g for 3 h at 4 °C. In parallel, quality control samples were prepared by combining 10 μL of the aqueous phase from each sample and then filtered alongside the experimental samples.

The filtered aqueous phase was then freeze-dried in a vacuum concentrator at 4 °C, and then dissolved in 100 μL of water containing IS2 (50 µM 3-aminopyrrolidine dihydrochloride, 50 µM N,N-diethyl-2-phenylacetamide, 50 µM trimesic acid, 50 µM 2-naphtol-3,6-disulfonic acid disodium salt, dissolved in methanol; used to adjust the migration time). A total of 20 μL of re-dissolved solution was then loaded into an injection vial (cat. 9301-0978, Agilent; equipped with a snap cap (cat. 5042-6491, Agilent)). Before CE-MS analysis, the fused-silica capillary (cat. TSP050375, i.d. 50 µm × 80 cm; Polymicro Technologies) was installed in a CE/MS cassette (cat. G1603A, Agilent) on the CE system (Agilent Technologies 7100). The capillary was then pre-conditioned with Conditioning Buffer (25 mM ammonium acetate, 75 mM diammonium hydrogen phosphate, pH 8.5) for 30 min, followed by balanced with Running Buffer (50 mM ammonium acetate, pH 8.5; freshly prepared) for another 1 h. CE-MS analysis was run in anion mode, during which the capillary was washed by Conditioning Buffer, followed by injection of the samples at a pressure of 50 mbar for 25 sec, and then separation with a constant voltage at −30 kV for another 40 min. Sheath Liquid (0.1 μM hexakis(1H, 1H, 3H-tetrafluoropropoxy)phosphazine, 10 μM ammonium trifluoroacetate, dissolved in methanol/water (50% v/v); freshly prepared) was flowed at 1 mL/min through a 1:100 flow splitter (Agilent Technologies 1260 Infinity II; actual flow rate to the MS: 10 μL/min) throughout each run. The parameters of MS (Agilent Technologies 6545) were set as: a) ion source: Dual AJS ESI; b) polarity: negative; c) nozzle voltage: 2,000 V; d) fragmentor voltage: 110 V; e) skimmer voltage: 50 V; f) OCT RFV: 500 V; g) drying gas (N_2_) flow rate: 7 L/min; h) drying gas (N_2_) temperature: 300 °C; i) nebulizer gas pressure: 8 psig; j) sheath gas temperature: 125 °C; k) sheath gas (N_2_) flow rate: 4 L/min; l) capillary voltage (applied onto the sprayer): 3,500 V; m) reference (lock) masses: m/z 1,033.988109 for hexakis(1H, 1H, 3H-tetrafluoropropoxy)phosphazine, and m/z 112.985587 for trifluoroacetic acid; n) scanning range: 50-1,100 m/z; and o) scanning rate: 1.5 spectra/s. Data were collected using MassHunter LC/MS acquisition 10.1.48 (Agilent Technologies), and were processed using Qualitative Analysis B.06.00 (Agilent Technologies). Levels of AMP, ADP, and ATP were measured using full scan mode with m/z 346.0558, 426.0221, and 505.9885, respectively. Note that a portion of ADP and ATP could lose one phosphate group during in-source-fragmentation thus leaving same m/z ratios as AMP and ADP, and this was corrected according to their different retention times in capillary. Therefore, the total amount of ADP is the sum of the latter peak of the m/z 346.0558 spectrogram and the former peak of the m/z 426.0221 spectrogram, and ditto for ATP. For quantification of AMP, ADP and ATP, [U-^13^C, ^15^N]AMP, [U-^13^C, ^15^N]ADP and [U-^13^C, ^15^N]ATP dissolved in individual lysates were used to generate corresponding standard curves by plotting the ratios of detected labelled AMP, ADP or ATP (areas) to the products of IS1 and IS3, against the added concentrations of labelled AMP, ADP or ATP. Levels of [U-^13^C, ^15^N]AMP, [U-^13^C, ^15^N]ADP and [U-^13^C, ^15^N]ATP were measured using full scan mode with m/z 351.0408, 431.0071, and 510.9735, respectively. The amounts of AMP, ADP and ATP were estimated according to standard curves, and were then divided by protein wet weight. The protein wet weight of each sample was determined by Bradford assay after dissolving the naturally dried protein sediment with 0.2 M KOH at room temperature. Cell density of 1.1□g/mL (ref. ^175^) were used.

ATP, ADP, and AMP from intestinal and pancreatic β cell lines as specified in the legends, as well as from intestinal tissues, were analyzed by standard HPLC-MS as previously described,^176,177^ with minor modifications. In brief, each measurement required cells collected from a 10-cm dish (60-70% confluence). Cells were rinsed with 20 mL of PBS and instantly frozen in liquid nitrogen. To analyze adenylates in intestinal tissues, mice were anaesthetized, and the duodenum or jejunum tissues were quickly excised. The tissues were then rinsed with 20 mL of PBS and freeze-clamped. Following this, cells and tissues were lysed with 1 mL of methanol. The lysate was then mixed with 1 mL of chloroform and 400□μL water (containing 4□μg/mL [U-^13^C]-glutamine used for standardization of the metabolite intensity and to adjust the migration time), by 20 sec of vortexing. After centrifugation at 15,000 g for 15 min at 4 °C, 450 μL of aqueous phase was collected, lyophilized in a vacuum concentrator at 4□°C and then dissolved in 80□μL of 50% (v/v) acetonitrile. Some 50□μL of supernatant was loaded into an injection vial (5182-0714, Agilent Technologies; with an insert (HM-1270, Zhejiang Hamag Technology)) equipped with a snap cap (HM-2076, Zhejiang Hamag Technology). Measurements of adenylate levels were based on ref. ^178^ using a QTRAP MS (QTRAP 5500, SCIEX) interfaced with a UPLC system (ExionLC AD, SCIEX). A total of 2□μL of each sample was loaded onto a HILIC column (ZIC-pHILIC, 5 μm, 2.1□×□100□mm, PN: 1.50462.0001, Millipore). The mobile phase consisted of 15□mM containing 3□mL/L ammonium hydroxide (>28%, v/v) in LC-MS grade water (mobile phase□A) and LC-MS grade 90% (v/v) acetonitrile in LC-MS grade water (mobile phase□B) run at a flow rate of 0.2□mL/min. Metabolites were separated with the following HPLC gradient elution program: 95% B held for 2□min, then to 45% B in 13□min, held for 3□min, and then back to 95% B for 4□min. The MS was run on a Turbo V ion source in negative mode with a spray voltage of −4,500□V, source temperature of 550□°C, gas no.1 at 50□psi, gas no.2 at 55□psi, and curtain gas at 40□psi. Metabolites were measured using the multiple reaction monitoring mode, and declustering potentials and collision energies were optimized using analytical standards. The following transitions were used for monitoring each compound: 505.9/158.9 and 505.9/408.0 for ATP; 425.9/133.9, 425.9/158.8 and 425.9/328.0 for ADP; 345.9/79.9, 345.9/96.9 and 345.9/133.9 for AMP; and 149.9/114 for [U-^13^C]-glutamine. Data were collected using Analyst software (v.1.7.1, SCIEX), and the relative amounts of metabolites were analyzed using MultiQuant software (v.3.0.3, SCIEX). Similar to CE-MS analysis, a portion of ADP and ATP could lose one or two phosphate groups during in-source-fragmentation, thus leaving the same m/z ratios as AMP and ADP, which were corrected according to their different retention times in the column. For quantification of AMP, ADP and ATP, [U-^13^C, ^15^N]AMP, [U-^13^C, ^15^N]ADP and [U-^13^C, ^15^N]ATP dissolved in individual lysates were used to generate corresponding standard curves as in CE-MS analysis, except that [U-^13^C]-glutamine was used as an internal standard. The following transitions were used for monitoring each compound: 520.9/158.9 for [U-^13^C, ^15^N]ATP; 440.9/143.9 for [U-^13^C, ^15^N]ADP; and 360.9/79.9 for [U-^13^C, ^15^N]AMP. The amounts of AMP, ADP and ATP were estimated according to standard curves, and were then divided by protein wet weight as described in the CE-MS analysis.

The dissolved samples were left on ice or at 4 °C for no more than 4 h before injection into the MS.

#### Measurement of glycolytic metabolites

Levels of FBP from L cells were analyzed using CE-MS as described previously.^173^ The sample preparation, CE conditions, and parameters for mass spectrometry are outlined in the CE-MS analysis section of “Measurement of Adenylates”, except that an m/z value of 338.9888 was used to quantify FBP.

The levels of 2-PG and 3-PGA in L cells were analyzed using an improved HPLC-MS method that offers enhanced sensitivity compared to standard HPLC-MS. The HPLC conditions and separation program were revised based on the chemical properties of 2-PG and 3-PGA. Briefly, samples were collected and processed following the standard HPLC-MS analysis described in the section “Measurement of Adenylates.” The lyophilised samples were re-dissolved in 60□μL of 50% (v/v) acetonitrile. Some 50□μL of supernatant was then loaded into an injection vial (5182-0714, Agilent Technologies; with an insert (HM-1270, Zhejiang Hamag Technology)) equipped with a snap cap (HM-2076, Zhejiang Hamag Technology). Measurements of 2-PG and 3-PGA levels were performed using a triple quadrupole MS (Xevo TQ Absolute; Waters) interfaced with a UPLC system (ACQUITY Premier; Waters). A total of 2□μL of each sample was loaded onto a HILIC column (Atlantis Premier BEH Z-HILIC VanGuard FIT Column, 1.7□μm, 2.1□×□100□mm, PN: 186009982; Waters). The mobile phase consisted of 15□mM ammonium bicarbonate supplemented with 2.5 µM medronic acid in LC-MS grade water (mobile phase□A) and 15□mM ammonium bicarbonate containing 2.5 µM medronic acid in LC-MS grade 90% (v/v; in LC-MS grade water) acetonitrile (mobile phase□B), run at a flow rate of 0.35□mL/min. Metabolites were separated using the following HPLC gradient elution program: 90% B held for 1 min, then reduced to 65% B over 8 min, held for an additional 2 min, and returned to 90% B in 30 sec, holding for 210 sec. The MS was run on a Zspray ion source in negative mode with the following parameters: a) capillary voltage: −2,000□V; b) desolvation temperature: 500□°C; c) desolvation gas flow: 500 l/h; and d) cone gas flow: 150 l/h. Metabolites were measured using the multiple reactions monitoring mode, with parameters of cone voltage and collision energy optimized using analytical standards. The following transitions were used for monitoring each compound: 185.00/97.00 for 2-PG, 185.01/97.00 for 3-PGA, and 150/114 for [U-^13^C]-glutamine. Data were collected, and the relative amounts of metabolites were analyzed using MassLynx (version 4.2; Waters).

Levels of other glycolytic intermediates in L cells were measured using standard HPLC-MS as described in the section “Measurement of Adenylates”, except that 3 μL of sample was loaded onto the HILIC column. The following transitions were used for monitoring each compound: 179.1/89.1 for glucose; 259.0/96.9 for G6P; 259.0/97.0 for F6P; 169.0/97.0 for DHAP, 169.0/79.0 for G3P, 265.0/78.9 for 1,3-BPG, 166.9/78.9 for PEP, and 87.0/43.0 for pyruvate. Data were collected using Analyst software (v.1.7.1, SCIEX), and the relative amounts of metabolites were analyzed using MultiQuant software (v.3.0.3, SCIEX).

#### Protein Expression

The expression plasmids for GK and mutants were constructed by inserting respective cDNAs into pGEX-4T-1 vectors for expressing GST-tagged recombinant proteins as described previously.^104,162^ Briefly, plasmids were transformed into the *E*. *coli* strain BL21 (DE3) (cat. EC0114; Thermo), followed by culturing in LB medium in a shaker at 200 rpm at 37 °C. Expression of the proteins in the bacteria were induced with 0.1 mM IPTG at an OD_600_ of 1.0. After incubating for another 12 h at 160 rpm at 16 °C, the cells were collected, and homogenized in a GST binding buffer (PBS supplemented with 10 mM β-mercaptoethanol and 1% Triton X-100) on ice. The homogenates were then sonicated on ice and subjected to centrifugation at 150,000 g for 30 min at 4 °C, followed by purification with Glutathione Sepharose 4 Fast Flow Gel (pre-balanced with GST binding buffer) at 4 °C. The Glutathione Sepharose gel was then washed with 100 times the volume of ice-cold PBS, and then incubated with 10 U/mL thrombin (dissolved in PBS, freshly prepared) at 4 °C for 8 h to cleave the GST tag from GK, releasing the GK from the Glutathione Sepharose gel. Proteins were concentrated to approximately 3 mg/mL by ultrafiltration (Millipore, UFC905096) at 4 °C, then subjected to gel filtration (Cytiva, Superdex 200) balanced with the same buffer as that used in the enzymatic assays or patch-clamping assays before proceeding with each experiment.

#### Enzymatic activity

Enzymatic activity of GK was determined using glucose as a substrate and ATP as a co-substrate, employing two distinct methods: one to determine the velocity of G6P formation to represent the catalytic activity of GK towards glucose, which was coupled with the G6PD enzyme,^179^ and the other measured the velocity of ATP consumption by coupling the PK-LDH enzymes,^161^ as previously described.

For the G6PD-coupled assay, 1.8 mL of G6PD Reaction Buffer (20 mM HEPES, pH 7.7, 125 mM KCl, 7.5 mM MgCl_2_, 5 mM ATP, 0.5 mM NADP^+^, 0.5 U/mL G6PD, and the desired concentrations of glucose) was added into a 4-mL cuvette on a Lambda 365 Spectrophotometer (PerkinElmer) equipped with magnetic stirrer, set to stir at 140 rpm and at a temperature of 37 °C. After a 10-min incubation, the cuvette was taken out and gently tapped to eliminate bubbles. The cuvette was then placed back on the spectrophotometer until the baseline OD_340_ reading became flat and stable, with the change in the value remaining less than 0.001. The reaction was initiated by addition of 6 μL of GK diluted with the Reaction Buffer to the desired concentration and pre-incubated with its activator at the chosen concentration at 37 °C for 5 min. Data were collected using UV WinLab software (v.7.1.0.68, PerkinElmer) and exported to GraphPad Prism (ver. 10.1.2) for further analysis. The concentrations of glucose consumed were calculated by the extinction coefficient for NADPH at 340□nm, which is 6,220□cm^−1^ M^−1^. The initial velocity of glucose consumption (measured within 30 sec to 1 min after the reaction was initiated), represented by the initial rates of NADPH formation, was then calculated at each glucose concentration, plotted, and the *K*_m_ and V_max_ values were determined based on the curves fitted to the Michaelis-Menten equation.

The PK-LDH-coupled assay was performed in a similar manner to the G6PD-coupled assay, with the use of PK-LDH Reaction Buffer (100 mM Tris-HCl, pH 7.9, 100 mM KCl, 20 mM MgCl_2_, 0.2 mM NADH, 1.0 mM PEP, 20 U/mL LDH, 15 U/mL PK, 5 mM ATP, and the desired concentrations of glucose). The concentrations of ATP consumed were calculated using the extinction coefficient for NADH at 340 nm, which is 6,220□cm^−1^ M^−1^. Following this, the initial rates of NADH formation were calculated to reflect the velocity of ATP consumption.

#### Determination of binding affinity of GK towards glucose analogs

The binding affinity of GK towards glucose analogs was determined using fluorescence-labelled differential scanning fluorimetry (FL-DSF) methods, as described previously.^180^ Briefly, GK was conjugated with FITC-NHS dye via the amine coupling method prior to FL-DSF analysis. Before conjugation, 1 mL of Labelling Buffer (8 mM Na_2_HPO_4_, 137 mM NaCl, 1.47 mM KH_2_PO_4_ and 2.68 mM KCl, pH 8.0) containing 20 µM GK was mixed with 1 µL of FITC-NHS dye (100 µM, prepared from a 100 mM stock solution dissolved in anhydrous DMSO). The mixture was gently rotated for 30 min at 4 °C in the dark. The reaction was then terminated by adding a Tris-HCl solution (1 M stock solution, pH 8.5) to achieve a final concentration of 10 mM. Unconjugated dye was removed using a desalting column (Zeba Spin, 7 kDa MWCO; cat. 89891; Thermo) pre-equilibrated with the Labelling Buffer. Some 10 µL of FITC-GK conjugate was then incubated with 10 µL of MDGP, 2-DG, or glucose as a control, at varying concentrations at room temperature for 5 min in a PCR tube.

FL-DSF analysis was conducted using a CFX96 Touch Real-Time PCR Detection System (Bio-Rad) with the “SYBR” program. A temperature gradient was established, ranging from 25 °C to 95 °C, with increments of 0.2 °C. The system held each temperature for 5 sec before taking measurements. Data, represented as fluorescent intensity (F), were collected at 2-s intervals. Following this, melting curves were generated using the CFX Manager software (version 3.1.1517.0823; Bio-Rad). The negative first derivative (-dF/dT) at each temperature (T) of the melting curve was calculated using the “Melt Curve Derivative Results” function of the software. The melting temperature (T_m_), defined as the maximum value of -dF/dT, was obtained using the “Draw Data” function of OriginPro 2024 software. The Δ*T*_m_ was calculated by subtracting the T_m_ of concentration of each glucose analog from the T_m_ of the glucose analog-free group. These Δ*T*_m_ values were then plotted against the concentrations of the glucose analogs using GraphPad Prism 10.1.2 software. The binding constants (*K*_D_) were determined using a one-site specific binding model.

#### Electrophysiology

Whole-cell patch clamping was performed as described previously, with minor modifications.^181,182^ Briefly, *LRRC8A*-KO HEK293T cells were used for minimizing the current caused by chloride ion efflux through volume-regulated anion channels (VRACs), which could otherwise interfere with the K_ATP_-mediated current resulting from K^+^ influx when a negative voltage was applied during whole-cell patch clamping.^183,184^ Cells were transfected with Kir6.2 and SUR1 as described in the “Cell lines” section, except that Lipofectamine 3000 was used as the transfection reagent, and cells at a density of 10^6^ cells/mL were infected with a lentivirus carrying wildtype GK or its G80A and L309P mutants as described in the “Cell lines” section. For the other experiments, STC-1 and GLUTag cells were used, and were directly infected with lentiviruses carrying wildtype GK or its mutants. These cells were then incubated in glucose-free DMEM for 2 h, followed by trypsinization. The cells were then cultured in a 12-well dish containing a poly-L-lysine-coated coverslip in glucose-free DMEM for another 1 h in a humidified chamber with 5% CO_2_ at 37 °C, for cells to attach to the coverslips. These coverslips were then transferred to a 500-μL recording chamber that was placed on the objective lens of an inverted Axio Examiner microscope (Zeiss) and filled with bath (extracellular) solution. This solution contained 10 mM HEPES, 120 mM KCl, 5 mM NaCl, 1 mM MgCl_2_, 2 mM CaCl_2_ and various concentrations of glucose, supplemented with NMDG and MSA to bring the pH to ∼ 7.2 and final osmolarity to ∼ 300 mOsm, as determined by a VAPRO vapor pressure osmometer (cat. 5600; ELITechGroup). Each bath solution was loaded into the syringe of a perfusion system driven by a multichannel peristaltic pump set to a flow rate of 2 mL/min. Borosilicate glass electrodes (micropipettes) with a tip resistance of 2-4 MΩ were pulled using a Next Generation Micropipette Puller (P-1000; Sutter Instrument) and filled with pipette (cytosolic face) buffer containing 150 mM KCl, 2 mM MgCl_2_, 20 mM HEPES supplemented with 2 mM Mg-ATP, 200 μM ADP (freshly prepared), and the desired concentrations of glucose (supplemented with NMDG and MSA to bring the pH to ∼ 7.2 and final osmolarity to ∼ 300 mOsm). An agar bridge filled with 3 M KCl served as the reference electrode. The plasma membrane of a cell was suctioned using the micropipette to form a gigaohm seal, followed by membrane rupture via gentle suction to achieve whole-cell access. The whole-cell K^+^ current was recorded by applying a ramp protocol from −120 mV to +120 mV (for HEK293T cells) or from −140 mV to +60 mV (for STC-1 and GLUTag cells) over a duration of 400 msec, repeated every 2 sec, using an Axon Axopatch 200B Microelectrode Amplifier (Molecular Devices) equipped with a Digidata 1440A Low-noise Data Acquisition System (Molecular Devices). The glucose concentration was altered either in the bath solution using the perfusion system or in the pipette solution by using pipette filled with different glucose concentrations. Data collection was performed using Clampex 10.7 software (Molecular Devices). All recordings were conducted at room temperature, and data were analyzed using Clampfit 10.7 (Molecular Devices) and Prism 10 (GraphPad).

Inside-out patch clamping was performed as described previously,^141,185^ with minor modifications. Briefly, HEK293T cells were transfected with wildtype Kir6.2 or its Kir6.2-205-228A mutant, as described above. After 48 h of transfection, the cells were aliquoted and cultured on the coverslips as outlined in the whole-cell patch clamping section. Borosilicate glass electrodes with a tip resistance of 6-9 MΩ were pulled. The pipette (extracellular) solution contained 140 mM KCl, 2.6 mM CaCl_2_, 1 mM NaCl, 1.2 mM MgCl_2_, and 10 mM HEPES (pH 7.4, by KOH), while the bath (cytosolic) solution contained 130 mM NaCl, 5 mM KCl, 2 mM CaCl_2_, 1 mM MgCl_2_, 10 mM HEPES (pH 7.4, by NaOH) supplemented with desired concentrations of glucose and NMDG to bring the final osmolarity of 300 mOsm. Before the experiment, the bath solution was freshly supplemented with 2 mM Mg-ATP, 200 μM ADP, and GK protein (purified as described in the “Protein Expression” section, which involved gel filtration using a column balanced and run with the bath solution). The plasma membrane of a cell was suctioned using the micropipette to form a gigaohm seal, after which the pipette was rapidly withdrawn and immersed back into the bath solution to establish the inside-out configuration. Excised patches were continuously perfused with bath solution at a flow rate of 1 mL/min. Single-channel recording was performed using a step protocol at +60 mV. Inward current (i.e., cation flux from the extracellular to the cytosolic side of the membrane) is indicated as downward deflections in all recorded traces. Signals were acquired at 20 kHz and low-pass filtered at 5 kHz. Data were analyzed with Clampfit 10.7 (Molecular Devices) software, with a filter set at 0.5 kHz.

#### Protein and peptide MS

Protein levels of K_ATP_ channel subunits expressed in STC-1 cells were quantified using liquid chromatography coupled with data-independent acquisition (DIA) mass spectrometry, as previously described,^151,186^ with modifications. In brief, peptides from STC-1 cells were generated using a Single-Pot Solid-Phase-enhanced Sample Preparation (SP3) protocol,^187^ with minor modifications. First, cells were lysed using 1% (v/v) SDS dissolved in distilled water, followed by sonication and centrifugation as described in the “Immunoprecipitation and Immunoblotting” section. The lysates were then adjusted to a protein concentration of 100 µg/mL using the BCA assay. Some 100 µL of lysates were then incubated with 10 mM dithiothreitol (final concentration, dissolved in water) on a thermomixer at 56 °C, at 1,000 rpm for 30 min, followed by an incubation with 20 mM iodoacetamide (final concentration, dissolved in water) at 25 °C, at 1,000 rpm for another 30 min in the dark. The reaction was terminated by adding 20 mM dithiothreitol at 25 °C, at 1,000 rpm for an additional 10 min. Subsequently, 100 µg of SP3 beads was added to the lysate, followed by addition of an equal volume of ethanol and incubation at 25 °C, at 1,000 rpm for 30 min. The beads were washed three times with 80% (v/v) ethanol and then resuspended in 50 µL of 50 mM ammonium bicarbonate (dissolved in water), followed by overnight incubation with 0.5 µg of trypsin. The resulting peptide supernatants were mixed with 1% (v/v) trifluoroacetic acid and desalted using a C18 StageTips column (cat. MC18TB; Nanomics). Before desalting, the column was washed twice with 50% (v/v) acetonitrile and equilibrated three times with 0.1% (v/v) trifluoroacetic acid. Peptides were loaded onto the column, washed three times with 0.1% (v/v) trifluoroacetic acid, and eluted twice with 80% (v/v) acetonitrile (40 µL each). The peptide eluents were then combined and dried in a vacuum centrifugal concentrator, followed by resuspension in 20 µL of 0.1% (v/v) formic acid. Some 200 ng of peptides were then separated on a Vanquish Neo upgraded ultra-high-performance liquid chromatography (UHPLC) system (Thermo), which was tandemly equipped with a PepMap Neo Trap Cartridge (5 mm × 300 µm, 5 µm, cat. 174500; Thermo) and an EASY-Spray HPLC Column (150 µm × 150 mm, 2 µm, cat. ES906; Thermo). The mobile phase consisted of 0.1% (v/v) formic acid in LC-MS grade water (designated as mobile phase A) and 0.1% (v/v) formic acid in 80% (v/v) acetonitrile (in LC-MS grade water, mobile phase B). The peptides were eluted over a period of 13 min using gradients that ranged from 4% to 99% mobile phase B. The gradient began with 4% B for 0.3 min at a flow rate of 1.8 µL/min, followed by a linear increase to 8% B over the next 0.7 min, and then a further linear increase to 22.5% B over the subsequent 6.7 min. The gradient then continued with a linear increase to 35% B over the following 3.7 min, during which the flow rate changed to 2.5 µL/min. Next, the gradient increased linearly to 55% B within 0.4 min, followed by another linear increase to 99% B in 0.5 min, and finally held steady at 99% B for an additional 0.7 min.

Data acquisition was performed using an Orbitrap Astral mass spectrometer (Thermo) equipped with an EASY-Spray Nanosource in DIA mode. During measurements, the parameters for MS1 were set to a resolution of 240,000, covering a m/z range of 380 to 980, with a normalized automatic gain control (AGC) value of 500% and a maximum injection time (maxIT) of 5 msec. For fragmentation, peptides were processed by the astral detector using normalized higher energy collisional dissociation (HCD) set at 25%, with a normalized AGC target value of 500% and a maxIT of 3 msec. The fixed window number was set to 300, with a 2 m/z isolation window to cover the mass range from 150 to 2000 m/z. The DIA raw files were analyzed using DIA-NN software (v1.8.1),^144^ and peptide and protein identifications were conducted using the UniProt database (*Mus musculus*) and visualized using IBS software (v 2.0).^145^ A confidence level of 99% or higher for peptide-spectrum matches (PSM) was established as the filter criterion, and a false discovery rate (FDR) validation was applied to exclude peptides and proteins with an FDR greater than 1%. For a complete set of parameters used for the identification and quantification of target proteins, please refer to the source data files deposited in the iProX partner repository accompanying the paper.

### QUANTIFICATION AND STATISTICAL ANALYSIS

Statistical analyses were performed using Prism 9 (GraphPad Software). Each group of data was subjected to Kolmogorov-Smirnov test, Anderson-Darling test, D’Agostino-Pearson omnibus test or Shapiro-Wilk test for normal distribution when applicable. An unpaired two-sided Student’s *t*-test was used to determine the significance between two groups of normally distributed data. An unpaired two-sided Mann-Whitney test was used to determine the significance between data without a normal distribution. For comparisons between multiple groups with a fixed factor, an ordinary one-way ANOVA was used, followed by Tukey, Sidak or Dunnett as specified in the legends. The assumptions of homogeneity of error variances were tested using F-test (*P* > 0.05). For comparison between multiple groups with two fixed factors, an ordinary two-way ANOVA was used, followed by Tukey’s or Sidak’s multiple comparisons test as specified in the legends. Geisser-Greenhouse’s correction was used where applicable. The adjusted means and SEM were recorded when the analysis met the above standards. Differences were considered significant when *P* < 0.05, or *P* > 0.05 with large differences of observed effects (as suggested in refs. ^188,189^).

---

**Figure S1.**
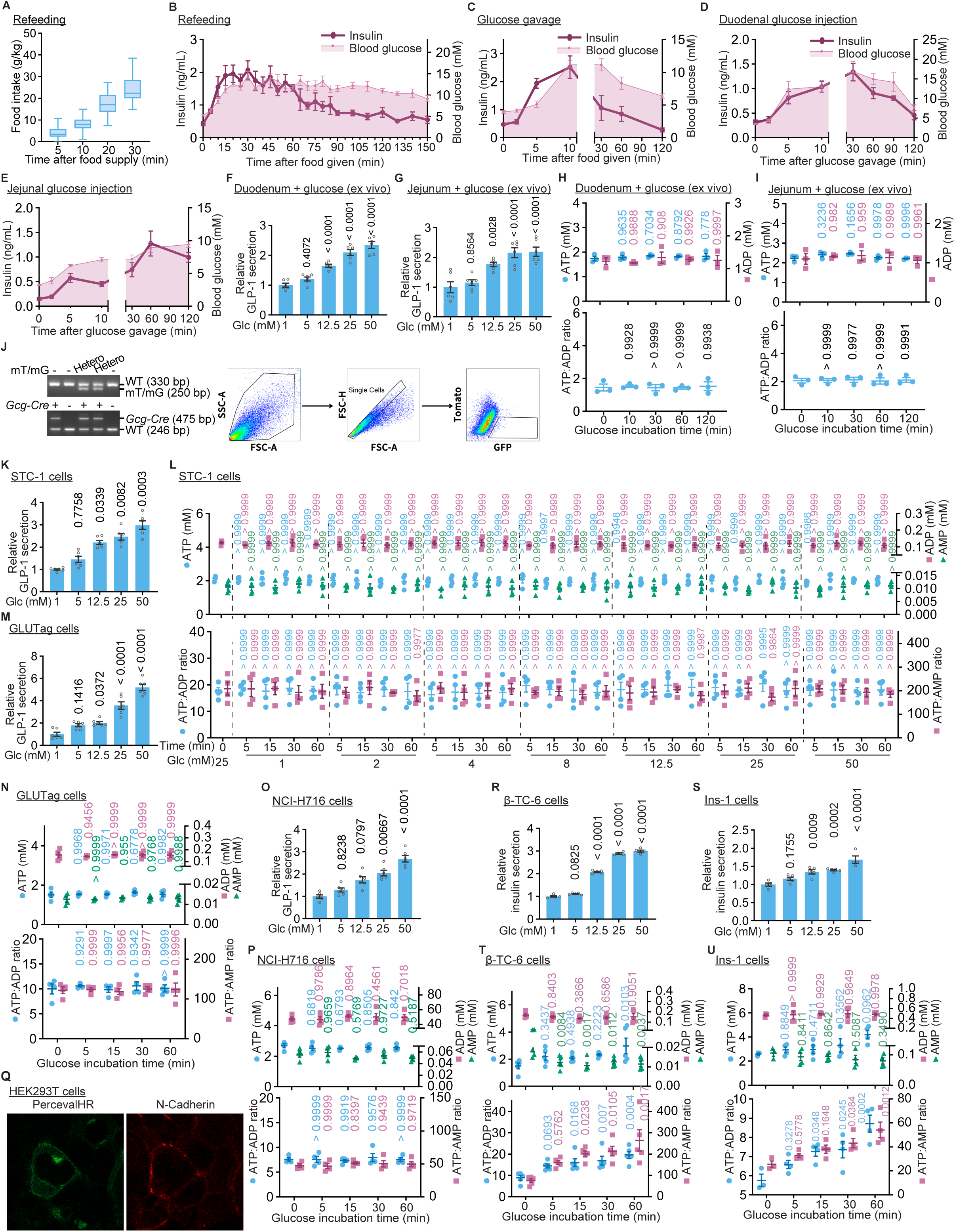
High glucose does not elevate ATP in L cells during GLP-1 secretion, but elevates ATP in pancreatic β cells during insulin secretion, related to Figure 1. (A) Determination of food amounts that can be consumed within 10 min by mice fasted for 16 h. Mice were fasted for 16 h and then re-fed with normal chow diet. At different time points after the supply of chow, the food consumption was measured and shown as box-and-whisker plots (the lower and upper bounds of the box represent the first and the third quartile scores, the center line represents the median, and the lower and upper limits denote minimum and maximum scores, respectively); n = 60 mice for each time point. For each experiment performed as in Figure 1A, only the mice whose 10-min food consumption fell between the first and third quartile scores were included for calculating glucose, GLP-1, and insulin levels. (B-E) Insulin levels were measured in mice treated as in Figures 1A-D. Mice were fasted for 16 h, and re-fed with normal chow diet for 10 min (B; each mouse consumed 10 g/kg of body weight food, as validated in Figure S1A), or received a gavage of 1.5 g/kg glucose (C) or an injection of 75 g/L glucose directly into the duodenum (D), or jejunum (E), at a volume of 10 µL/g after enterostomy. The serum insulin and the blood glucose levels were then measured. Results are shown as mean ± SEM; n = 8 (B), 7 (C), or 6 (D and E) mice for each time point. (F-I) Glucose stimulates GLP-1 secretion without changing ATP levels in different intestinal segments. Mice were fasted for 16 h; the freshly excised duodenum (F and H) and jejunum (G and I) tissues were incubated in the GLP-1 KRBH buffer (GLP-1 Krebs-Ringer bicarbonate-HEPES) containing glucose at indicated concentration (F and G), or in the GLP-1 KRBH buffer containing 25 mM glucose for indicated time durations (H and I). The concentrations of GLP-1 in the KRBH medium (F and G), and the adenylate levels in the tissues (H and I; by HPLC-MS) were measured. Results are shown as mean ± SEM; n = 3 (H and I) or 6 (others) mice or tissues for each treatment, and *p* value by one-way ANOVA, followed by Tukey (F) or Dunnett (others), compared to the 1 mM Glc group (F and G) or the untreated group (others). (J) Isolation of L cells from mice engineered to express GFP specifically in L cells. Mice with L cells specifically expressing GFP (genotyping data are shown in the first panel) were fasted for 16 h, and the intestines were excised and digested, followed by sorting of GFP-expressing L cells using flow cytometry. During sorting, intact cells from each sample were selected based on FSC-A and SSC-A (second panel, using a linear scale), followed by FSC-H and FSC-A to exclude doublets (third panel, using a linear scale). To identify GFP-expressing L cells, gating was performed using a combination of FITC (GFP) and PE-Texas Red (tdTomato) (last panel, using a logarithmic scale). Cells positive for FITC and negative for PE-Texas Red represented the L cells. (K-P) High glucose does not elevate ATP in L cell lines during stimulation of GLP-1 secretion. STC-1 cells (K and L), GLUTag cells (M and N), and NCI-H716 cells (O and P) were incubated in GLP-1 KRBH buffer containing varying concentrations of glucose for 1 h (K, M, and O), or in GLP-1 KRBH buffer containing 25 mM glucose for indicated time durations (L, N and P), followed by determination of the concentrations of GLP-1 in the medium (K, M, and O) and the levels of adenylates (L, N and P; by CE-MS). Results are shown as mean ± SEM; n = 4 (L, N and P) or 6 (others) biological replicates for each glucose concentration or time point, and *p* value by two-way ANOVA, followed by Dunnett (adenylate ratios of L) or Tukey (others of L), or one-way ANOVA, followed by Tukey (K, M, and O) or Dunnett (N and P). (Q) Validation for the expression and proper localization of the plasma membrane-localized PercevalHR indicator in cells. HEK293T cells were infected with lentivirus carrying a PercevalHR indicator fused to the Lck protein at its N-terminus (depicted in the right panel). The infected cells were fixed with formaldehyde and incubated with anti-N-cadherin antibody, followed by incubation with Alexa Fluor 594-conjugated, goat anti-rabbit IgG antibody. Imaging was performed using confocal microscopy with laser beams set to 490 nm and 594 nm, respectively (representative images are shown in the left panel). (R-U) High glucose elevates ATP levels in pancreatic β cells during the insulin secretion. β-TC-6 (R and T) and Ins-1 cells (S and U) were treated with 25 mM glucose for indicated time durations (T and U), or incubated in Insulin KRBH buffer containing varying concentrations of glucose for 1 h (R and S), followed by determination of the adenylate levels (T and U; by CE-MS), and the concentration of insulin in the buffer (R and S). Results are shown as mean ± SEM; n = 3 (1min of U), 4 (25 mM of R, 50 mM of S,5min, 15min, 30min, and 60min of U), or 5 (others) biological replicates for each treatment, and *p* value by one-way ANOVA, followed by Dunnett. Experiments in this figure were performed three times, except A, T, and S four times, N five times, and K and R six times.

**Figure S2.**
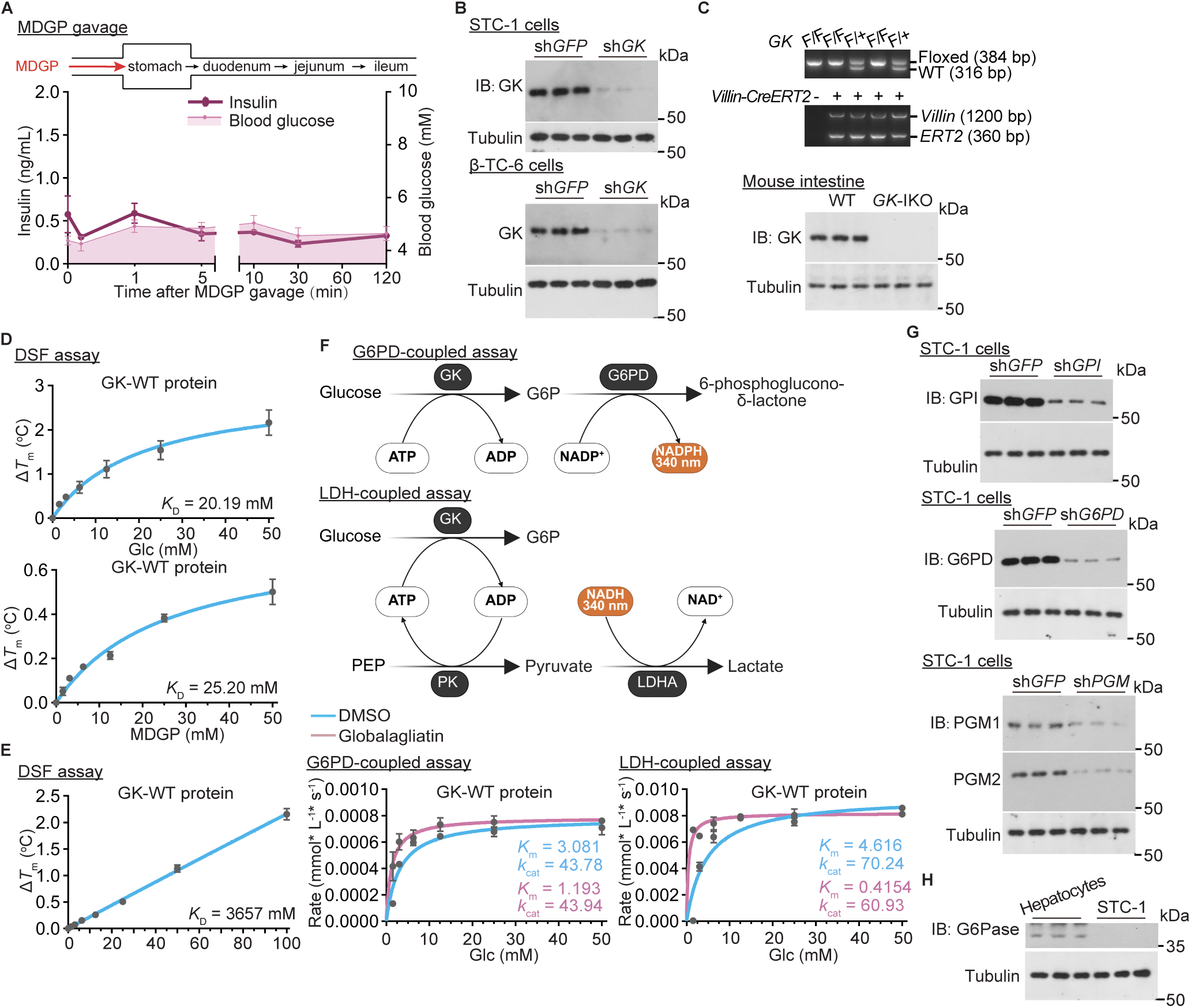
Glucokinase senses glucose to trigger GLP-1 secretion, related to Figure 2. (A) Insulin levels measured in mice gavaged with MGDP. Mice were fasted for 16 h, and then gavaged with 1.5 g/kg MGDP (as in Figure 2A). The serum insulin and the blood glucose levels were then measured. Results are shown as mean ± SEM; n = 8 mice for each time point. (B and G) Validation of knockdown efficiency of GK and downstream enzymes in L cells. STC-1 (B and G) and β-TC-6 (B) cells were infected with lentivirus carrying the shRNAs targeting *GK* (B), *GPI* (G, upper panel), *G6PD* (both *G6PD2* and *G6PDx*; G, middle panel), and *PGM* (both *PGM1* and *PGM2*; G, lower panel). Cells were lysed, and the protein levels of each enzyme were determined by immunoblotting. (C) Validation of intestinal-specific GK knockout (*GK*-IKO) in mice. *GK*-floxed mice were crossed with *Villin*-*CreERT2* mice to generate intestine-specific knockout of *GK* after tamoxifen injection. The genotyping and immunoblotting data related to the expression levels of GK in intestinal tissues are shown. (D and E) Nonmetabolizable glucose analog MDGP, but not 2-DG, binds GK. Some 20 µM GK, conjugated with FITC-NHS, was incubated with varying concentrations of MDGP (D) or 2-DG (E) in PBS. The binding affinities were assessed using differential scanning fluorimetry. The fluorescence intensities of FITC were monitored during thermal denaturation (25-95 °C, ramp rate 1 °C/min), and the ligand concentration-dependent changes in melting temperature (Δ*T*_m_) were plotted, from which *K*_D_ values were derived by fitting Δ*T*_m_ versus ligand concentration to the Hill equation (n = 3 (1.56 mM MDGP of D, and 1.56 mM and 3.125 mM 2-DG of E) or 4 (others) biological replicates). (F) The GK activator globalagliatin increases the binding affinity of GK for glucose. GK activity was measured by its ability to catalyze the conversion of glucose to G6P, coupled with the production of ADP from ATP. The yielded G6P was then converted to 6-phosphoglucono-δ-lactone with concomitant reduction of NADP^+^ to NADPH through the catalysis of G6PD. The increase of OD_340_ of NADPH was then quantified through a microplate reader, and then converted to the changes of amount of NADPH by the Beer-Lambert law to determine the activity of GK (depicted in upper panel). Alternatively, the GK activity was also assessed by measuring the amount of ADP generated, which is converted back to ATP through pyruvate kinase (PK) in conjunction with formation of pyruvate from phosphoenolpyruvate (PEP). The resultant pyruvate was then determined by quantifying the decrease of OD_340_ of NADH levels, catalyzed by lactate dehydrogenase A (LDHA) (depicted in the upper panel). In both assays, 50 nM GK was incubated with 5 mM ATP and 10 µM globalagliatin. Initial reaction rates were measured at varying glucose concentrations. Data points represent mean ± SEM from triplicate experiments. The curve was fitted to the Michaelis-Menten equation using nonlinear regression. (H) G6Pase expression is marginal in L cells. STC-1 cells were lysed, followed by determination of G6Pase expression by immunoblotting. As a control, primary hepatocytes were treated with serum-free Williams’ E medium containing 1 µM dexamethasone plus 500 µM 8-bromo-cAMP for 12 h to induce the expression of G6Pase. Experiments in this figure were performed three times.

**Figure S3.**
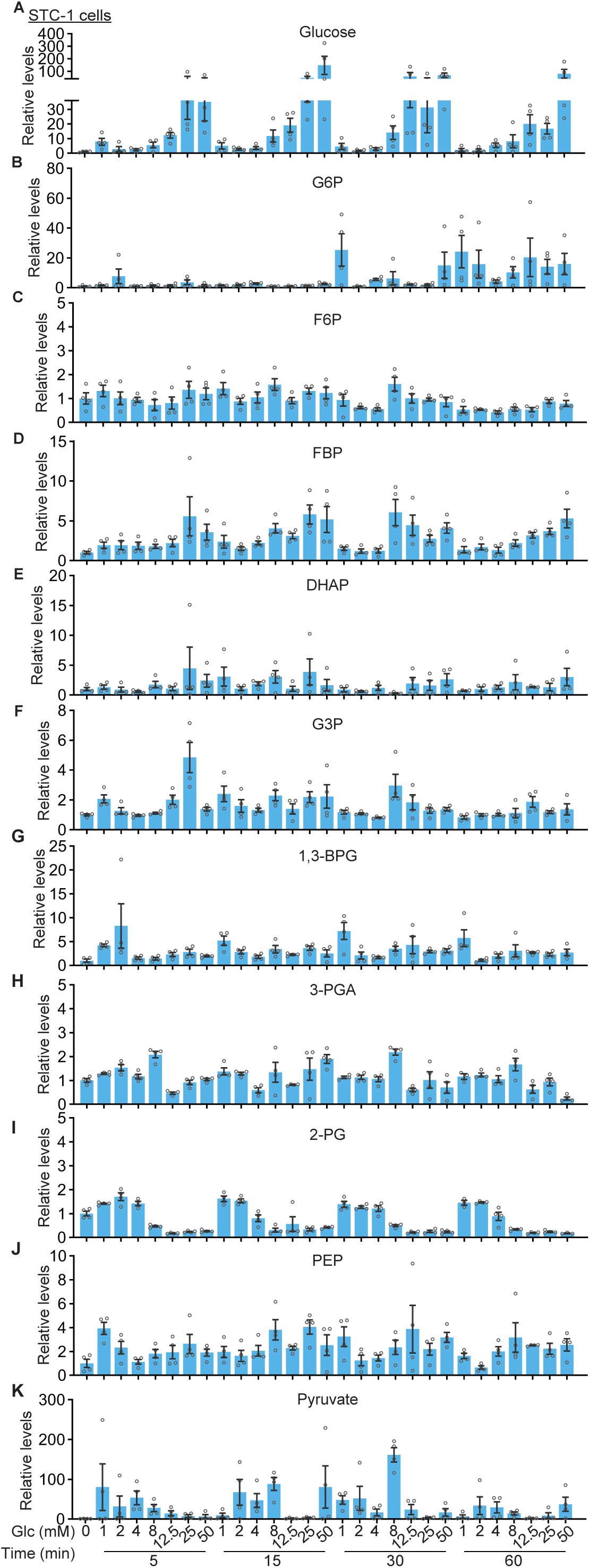
GLP-1 secretion is positively correlated with intracellular glucose concentrations but not glycolytic intermediates, related to Figure 2. (A-K) STC-1 cells were treated with varying concentrations of glucose for indicated time durations. The intracellular levels of glucose (A) and glycolytic intermediates including G6P (B), fructose-6-phosphate (F6P; C), FBP (D), glyceraldehyde-3-phosphate (G3P; E), dihydroxyacetone phosphate (DHAP; F), 1,3-bisphosphoglycerate (1,3-BPG; G), 3-phosphoglycerate (3-PGA; H), 2-phosphoglycerate (2-PG; I), PEP (J), and pyruvate (K), were then determined using CE-MS (for FBP) or HPLC-MS (for others). Results are shown as mean ± SEM; n = 3 (1 mM and 50 mM Glc at 60 min of I, and 1 mM Glc at 60 min of H) or 4 (others) biological replicates for each condition. Experiments in this figure were performed three times.

**Figure S4.**
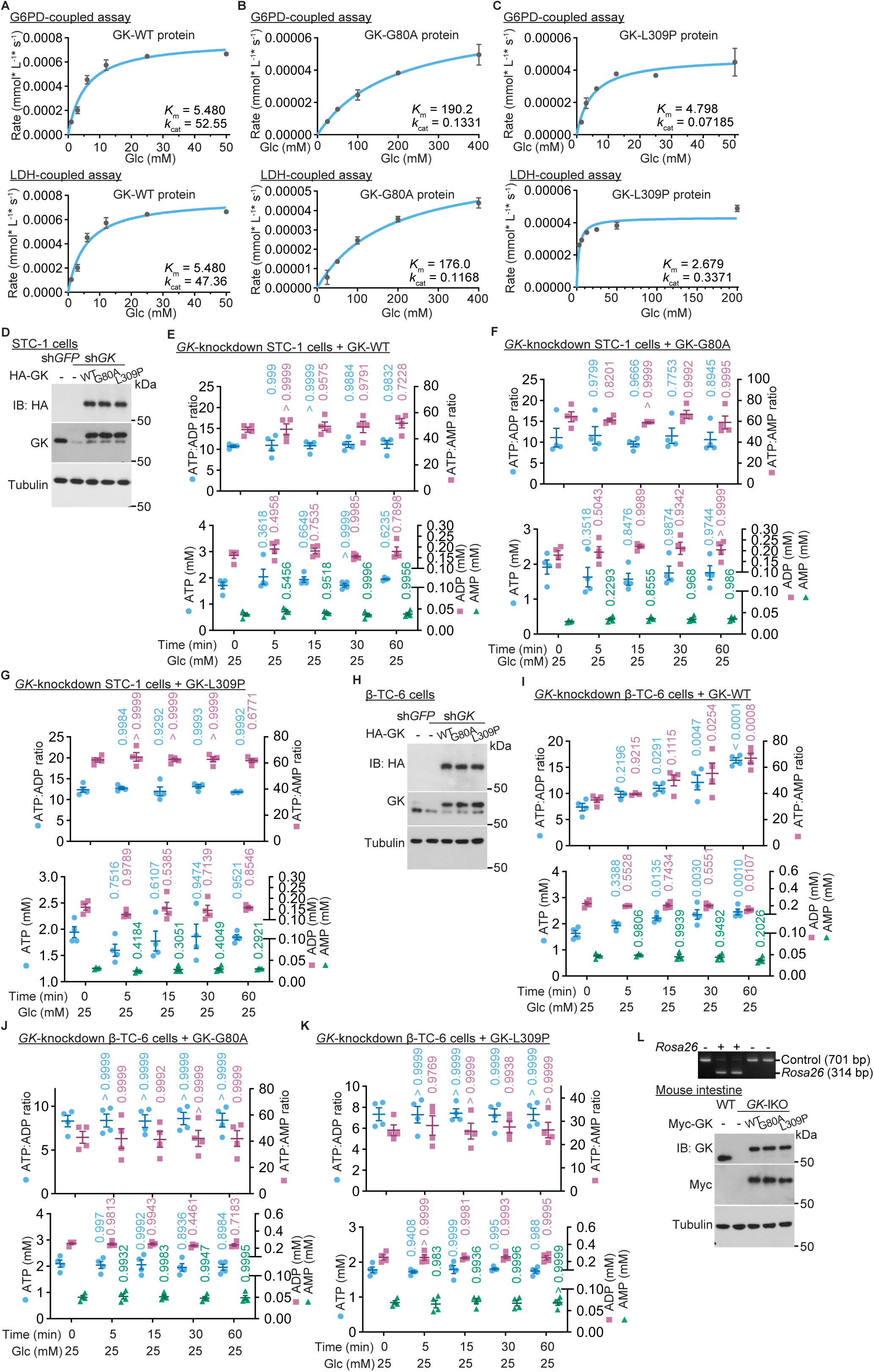
Glucose-binding affinity but not catalytic activity of GK determines its ability to GLP-1 secretion in L cells, related to Figure 3. (A-C) The G80A mutant of GK fails to bind glucose, while the L309P mutant of GK binds glucose constitutively, although both mutants lack catalytic activity towards glucose. Enzymatic activities of GK were measured as in Figure S2F, except that the GK-G80A (B) and the GK-L309P mutants (C), and wildtype GK as a control (A) were used. Data points represent mean ± SEM from triplicate experiments. The curve was fitted to the Michaelis-Menten equation using nonlinear regression. (D) Validation of reintroduction of GK into the *GK*-knockdown STC-1 cells. *GK*-knockdown STC-1 cells were infected with lentivirus carrying either GK-G80A, GK-L309P mutant, or wildtype GK, followed by determination of the expression levels of GK and its mutants by immunoblotting. (E-G) Catalysis-dead GK mutants do not alter cellular ATP levels. GK-G80A (F) and the GK-L309P mutant (G), along with wildtype GK as a control (E), were re-introduced into *GK*-knockdown STC-1 cells via lentiviral infection. Cells were then treated 25 mM glucose for indicated time durations, followed by determination of adenylate levels using HPLC-MS. Data are shown as mean ± SEM; n = 4 biological replicates for each treatment/genotype, and *p* value by Kruskal-Wallis test (ATP:AMP of F), or by one-way ANOVA, followed by Dunnett (others). (H) Validation of GK reintroduction into *GK*-knockdown β-TC-6 cells. *GK*-knockdown β-TC-6 pancreatic cells were infected with lentivirus carrying either GK-G80A, GK-L309P mutant, or wildtype GK, followed by determination of the expression levels of GK and its mutants by immunoblotting. (L) Validation of intestine-specific GK replacement with its mutants in mice. The *GK*-IKO mice (*GK*-floxed mice crossed with *Villin*-*CreERT2* mice; validated in Figure S2C) were crossed with the mice expressing either the GK-G80A mutant, the GK-L309P mutant, or wildtype GK (as a control) driven by the *Rosa26*-*LSL*(*LoxP*-*Stop*-*LoxP*) system. In this system, the *LSL* cassette ahead of introduced GK and its mutants was removed by Cre expression in the intestine, resulting in the expression of GK. The genotyping and immunoblotting data regarding the expression levels of GK in intestinal tissues are shown. (I-K) Catalysis-dead GK mutants lead to decreased ATP levels in β-TC-6 pancreatic cells exposed to high glucose. GK-G80A (J) and GK-L309P mutant (K), along with wildtype GK as a control (I), were re-introduced into *GK*-knockdown β-TC-6 cells via lentiviral infection. Cells were then treated with 25 mM glucose for indicated time points, followed by determination of adenylate levels using HPLC-MS. Data are shown as mean ± SEM; n = 3 (5min of I), or 4 (others) biological replicates for each treatment/genotype, and *p* value by Kruskal-Wallis test (ATP:ADP of J), or by one-way ANOVA, followed by Dunnett (others). Experiments in this figure were performed three times.

**Figure S5.**
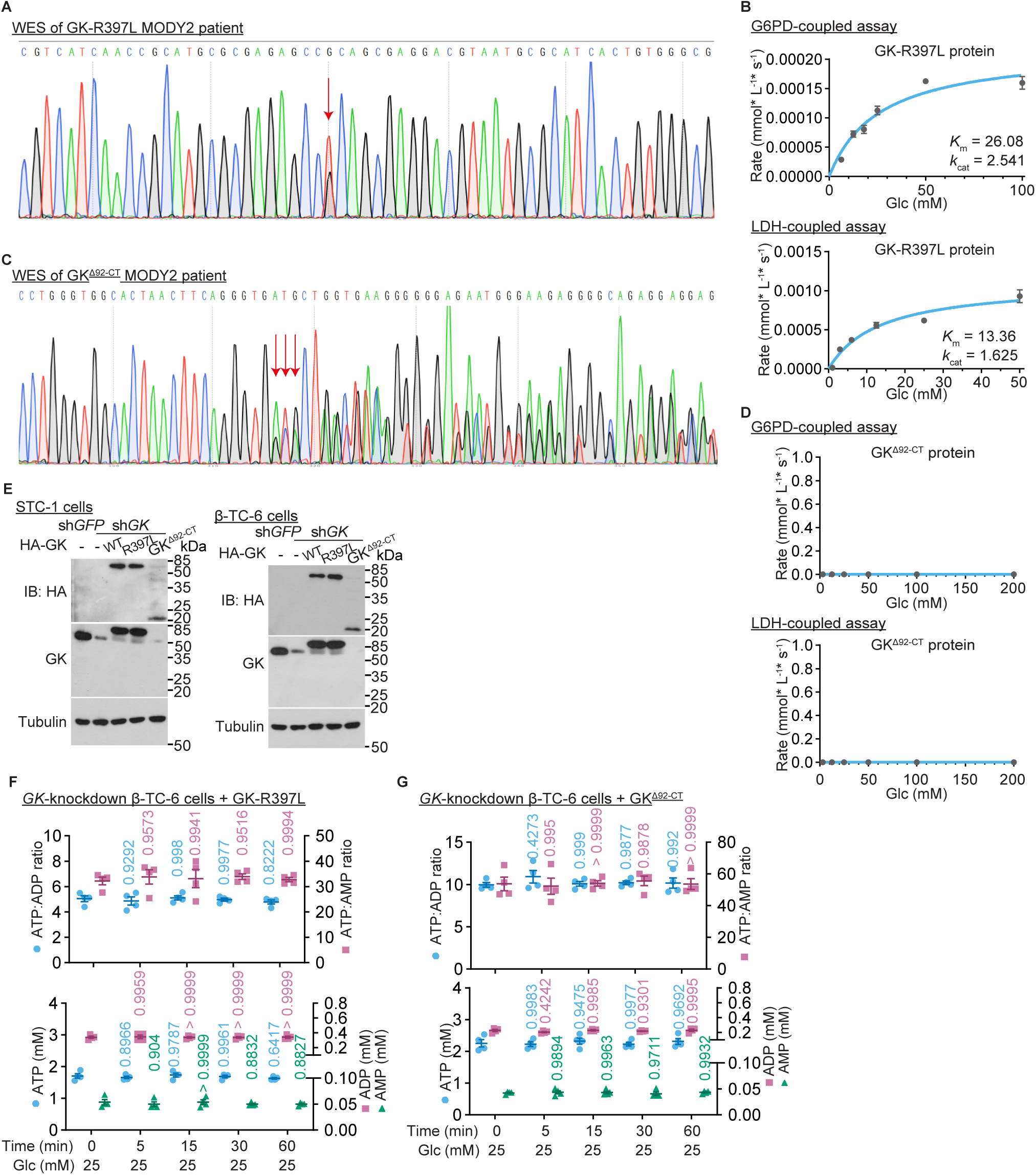
Binding affinities of GK for glucose determines its activity to trigger GLP-1 secretion in humans, related to Figure 3. (A and C) Identifications of GK mutations in MODY2 patients. Genomic DNA was extracted from the peripheral blood of MODY2 patients, followed by whole-exome sequencing (WES). The electropherograms of two patients: one carrying the heterozygous mutation, NM_000162.5:exon9:c.1190G>T (p.Arg397Leu) missense mutation (A, GK-R397L; arrow indicates dual peaks), and the other having a duplication, NM_000162.5:exon3:c.242_258dup (p.M87Afs*6) (B, GK^Δ92-CT^), are shown. (B and D) GK^Δ92-CT^ fails to bind glucose, while GK-R397L binds glucose constitutively, even though both mutants lack catalytic activity towards glucose. The enzymatic activity of GK was determined as in Figure S3E, except that the GK-R397L (B) and the GK^Δ92-CT^ (D) mutants were used. Data points represent mean ± SEM from triplicate experiments. The curve was fitted to the Michaelis-Menten equation using nonlinear regression. (E) Validation of GK^Δ92-CT^ and GK-R397L reintroduction in *GK*-knockdown STC-1 intestinal and β-TC-6 pancreatic cells. *GK*-knockdown STC-1 (left panel) or β-TC-6 cells (right panel) were infected with lentivirus carrying either GK^Δ92-CT^ or GK-R397L, followed by determination of the expression levels of GK mutants by immunoblotting. (F and G) Mutants GK^Δ92-CT^ and GK-L309P lead to decreased ATP levels in β-TC-6 pancreatic cells in high glucose. Mutant GK-R397L (F) and GK^Δ92-CT^ (G), were separately re-introduced into *GK*-knockdown β-TC-6 cells via lentiviral infection. Cells were then treated with 25 mM glucose for indicated time durations, followed by determination of ATP, ADP, and AMP levels using HPLC-MS. Data are shown as mean ± SEM; n = 4 biological replicates for each treatment/genotype, and *p* value by one-way ANOVA, followed by Dunnett. Experiments in this figure were performed three times.

**Figure S6.**
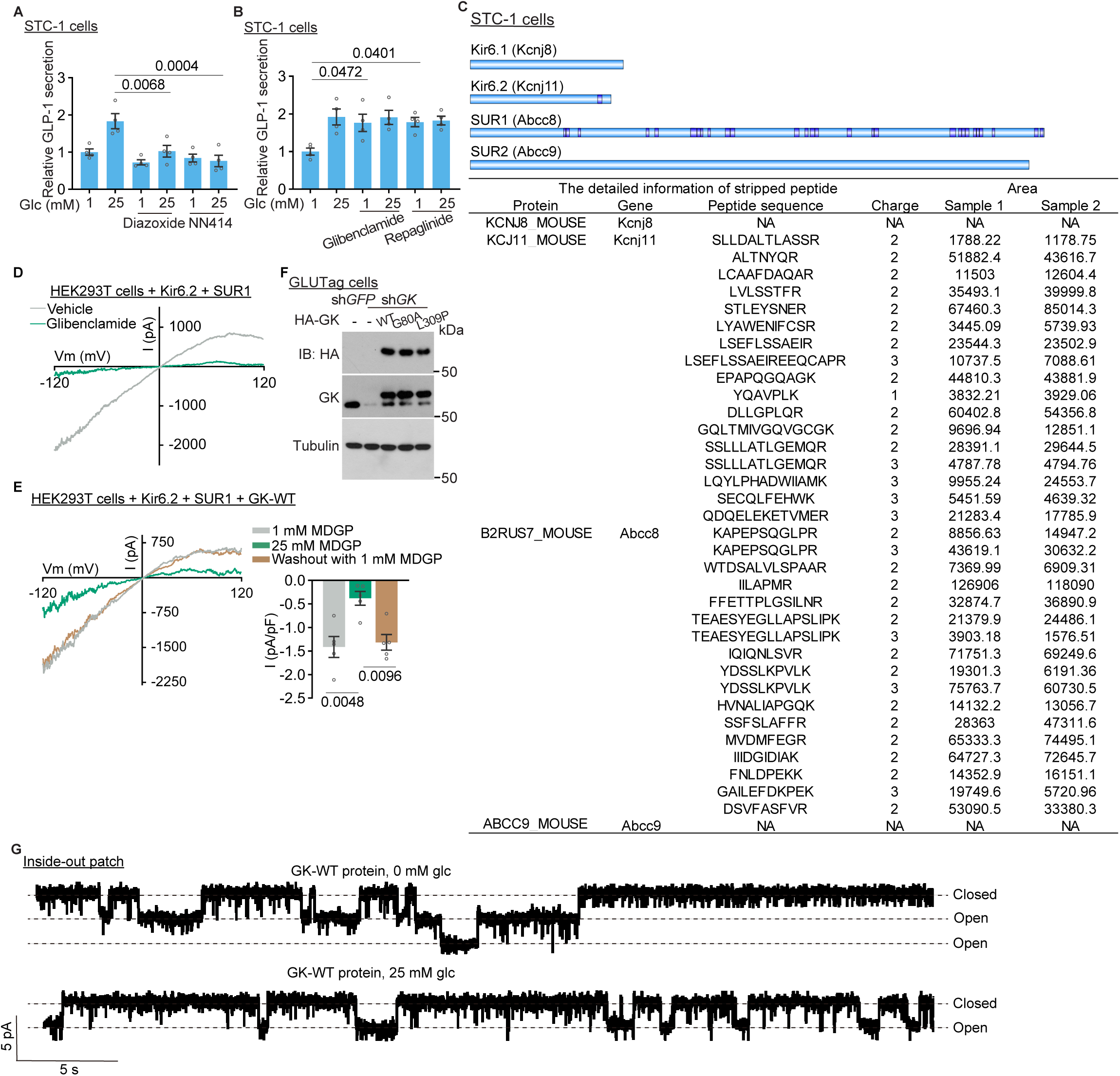
Binding of glucose is a prerequisite for GK to inhibit K_ATP_, related to Figure 4. (A and B) Inhibition of the K_ATP_ is a necessary step for GLP-1 secretion. STC-1 cells were treated with for 1 h with K_ATP_ openers diazoxide and NN414 (A) or K_ATP_ inhibitors glibenclamide and repaglinide (B), at a concentration of 10 µM. After the treatment, GLP-1 secretion was measured following an additional 1-h incubation with different concentrations of glucose. Results are shown as mean ± SEM; n = 4 biological replicates for each treatment, and *p* value by one-way ANOVA, followed by Tukey. (C) Kir6.2 and SUR1 are the major subunits of K_ATP_ channel expressed in L cells. STC-1 cells were lysed, followed by quantifying the abundance of all peptide precursors, with respective matched proteins using high-resolution mass spectrometry. The normalized quantities for precursors of K_ATP_ channel subunits—Kir6.1, Kir6.2, SUR1, and SUR2 (lower panel, from two independent samples) and their locations on the channel subunits (upper panel) are shown, indicating that Kir6.2 and SUR1 were predominantly detected in both cell types, with expression levels higher compared to Kir6.1 and SUR2, which were either barely detectable or absent. (D) Validation of the establishment of K_ATP_ current in HEK293T cells. HEK293T cells deficient of VRAC channels were transfected with Kir6.2-mCherry and Myc-SUR1. The whole-cell K^+^ currents were then measured as shown in Figure 4F, using 10 µM glibenclamide as a positive control. The representative trace of the ramp-induced currents recorded at −120 mV, and the corresponding steady-state I-V curves, are shown. (E) Nonmetabolizable glucose analog MDGP binds to GK and inhibits K_ATP_ channel. HEK293T cells deficient of VRAC channels were transfected with Kir6.2-mCherry and Myc-SUR1. Cells were starved for glucose for 2 h, and then subjected to determination of K^+^ currents by whole-cell patch clamping as in Figure 4D. During the recording, the MDGP concentration in the bath (extracellular) solution was altered from 1 mM (depicted as grey bar) to 25 mM (green bar), and then back to 1 mM (washout with 1 mM solution; yellow bar) using a perfusion system. The representative steady-state I-V curves and the statistical analysis of the ramp-induced currents recorded at −120 mV are shown in the left and right panels, respectively (mean ± SEM, n = 5 cells for each treatment, with the *p* value determined by one-way ANOVA, followed by Sidak). (F) Validation of reintroduction of GK into the *GK*-knockdown GLUTag cells. *GK*-knockdown GLUTag cells were infected with lentivirus carrying either GK-G80A, GK-L309P mutant, or wildtype GK, followed by determination of the expression levels of GK and its mutants by immunoblotting. (G) Representative current traces for wildtype GK added in the inside-out patch clamping performed in Figure 4H, with downward deflections indicating the channel openings. Experiments in this figure were performed three times, except A and B five times.

**Figure S7.**
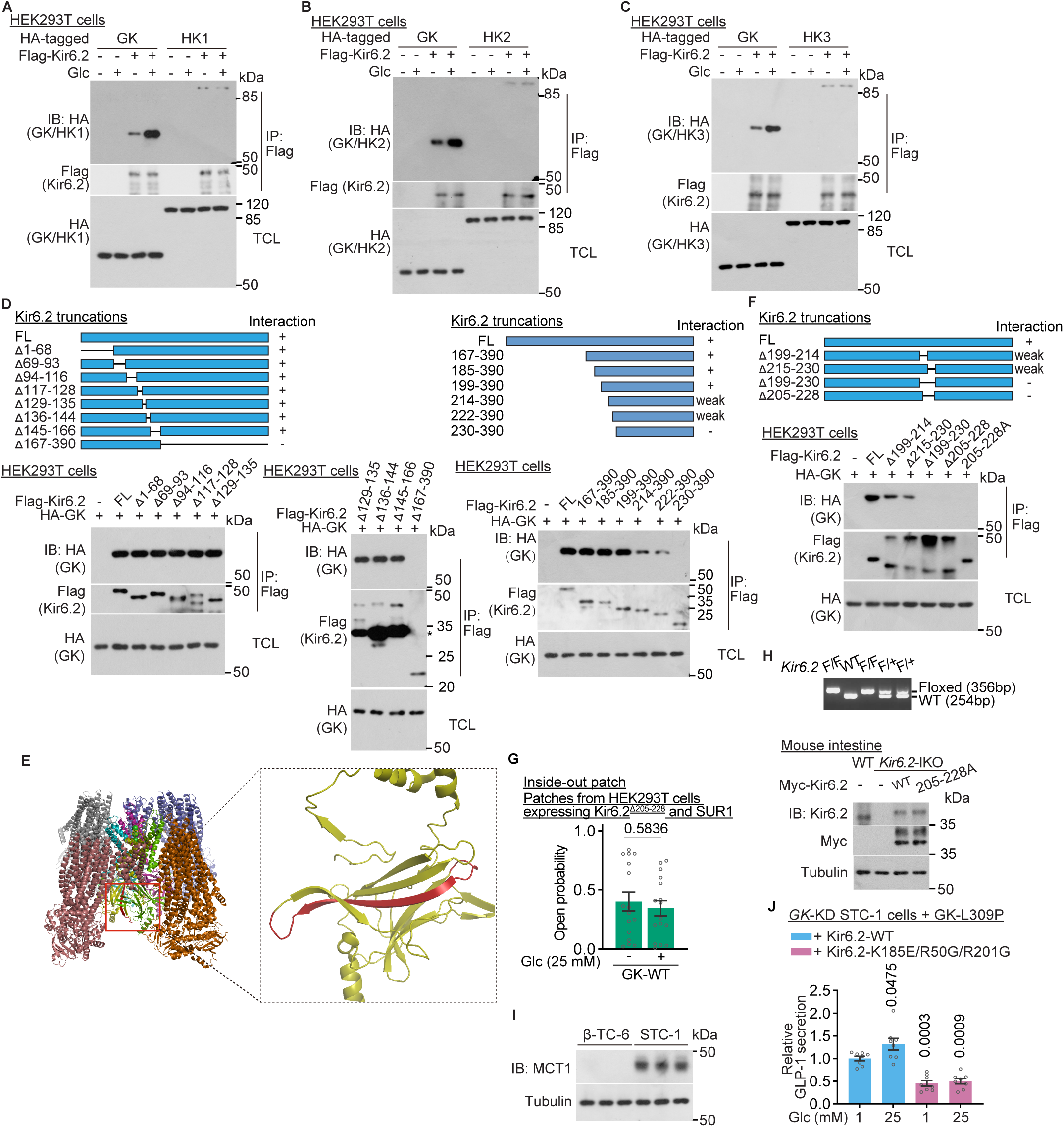
Glucose stimulates GK to interact with K_ATP_, related to Figure 5. (A-C) Hexokinases 1-3 hardly interact with K_ATP_. HEK293T cells were transfected with different combinations of HA-tagged Hexokinase 1 (HK1) (A), HK2 (B), HK3 (C), and GK as a control (A-C), alongside FLAG-tagged Kir6.2 (A-C). At 24 h post-transfection, cells were incubated in DMEM medium containing 0 mM or 25 mM glucose, followed by immunoprecipitation of the FLAG epitope tag and immunoblotting using the indicated antibodies. (D-F) GK interacts with K_ATP_ through the region of amino acid 205-228 in Kir6.2. HEK293T cells were transfected with various FLAG-tagged truncation mutants of Kir6.2 alongside HA-tagged GK, followed by immunoprecipitation of the FLAG epitope tag and immunoblotting (D). This analysis identified the region aa 199-229 of Kir6.2 as necessary for GK binding. Further refinement pinpointed the critical segment to aa 205-228, since aa 200-204 and 229-230 belong to another beta sheet adjacent to the main beta sheet structure formed by the aa 199-229 segment (E; colored in red), based on the reported structure (PDB ID: 6c3o). Due to severe degradation upon expression of the Kir6.2^Δ205–228^ truncation in HEK293T cells (F), a mutation was created where aa 205-228 were replaced with alanine (FLAG-tagged Kir6.2-205-228A). When co-transfected with HA-tagged GK in HEK293T cells, immunoprecipitation revealed that no co-immunoprecipitated Kir6.2-205-228A was detected (F). (G) Mutant Kir6.2-205-228A impairs high glucose-induced inhibition of K_ATP_ without affecting K_ATP_ activity in low glucose. Inside-out patch clamping was performed as in Figure 4F, except that Kir6.2-205-228A was expressed in HEK293T cells for the preparation of membrane patches. The channel open probability was quantified represented as the percentage of total opening time within a 1-min analysis window. Data are mean ± SEM; n = 16 membrane patches for each treatment, with the *p* value determined by unpaired two-sided Student’s *t*-test. (H) Validation of intestine-specific Kir6.2 replacement with its GK-binding-defective mutant in mice. The *Kir6.2*-IKO mice (*Kir6.2*-floxed mice crossed with *Villin*-*CreERT2* mice) were crossed with the mice expressing either the GK-binding-defective, Kir6.2-205-228A mutant, or wildtype Kir6.2 as a control, all driven by the *Rosa26*-*LSL* system. The genotyping and immunoblotting data regarding the expression levels of Kir6.2 in intestinal tissues are shown. (I) MCT1, which transports pyruvate and lactate as substitutes for glucose to generate ATP in low-glucose conditions, is expressed at higher levels in L cells than in pancreatic β cells. STC-1 cells and β-TC-6 cells were lysed, followed by immunoblotting to assess MCT1 expression. (J) Prior binding of ATP to K_ATP_ is still essential for glucose-induced GLP-1 secretion. *GK*-knockdown STC-1 cells infected with a lentivirus carrying GK-L309P were infected with lentivirus carrying the ATP-binding-defective Kir6.2-K185E/R50G/R201G or wildtype Kir6.2 as a control. Cells were then treated with indicated concentrations of glucose for 1 h, followed by determination of GLP-1 secretion. Data are shown as mean ± SEM; n = 8 biological replicates for each treatment, and *p* value by one-way ANOVA, followed by Tukey. Experiments in this figure were performed three times, except D four times and F five times.

