## Supplementary material for "Glucose is a ligand-like trigger for secretion of postprandial GLP-1": Full scan of western blot

Fig. 5A

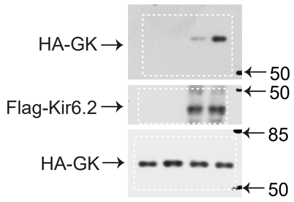


Fig. 5B

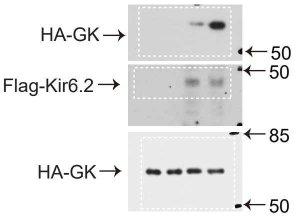


Fig. 5C

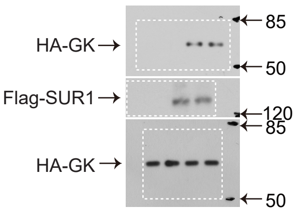


Fig. 5D

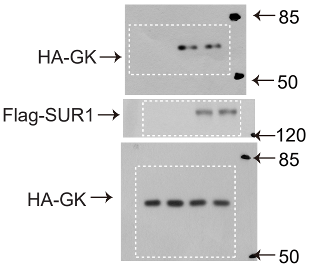


Fig. 5E

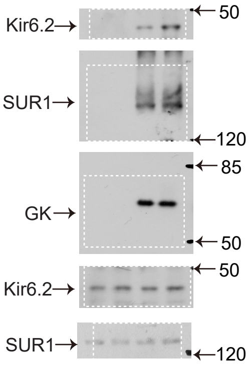


Fig. 5F

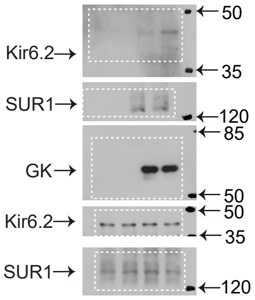


Fig. 5G

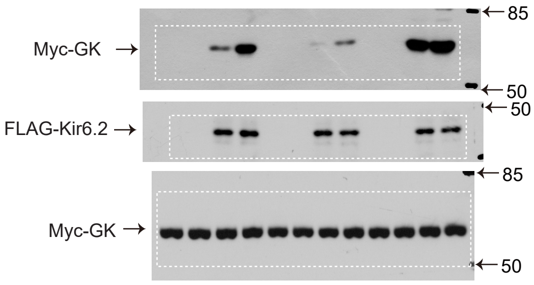


Fig. 5H

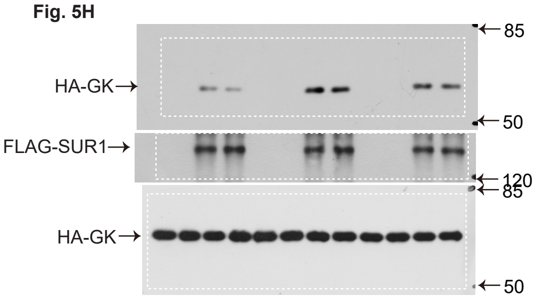


Fig. 5J

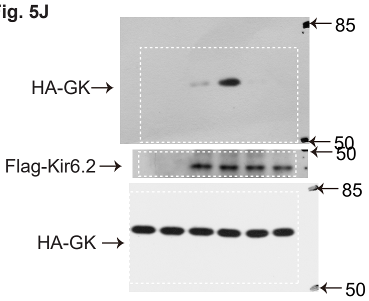


Fig. S2B

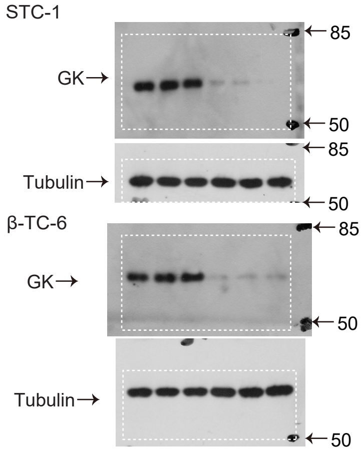


Fig. S2C

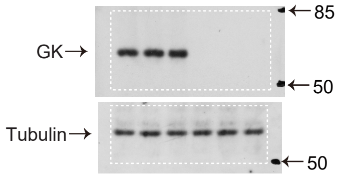


Fig. S2G

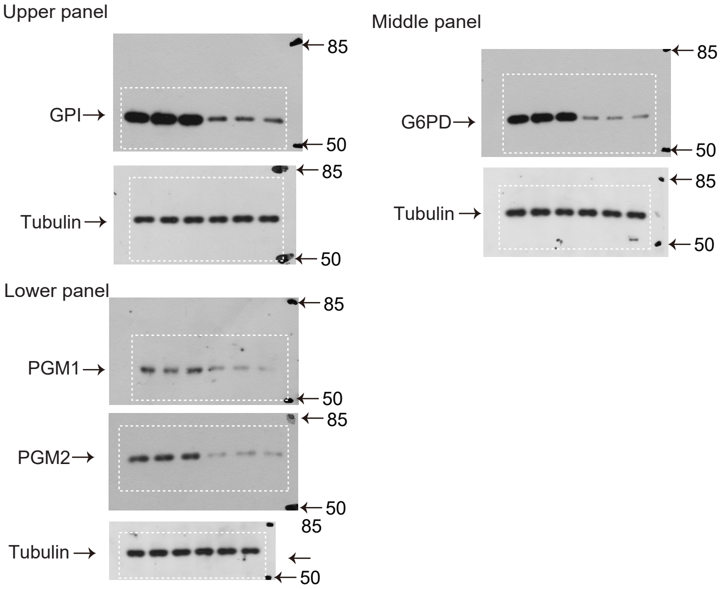


Fig. S2H

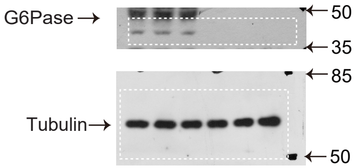

**Fig. S4D**

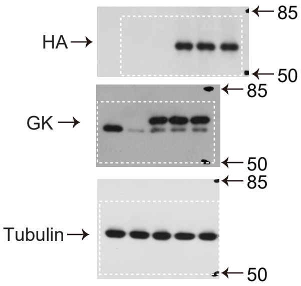


**Fig. S4H**

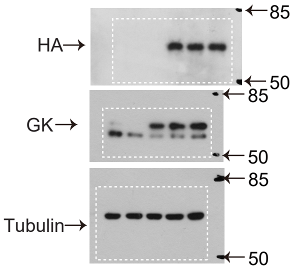


**Fig. S4L**

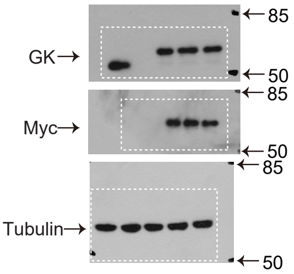


**Fig. S6F**

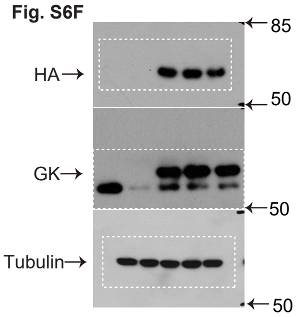


**Fig. S5E**

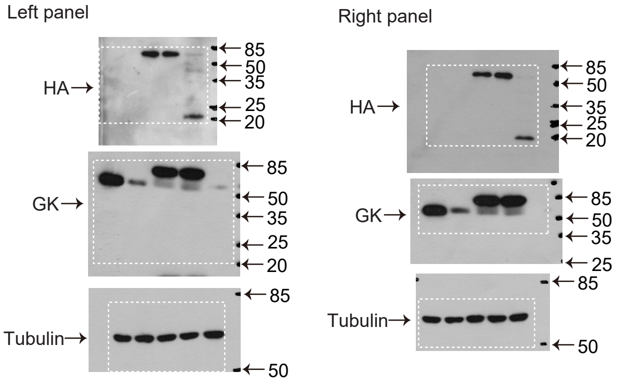


**Fig. S7A**

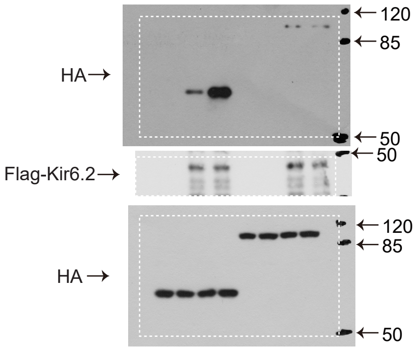


**Fig. S7B**

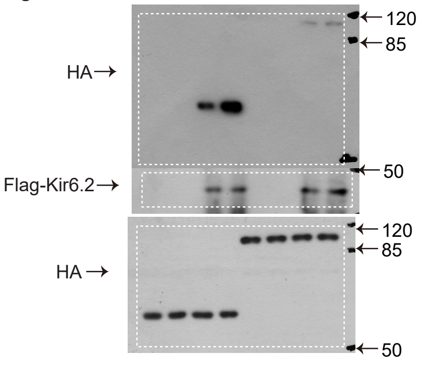


**Fig. S7C**

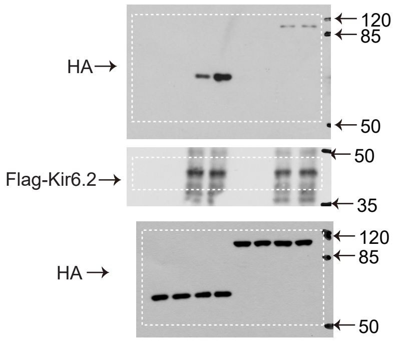


**Fig. S7D**

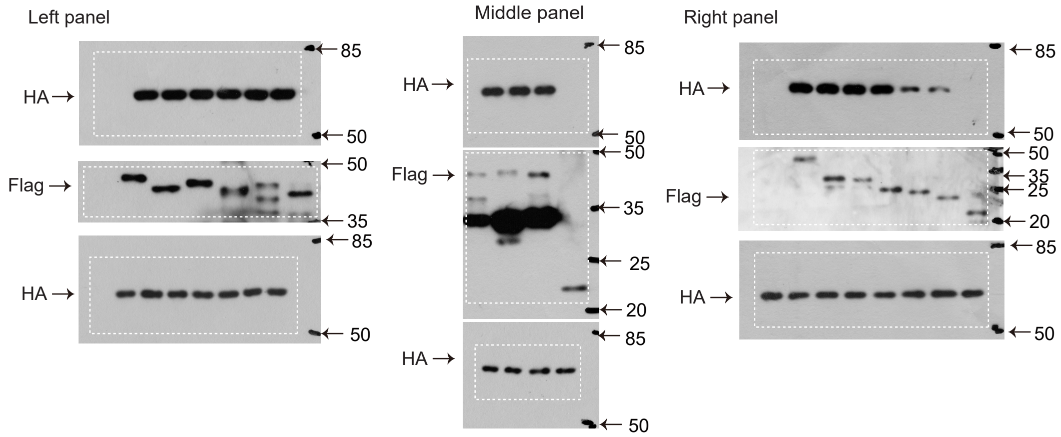


**Fig. S7F**

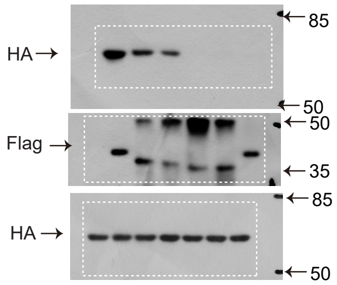


**Fig. S7H**

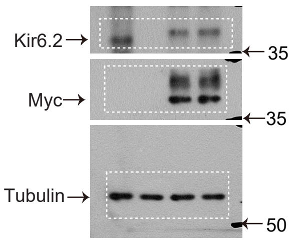


**Fig. S7I**

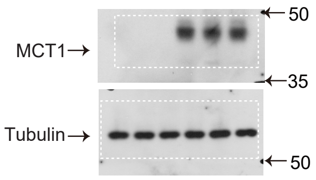
